# A Two-Step Synthesis of Covalent Genetically-Encoded Libraries of Peptide-Derived Macrocycles (cGELs) enables use of electrophiles with diverse reactivity

**DOI:** 10.1101/2025.08.25.672157

**Authors:** James H. Walker, Arunika I. Ekanayake, Nichole Pedowitz, Ryan Qiu, Peter Girnt, Brett M. Babin, Alexey Atrazhev, Lela Vukovic, Matthew M. Bogyo, Ratmir Derda

## Abstract

Genetically-encoded libraries of peptide-derived macrocycles containing electrophile ‘warheads’ (cGELs) can be used to identify potent and selective covalent ligands for protein targets. Such cGELs are synthesized either by incorporation of unnatural amino acids that display mild electrophiles on their side chains or by chemical post-translational modification (cPTM) of mRNA or phage-displayed peptide libraries. Here we investigate fundamental barriers to the synthesis of cGELs. We observe that a previously reported cPTM that proceeds in neutral-to-basic conditions creates mixtures of regioisomers. The complexity of the resulting mixture scales with the electrophilicity of the warhead used in the linker, with some electrophiles being not suitable for use under basic conditions. In contrast, use of a Knorr-pyrazole cPTM enables attachment of electrophiles in acidic pH, thus preventing unwanted reactions with nucleophilic sidechains. The Electrophile is activated only upon mixture with the desired protein target in neutral pH. We use this approach to generate a cGEL with alkyne-bearing macrocycles and use it to identify covalent macrocyclic ligands for pyruvate kinase 2 (PKM2). Our results suggest that construction of cGELs should be performed in conditions that silence the electrophiles (e.g., acidic environment) to prevent unwanted side reactions. In addition to the Knorr-pyrazole method, many other biocompatible bond-forming processes that proceed in mildly acidic pH are likely to be equally effective in constructing cGELs.

## Introduction

Peptide-based therapeutics are a rapidly growing class of drug candidates that bridge the gap between small molecules and biologics.^1, 2^ Their size (typically 500–2000 Da) and structure make them well suited for targeting large, flat, or shallow protein interfaces, such as those found in protein–protein interactions (PPIs), which are often considered “undruggable” by conventional small molecules.^3–8^ Macrocyclization is a key strategy used to overcome the limitations of linear peptides, including poor metabolic stability and low cell permeability. Cyclization helps pre-organize peptide conformations, increases resistance to proteolysis, and can improve pharmacokinetics, in some cases enabling oral bioavailability.^1, 5, 9–18^ Genetically encoded library (GEL) technologies including phage, mRNA, yeast, and DNA-encoded libraries have accelerated discoveries of macrocyclic peptides. These platforms enable screening of highly diverse libraries (10^8^–10^13^ members), linking the structure of each peptide to a genetic tag, enabling amplification and identification of high-affinity binders.^7, 8, 14, 18–28^ GEL technologies have been used to identify many macrocyclic ligands, including clinical candidates, and have proven especially effective for targeting PPIs.^5, 29–32^

To expand the chemical diversity of GELs, strategies that incorporate non-natural amino acids (UAAs) and chemical post-translational modifications (cPTMs) have become increasingly common.^9, 18, 28, 33–36^ Late-stage diversification of libraries by cPTM draws inspiration from Nature’s approach to post-translational diversification of proteins. Modification of library members after translation of the libraries but before selection conveniently expands the accessible chemical space beyond 20 canonical amino acids and natural topologies without alteration of the underlining genetic code.^37–41^ Such expansion is especially useful for introduction of arbitrary unnatural fragments that might not be tolerated by ribosomal translation^9^. Such fragments include reactive electrophilic functionalities poised to form covalent bonds with biological nucleophiles.^28, 42^ Installing electrophilic warheads enables direct screening of libraries designed to form covalent bonds with target proteins.^7, 8, 15, 18, 23, 43–45^ The ability to directly screen libraries bearing reactive electrophiles has opened new paths for identifying potent and selective covalent ligands for a range of targets.^7^

New strategies are needed to enable the installation of reactive (“hot”) electrophilic warheads onto peptide libraries. Most current strategies introduce the electrophile under basic conditions (pH > 8) to drive the reaction.^7, 16, 18, 21, 22, 38, 39, 46, 47^ However, these conditions often activate the electrophile toward unwanted secondary reactions. We hypothesized that basic conditions increase the risk of non-selective reactions between grafted electrophiles and nucleophilic side chains on a protein (e.g. lysine, histidine, tyrosine, cysteine, or the N-terminus; Figure 1a).^7, 8, 35, 45^ These off-target reactions can neutralize the warhead, reduce the concentration of functional binders, and lead to false negatives in the selection (Figure 1a).^22, 48^ The same reactions that install the warhead may reduce its ability to covalently engage with target proteins. To resolve these current limitations, we investigated a cPTM that grafts electrophiles in conditions that silence their reactivity (Figure 1e). Specifically, we repurpose a previously reported Knorr-Pyrazole reaction^39^ to construct macrocyclic GE libraries containing an alkyne electrophile that is otherwise unsuitable for coupling by standard basic conditions (Figure 1b).^48^

**Figure 1.**
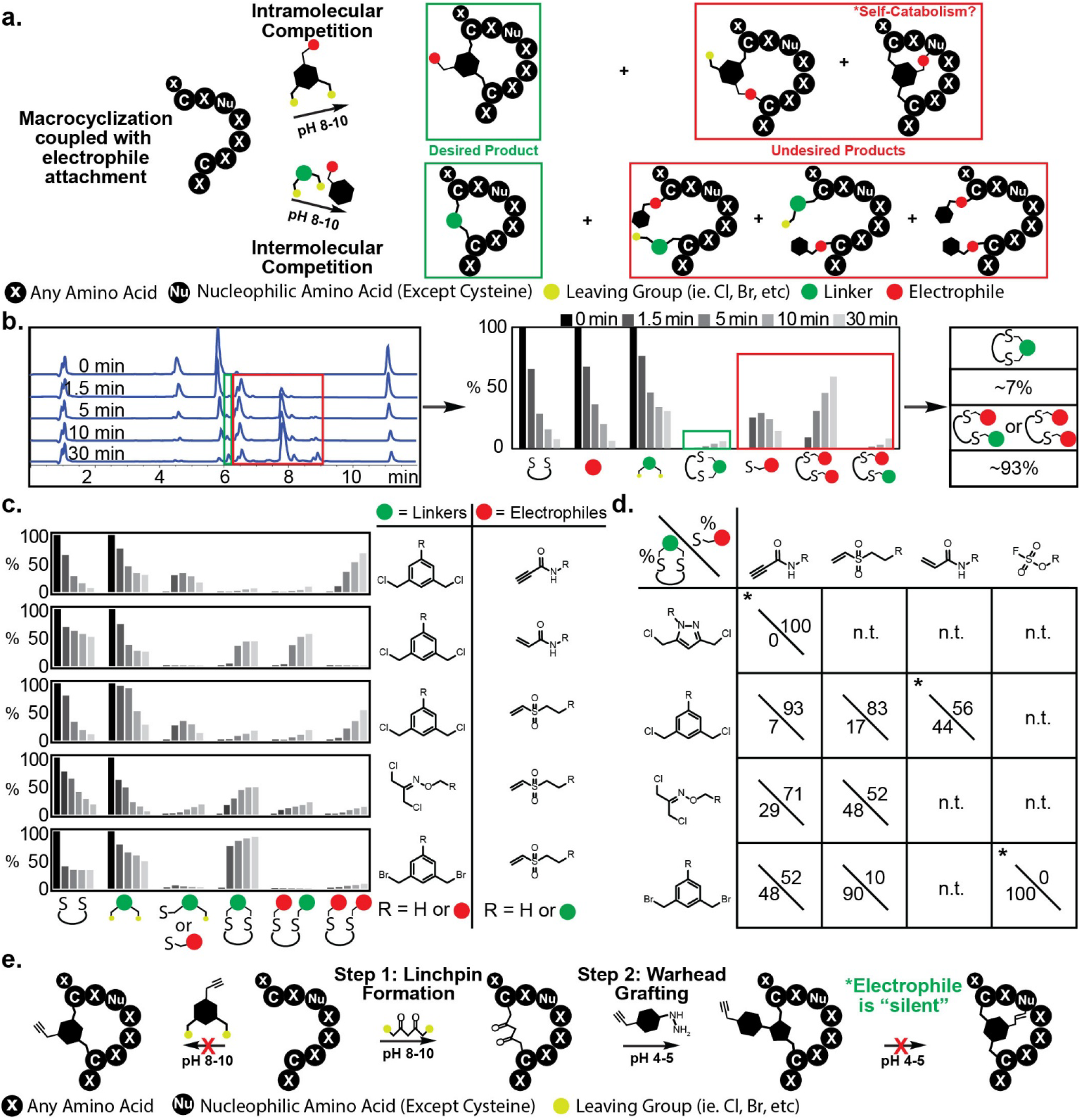
Installation of electrophiles using one step vs. two steps approaches. a) Attaching electrophiles in one step with bidentate linkers can lead to quenching of the electrophile by unwanted side reactions with other nucleophiles (top). Competition reactions between separate electrophilic warheads and linker compounds further illustrate the quenching of the electrophile (bottom). b) Reactions monitored by HPLC can be converted into bar graphs that show the consumption of the peptide, linker, and electrophile starting material to product a percent of product that reacted with either the linker or the electrophile. c) A comparison between dichloroacetone-oxime (DCA-Oxime), dichloro, and dibromo m-xylene (MCX and MBX) linkers and “hot” electrophiles. d) Percent of the undesired and desired product for different warheads and linkers (* intramolecular reactivity monitored using electrophilic warheads with bidentate linkers) (n.t. = not tested). e) Warhead/linker installation in one step vs. in 2 steps with the DKL and Knorr-Pyrazole cyclocondensation.

## Results and Discussion

### 1-step vs. 2-step electrophile installation

Several strategies have reduced unwanted reactions between electrophiles and nucleophiles during macrocyclization, including intrinsically less reactive warheads like aryl fluorosulfates^7, 38^, linkers that require N-terminal serine^16^, and sterically hindered electrophiles like dehydroalanine precursors.^22^ However, none of these fully separate the macrocyclization step from warhead installation. Pre-functionalized linkers avoid the reactivity problem during cyclization yet lack the flexibility of late-stage diversification. Some multi-step modification strategies also reduce phage viability, especially those involving long incubations at non-physiological pH.^40^

We first explored reactions (both intra- and intermolecular) between electrophilic warheads (Vinyl sulfone (VS), Propiolamide (PPA), Fluorosulfate(OSF)^7^, and acrylamide (AcA)) and bidentate linkers (Dichloro-m-xylene(MCX), Dichloroacetone-Oxime (DCO), Dibromo-m-xylene (MBX), dicloro-pyrazole (DCPyr)) (Figure 1a) (Figure S1.1–S1.9). Bar graphs generated from HPLC-monitored reactions show the production of the desired product (thiol-linker bond) and the undesired product (thiol-warhead bond) (Figure 1b). Using a less electrophilic linker, such as MCX or DCPyr, with a reactive electrophile, such as PPA, the reaction between the thiols and the warhead increased substantially (93% and 100% respectively; Figure 1c and 1d) compared with other linker– warhead combinations. However, by either decreasing the electrophilicity of the warhead (to an AcA or VS electrophile) or increasing the electrophilicity of the linker (DCA-Oxime or MBX), the ratio of the desired product to side reactions increases (17%/83% and 44%/56% or 48%/52% and 89%/11% respectively; Figure 1c and 1d). These results suggest that one-step installation of the linker and electrophile is unlikely to be feasible for many commonly used electrophiles (Figure 1d). Although minimal reactivity with the warhead can be achieved with MBX and terminal VS warheads (11%), it is not eliminated completely (Figure 1d) To overcome these limitations, we developed a chemical strategy for phage-displayed peptide libraries that decouples macrocyclization from electrophile installation using orthogonal reaction conditions. The process involves two steps. First, we perform macrocyclization under basic conditions (pH ∼8-8.5) using a diketone-containing linker with a 1,5-dichloropentane-2,4-dione (DPD), to generate a stable cyclic peptide scaffold on the phage-displayed peptide (Figure 1e).^39^ This reaction forms 1,3-diketone-bearing macrocyclic peptide (DKMP) libraries that serve as shelf-stable intermediates. In the second step, we install a hydrazine-containing electrophilic warhead onto the diketone via Knorr-pyrazole cyclocondensation under acidic conditions (pH 4.5–5.0; Figure 1e).^39^ At this pH, the reactivity of the warhead toward peptide nucleophiles is greatly reduced, minimizing off-target modification and preserving its ability to covalently bind to the intended protein target. The reaction proceeds chemoselectively, maintaining electrophile integrity during installation.

### 2-Step installation of hydrazinyl electrophilic warheads

We applied the two-step macrocyclization and electrophile installation strategy to discover covalent peptide inhibitors targeting pyruvate kinase M2 (PKM2), a key metabolic enzyme linked to cancer metabolism, inflammation, and cell proliferation.^48^ PKM2 plays multiple roles in cancer signaling, making it a therapeutically relevant and actively pursued target.^2^ Following macrocyclization, we functionalized the diketone-modified ANCMC peptide libraries with N-(4-hydrazinylphenyl) propiolamide (PPA; Figure S1.10, Scheme S1.2 and S1.3) to introduce the covalent warhead through pyrazole formation (Figure 2a, Figure 3a). We validated each modification by reacting with Biotin-PEG2-hydrazide (BH; Figure 2b) and capturing the functionalized phage with streptavidin-coated magnetic beads (Figure 2c and 2e). Reacting BH with the DPD-modified library (Figure 2c) resulted in 98% capture by the streptavidin-coated beads, indicating near-complete DPD modification (Figure 2d). After installing PPA onto the diketone library, the same biotin capture assay yielded 4% capture (Figure 2f), confirming that 96% of the library was successfully modified with the PPA warhead.

**Figure 2.**
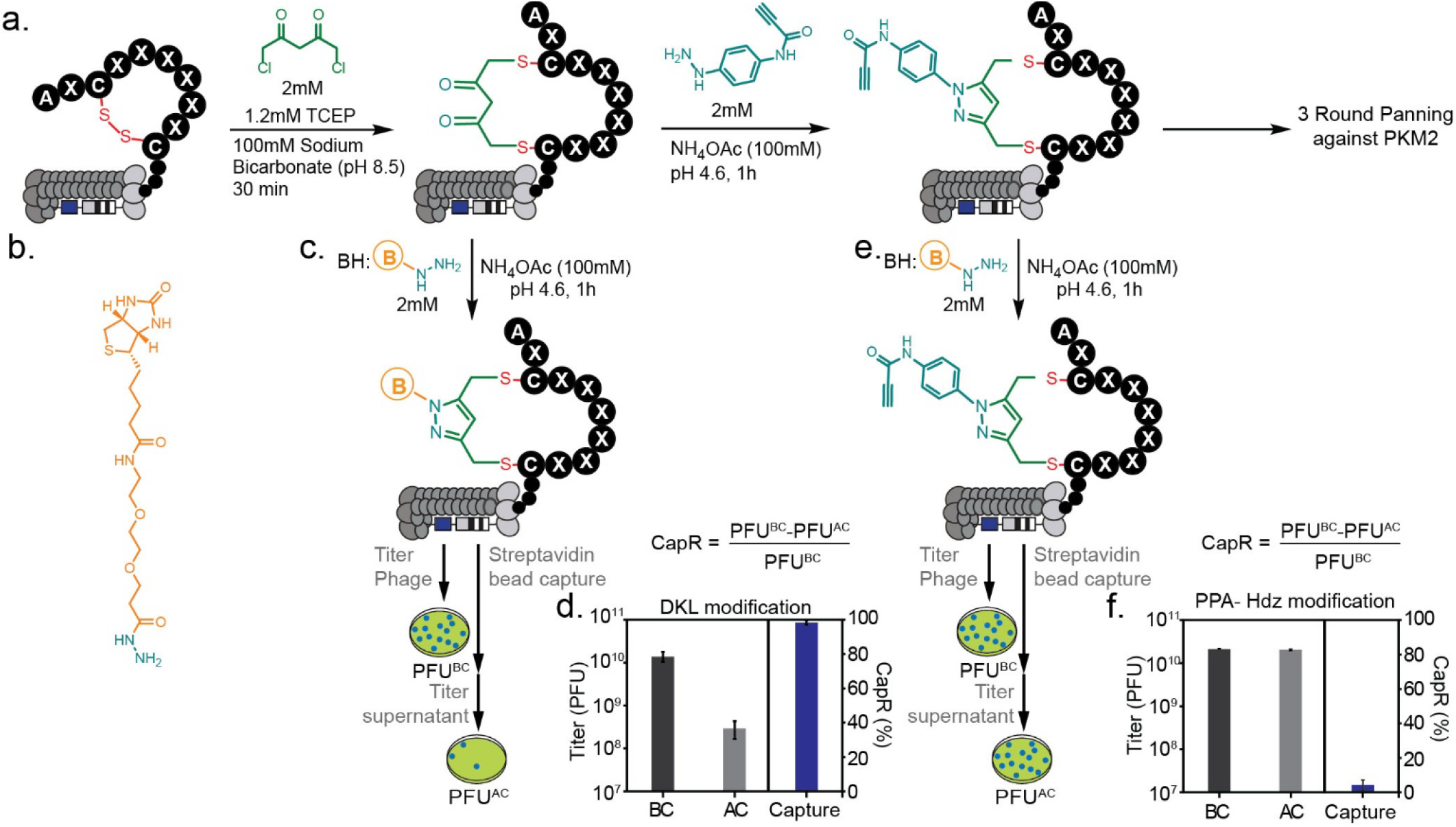
Phage library modification and confirmation by biotin capture. a) ANCMC phage library modified with DPD followed by attachment of the PPA warhead via Knorr–Pyrazole cyclocondensation. The modified library was subjected to three rounds of panning for binding to PKM2. b) Structure of the biotin-PEG2-hydrazide used for biotin capture. c) Confirmation of diketone modification after biotin capture. Phage titers were measured before (BC) and after (AC) capture. d) Biotin capture of DPD-modified library shows 98% capture after reaction with BH, indicating near-complete DPD modification. e) Confirmation of PPA modification via biotin capture was performed by reacting the diketone phage with BH and measuring phage titers before (BC) and after (AC) capture. f) Biotin capture of PPA-modified library shows 4% capture after reaction with BH, indicating near-complete PPA modification.

**Figure 3.**
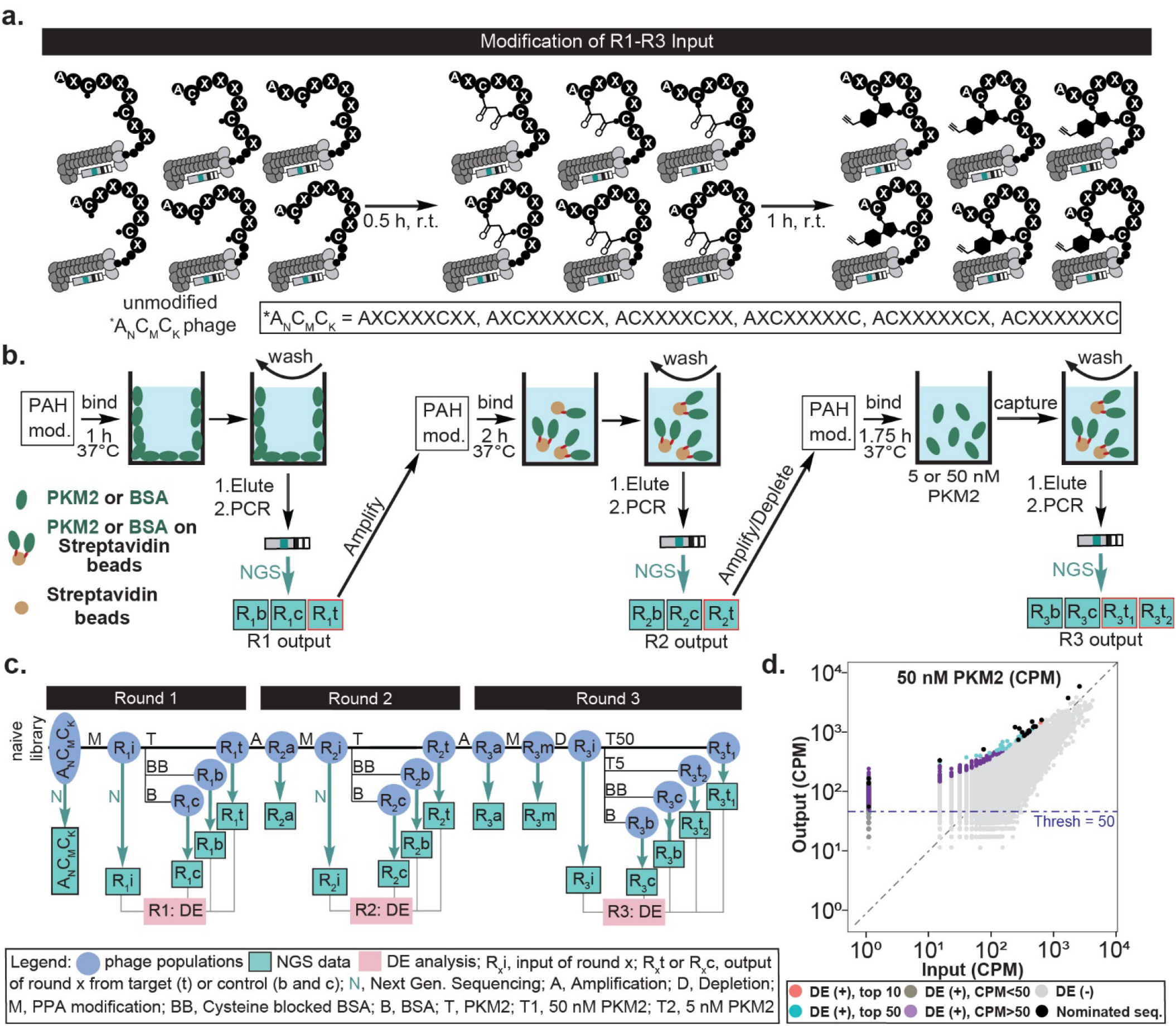
a) Two-step modification of phage library. In the first step, the alkylation of two cysteines installs a linchpin bearing 1,3-diketone. In the second step, Knorr-Pyrazole reaction introduces the covalent warhead. b) Panning procedure for each round (*left*) Round 1: panning against PKM2 coated wells, (*middle*)Round 2: Panning against PKM2 coated streptavidin beads, (*right*) Round 3: Biotin capture of PKM2 and bound phage in solution. c) Panning flow-chart describes the relation between the samples in multi-round panning, their conversion to NGS data and analysis of this data via Differential Enrichment (DE) analysis. d) Scatter plot describes DE analysis of Round 3 of panning that employed 50 nM of PKM2 as a bait. The sequences that passed DE-analysis criteria are highlighted. in purple, blue and black. The selected 25 sequences (black dots) maintained enrichment throughout all three rounds of panning.

### Phage Library Selection and Deep Sequencing Analysis

The starting point of the selection was a mixture of five phage-displayed libraries of disulfide-constrained peptides, each containing six variable positions that contain 19 amino acids encoded by TriNucleotid codons. The libraries denoted AXCX3CX2, AXCX4CX, ACX4CX2, ACX5CX, AXCX5C and ACX6C contain four different ring sizes and combined theoretical diversity of 6 × 19^6^ (2.8 × 10^8^) unique sequences. As in previous reports^9, 49^ these libraries were designed to have a uniform representation of amino acids and X represents 19 natural amino acids, excluding Cys encoded by a 1:1:…:1 mixture of TriNucleotide codons. While macrocycles of large surface area (>12 amino acids) are more likely to yield potent hits^50, 51^, we elected to use moderately sized peptides (9 amino acids) and rings (3-6 amino acids). These cycles offer a balance between sufficient diversity to discover a favorable binding ligand while retaining the potential to engineer cell permeability in selected hits.^50, 52^

After modification of the libraries to introduce the PPA warhead (Figure 3a), we performed three rounds of selection with increasing stringency in each round, using PKM2 as bait (Figure 3b). In R2, we incubated phage with streptavidin beads coated with PKM2 (with BSA or CB-BSA controls) using the amplified PKM2 output from R1 (Figure 3b). Finally, in R3 we mixed the amplified PKM2 R2 output with biotinylated PKM2 (at 5 and 50 nM) or biotinylated CB-BSA in solution. The phage: PKM2 complex was captured with streptavidin beads. The inputs, outputs and control panning samples R1-R3 were analyzed by next-generation sequencing as illustrated in the flow-chart in Figure 3c. The samples eluted from the targets in round n (R_n_t, n = 1, 2) were amplified and modified to yield the input for round (n + 1). Before panning R3, the amplified and modified output from R2 (R_3_m) is depleted to remove any nonspecific binders to BSA controls. Panning the depleted R3 modified phage library against 5 and 50 nM PKM2 enriched 1069 sequences (Figure 3d). From those, we selected 25 sequences for further validation based on their enrichment and overall structural similarities (Figure 3d, 4a; Table S1.1).

### Functional Validation of Selected Peptides through PKM2 Inhibition Assay

From the deep sequencing data, we selected and modified 25 candidate sequences with the DPD linker, followed by the propiolamide (PPA) electrophile (Scheme S1.4 and S1.5). Measurement of inhibitory activity of the 25 macrocycles in PKM2 inhibition assay identified a subset of active sequences spanning four diverse architectures. Measuring the dose-response of these sequences in the same inhibition assay identified six sequences with IC_50_ values that were < 10 µM (Figure 4b, Table S1.2). To test the contribution of the alkyne to IC_50_ values, we modified the selected six sequences with a non-electrophilic propanamide (PEA; Scheme S1.3(bottom)) isostere (Scheme S1.6). Interestingly, the sequences **15d** and **18d** maintained low micromolar inhibitory activity when the covalent PPA warhead was replaced by PEA. In contrast, PPA to PEA replacement in **20d, 25d** and **27d** increased the IC_50_ from 6 µM to 180-580 uM. This decrease in inhibitory activity confirms a critical role for the alkyne in the interactions between macrocycles **20d, 25d** and **27d** and PKM2. However, this observation alone does not confirm formation of a covalent bond. Taken together, these data indicate that the selected peptides interact with PKM2 thought non-covalent interactions with the peptide macrocycle that is enhanced by interactions with alkyne functionality in the PPA warhead. We designed the selection procedure to maximize the probability of discovering macrocycles that covalently react with PKM2. The identification of macrocycles that maintain their inhibitory activity without the alkyne warhead suggest that, even when using probes with electrophilic warheads, it remains possible to select hits based on reversible tight binding such that inhibition does not depend on the warhead.

**Figure 4.**
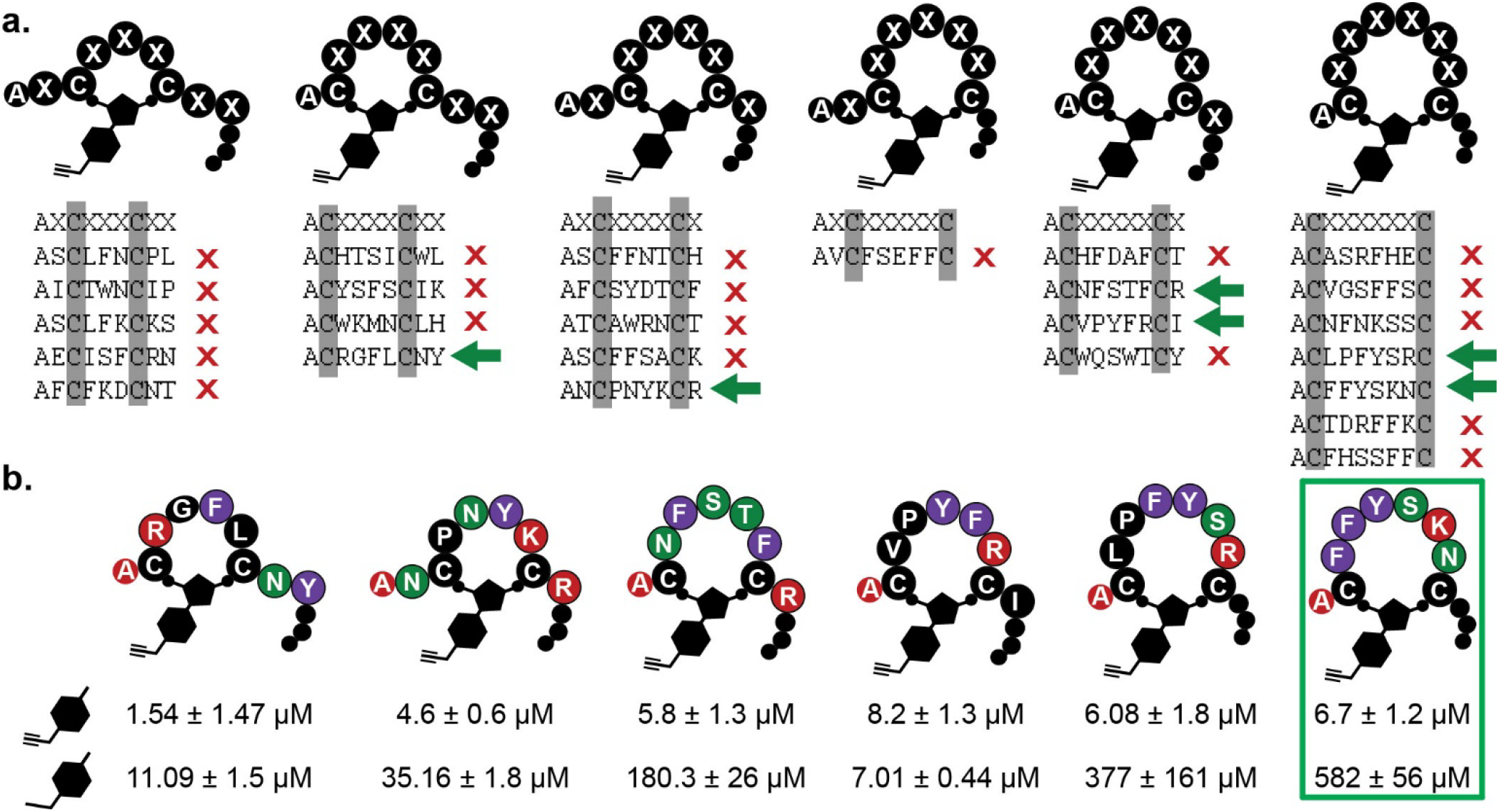
a) Peptide sequences that were selected based on their enrichment in the 3-rounds of panning against PKM2. Sequences identified as binders (green arrow) have an IC_50_ of < 10 µM, non-binding or weak binding sequences (red “X”) have an IC_50_ of > 40 µM. b) IC_50_ of the six selected sequences with the covalent warhead (top) and the control warhead (bottom). The final sequence (green square) was selected for further exploration with LCMS and Docking.

### Characterization of Covalent Adduct Formation by LCMS

Previous reports^7, 38, 53^ use time-dependent increase in inhibition to confirm formation of a covalent bond. However, to directly confirm covalent bond formation, mass spectrometry is required. Therefore, we employed a combination of mass-spectrometry and molecular docking to define the binding mode of our top hit. We focused this analysis on macrocycle **27c** because it was the most sensitive to the presence of warhead in the inhibitory assays. LCMS analysis confirmed covalent modification of PKM2 after incubation with **27c** (Figure 5a). Before incubation, the LCMS data show a single peak corresponding to the monoisotopic mass of 59.942 kD for the PKM2 monomer (Figure 5b, 5c). After incubation, two peaks with slight overlap appear, corresponding to PKM2 and PKM2+ inhibitor (Figure 5d). Analysis of the monoisotopic ion spectrum revealed formation of both 1:1 and 1:2 (PKM2: inhibitor) adducts (61.257 kD and 62.573 kD, respectively), indicating multiple accessible reaction sites on the protein (Figure 5e; Figure S1.18a). We then employed alanine scanning and N-terminal modification to analyze structure– activity relationships. Substitution of the tyrosine residue in **27c** with alanine (**32c**) decreased adduct formation, indicating the importance of this residue for optimal positioning of the covalent warhead (Figure S1.18c). Addition of a GGG(Z) tail (where Z represents propargyl glycine) to generate **30c** similarly reduced adduct formation, likely due to steric hindrance that limits productive binding conformations (Figure S1.18b). The combined modification, **33c**, incorporating both the tyrosine-to-alanine substitution and the GGG(Z) tail, showed improved adduct formation compared to **31c** (Figure S1.18d). Removal of the bulky tyrosine side chain alleviates steric constraints imposed by the C-terminal extension, allowing the peptide to adopt alternative binding conformations conducive to covalent bond formation.

**Figure 5.**
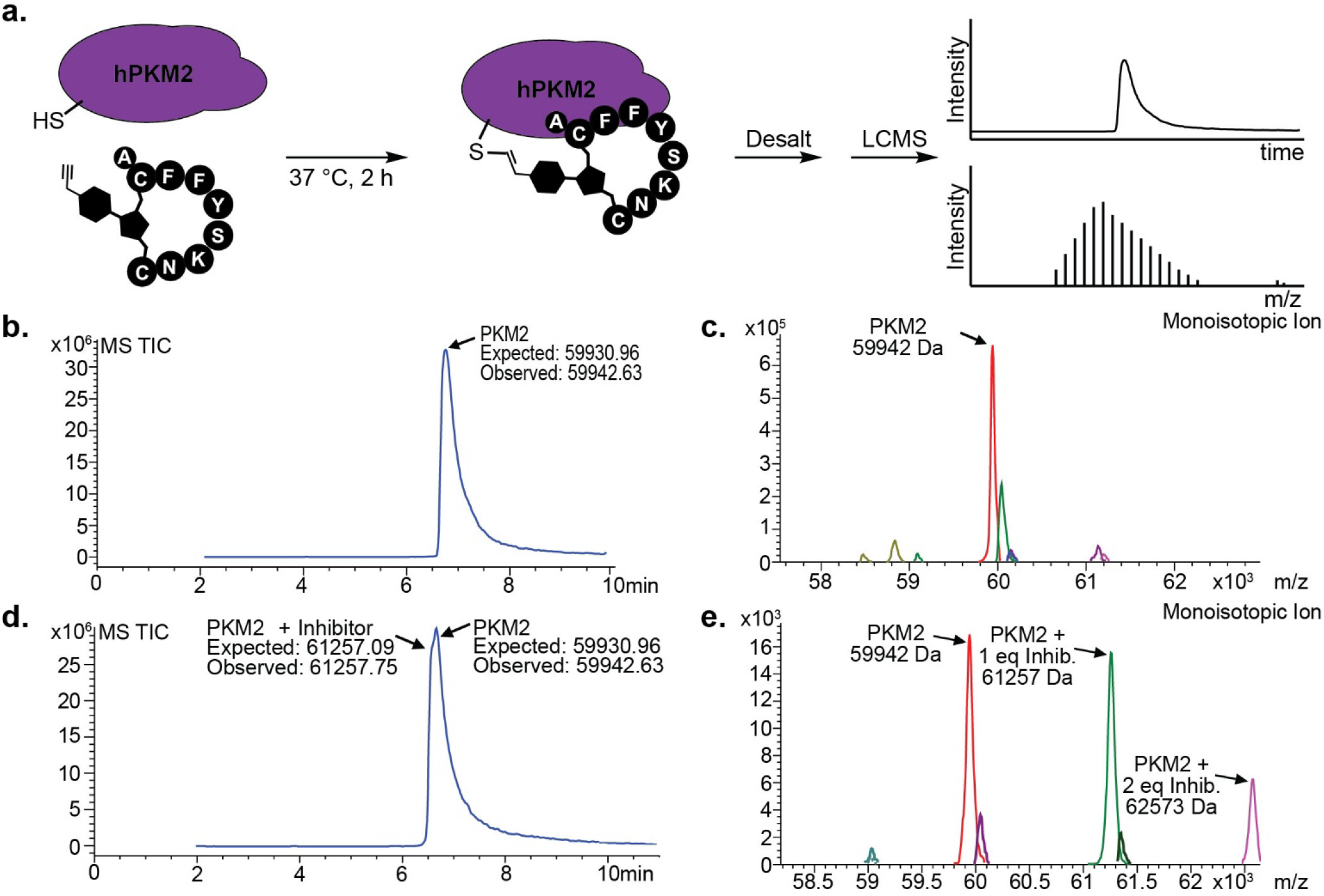
LCMS of PKM2 – **27(c)** adduct. a) PKM2 is incubated with compound **27(c)** for 2 h under physiological conditions (37 °C, pH 7.4) before desalting and analysis with LCMS to obtain an LC trace and Mass spectrum. b) LC trace and monoisotopic ion peak (59,942.63 Da) of PKM2 prior to incubation with **27(c).** c) LC trace and monoisotopic ion peak after incubation with **27(c)** containing PKM2 (59,942.63 Da) and the adduct formed with 1 eq. (61,257 Da) and 2 eq. (62,573 Da) of **27(c)**, suggesting multiple binding sights.

### Molecular Docking Studies Identify Potential Cysteine Targets

Molecular docking with PKM2 monomers and dimers predicted potential binding modes and identified cysteine residues targeted by **27c** (Figure 6a). The analysis identified 10 surface-accessible cysteine residues as potential sites for covalent modification (Figure 6a). Among these, Cys474 and Cys424 emerged as the most probable targets based on favorable docking scores and proximity of the reactive propiolamide warhead to the cysteine thiol group (Figure 6b). Cys474, located near the tetramer interface, displayed particularly strong interactions with docking scores of −6.6 kcal/mol and cysteine–warhead distances of 6.9 Å in the dimer model and 11.9 Å in the monomer model (Figure 6c). Identification of these interface-proximal cysteines aligns with the known allosteric regulation of PKM2 activity through quaternary structure modulation. Covalent modification at these sites could disrupt the equilibrium between PKM2’s less active dimeric and more active tetrameric states, providing a mechanistic explanation for the observed inhibitory effects.

**Figure 6.**
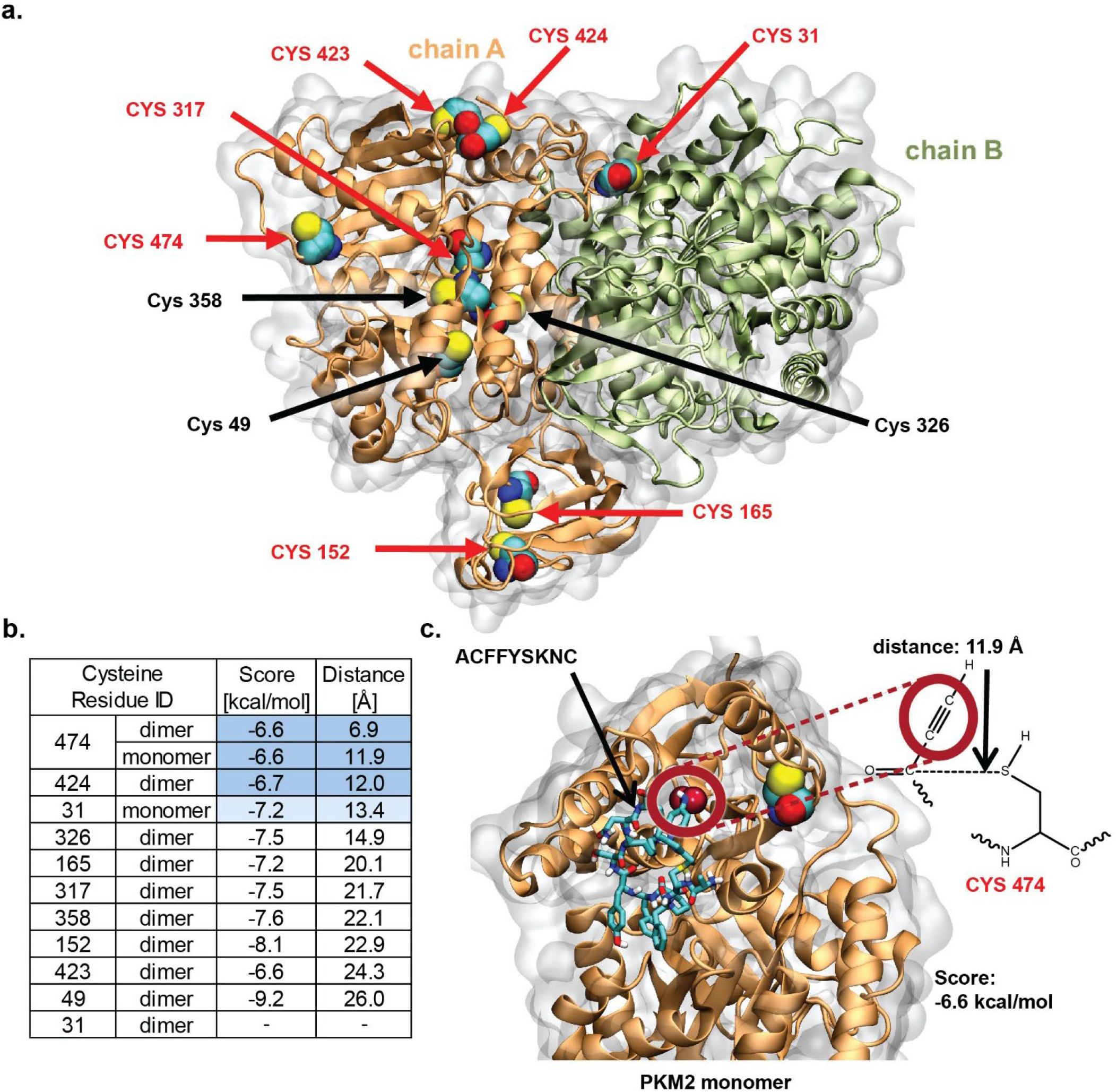
PKM2 Docking study with **27(c).** a) PKM2 dimer model displaying surface Cysteine residues available for binding. b) Table of docking scores and bond length at each cystine residue on the PKM2 monomer and/or dimer. c) **27c** docked to PKM2 monomer showing the docking score (−6.6 kcal/mol) and bond distance (11.9 Å).

## Conclusion

In summary, we introduce a robust and chemo selective strategy for generating genetically encoded libraries of covalent macrocyclic peptides. By decoupling macrocyclization and electrophile installation through pH-orthogonal chemistry, this method addresses a central challenge in display-based covalent inhibitor discovery, namely premature warhead reactivity with peptide nucleophiles. Using orthogonal diketone and Knorr-pyrazole warhead installation (at acidic pH) enables selective, late-stage chemistry to separate macrocyclization (at basic pH) and electrophile functionalization to prevent unwanted reactions with the electrophile. The successful identification and validation of PKM2 covalent inhibitors demonstrate the overall utility of the platform.

This strategy offers a versatile solution for installing electrophilic fragments onto phage-displayed macrocycles and can be extended to a wide range of warheads, affinity tags, and imaging probes, thus expanding the chemical space accessible via genetically encoded libraries.^9, 33, 35, 36, 40–42^ It also supports alternative linker chemistries and functional handles, making it suitable for developing targeted imaging agents or diagnostic tools.^15, 44^ We expect this approach will be broadly applicable for discovering covalent peptide modulators across diverse biological targets using a variety of display platforms.

Applying this methodology to Pyruvate Kinase M2 (PKM2), we identified six lead macrocyclic peptides with IC_50_ values in the low micromolar range (1-10 μM). Notably, our peptides demonstrated varied mechanisms of action, with some (20c, 26c, and 27c) exhibiting strong dependence on the covalent warhead as evidenced by large reductions in potency when the electrophilic was replaced with a non-covalent isostere. Other sequences (15d, 18d, and 21d) maintained significant activity with the non-covalent analog, suggesting a mixed binding mode combining non-covalent interactions with covalent bond formation. LC-MS analysis confirmed covalent adduct formation between our lead peptides and PKM2, while molecular docking studies identified Cys474 and Cys424 as the most probable sites for warhead attachment. These cysteines are strategically located near PKM2’s oligomeric interfaces, suggesting our inhibitors may exert their effects by modulating the equilibrium between the dimeric and tetrameric states of PKM2. Structure-activity relationship analysis through alanine scanning and n-terminal modifications revealed the importance of specific aromatic residues for optimal warhead positioning, while also highlighting the complex interplay between peptide sequence, conformation, and covalent bond formation. The varied binding profiles observed among our lead peptides demonstrate the versatility of our fragment-based discovery approach in accessing diverse inhibitory mechanisms.

Our pH-orthogonal strategy for late-stage diversification of phage-displayed peptide libraries represents a significant advancement in genetically encoded covalent inhibitor discovery. By preserving warhead integrity during installation, this method enhances the efficiency of selection for functional covalent binders. The approach is broadly applicable to various display platforms and compatible with diverse electrophilic warheads, affinity tags, and imaging probes, thus expanding the chemical space accessible through genetically encoded libraries.

## ASSOCIATED CONTENT

### Supporting Information 1

Competition reaction data and protocols(Figure S1.1-S1.9); PKM2 expression protocol; Summary of sequences for selected for assay analysis (Table S1.2); PCR amplification methods (Figure S1.12); panning summary and NGS files of modified libraries (Figure S1.11); NGS DE analysis files (Table S1.1); conditions for DKL and PPA (or PEA) modification; detailed DKL and hydrazinyl-warhead synthetic methods, data processing methods describing the analysis of the DNA sequencing data; Adduct formation monitoring by LCMS method; Activity Assay methods and IC_50_ curves.

### Supporting information 2

Synthesis summaries of peptides listed in Table S1.2 and S1.3 (Figures S2.1-S2.69); H-NMR spectra for compound **3** and **4** (Figure S2.70 and S2.71).

## Notes

The authors declare the following competing financial interest(s): R.D. is the C.S.O. and a shareholder of 48Hour Discovery Inc., the company that licensed US 9958437B2 patent which maintains https://48hd.cloud/ used for deep sequencing analysis. Data Availability: All raw deep-sequencing data is publicly available on https://48hd.cloud/ with data-specific URL listed in Supplementary Figure S1.11 and Table S1.1.

## ACKNOWLEDGMENT

We thank Dr. Randy Whittal and Béla Reiz at the University of Alberta mass spectrometry facility.

