## Supporting information 1 for "A Two-Step Synthesis of Covalent Genetically-Encoded Libraries of Peptide-Derived Macrocycles (cGELs) enables use of electrophiles with diverse reactivity"

**List of abbreviations:**

| | | | |
|---|---|---|---|
| AcA | Acrylamide | MeCN | acetonitrile |
| Boc | *tert*-butyloxycarbonyl | min | minute |
| BSA | Bovine Serum Albumin | MHz | megahertz |
| BSH | biotin-thiol | mL | milliliter(s) |
| CB-BSA | Cysteine blocked BSA | mM | millimolar |
| Da. | dalton(s) | min | minute(s) |
| DCM | dichloromethane | mmol | millimol |
| DCO | Dichloroacetone-Oxime | NGS | Next Generation Sequencing |
| DCPyr | 3,5-Bis(chloromethyl)-pyrazole | OSF | Fluorosulfate |
| DKL | diketone linker | PBS | phosphate buffered saline |
| DMF | *N, N*-Dimethylformamide | PCR | polymerase chain reaction |
| ESI | electrospray ionization | PEA | pyrazole ethanamide |
| equiv. | equivalent(s) | PPA | pyrazole propiolamide |
| EDT | 1,2-ethanedithiol | ppm | parts per million |
| h | hour(s) | r.t. | room temperature |
| HPLC | high-performance liquid chromatography | sec | second(s) (time) |
| | | TCEP | tris(2-carboxyethyl)phosphine) |
| HRMS | high-resolution mass spectrometry | TIS | triisopropylsilane |
| LCMS | liquid chromatography-mass spectrometry | TFA | trifluoroacetic acid |
| | | Tris | tris(hydroxymethyl)aminomethane |
| MBX | α,α’-Dibromo-m-xylene | v/v | volume/volume |
| MCX | α,α’-Dichloro-m-xylene | Z | propargyl glycine |

### Linker-Electrophilic Warhead Competition Reactions

In a vial, a solution of 1-1.5 mM peptide, 100 mM Tris (pH 8.5), 1-1.5 mM linker, and 1-2 mM electrophilic warhead analog was made 20% ACN/$H_2O$. Time points were quenched with 3% TFA prior to HPLC analysis. Linkers used include α,α'-Dichloro-m-xylene (**MCX**), α,α'-Dibromo-m-xylene (**MBX**), Dichloroacetone-Oxime (**DCO**), and 3,5-Bis(chloromethyl)-pyrazole (**DCPyr**). Electrophilic warheads used include (Vinyl sulfone (**VS**), Propiolamide (**PPA**), Fluorosulfate (**OSF**), and Acrylamide (**AcA**). ***DCPyr**, **AcA**, and **OSF** were tested intramolecularly with the linker/electrophilic warhead (Supplementary Figure 1, Supplementary Figure 2, and Supplementary Figure 9). Product and starting material peaks were integrated and percentages calculated for % of starting material area (for each of the starting material percentages) and % of total area of product formed (for product percentages). Each reaction shown in Supplementary Figures 1-9.

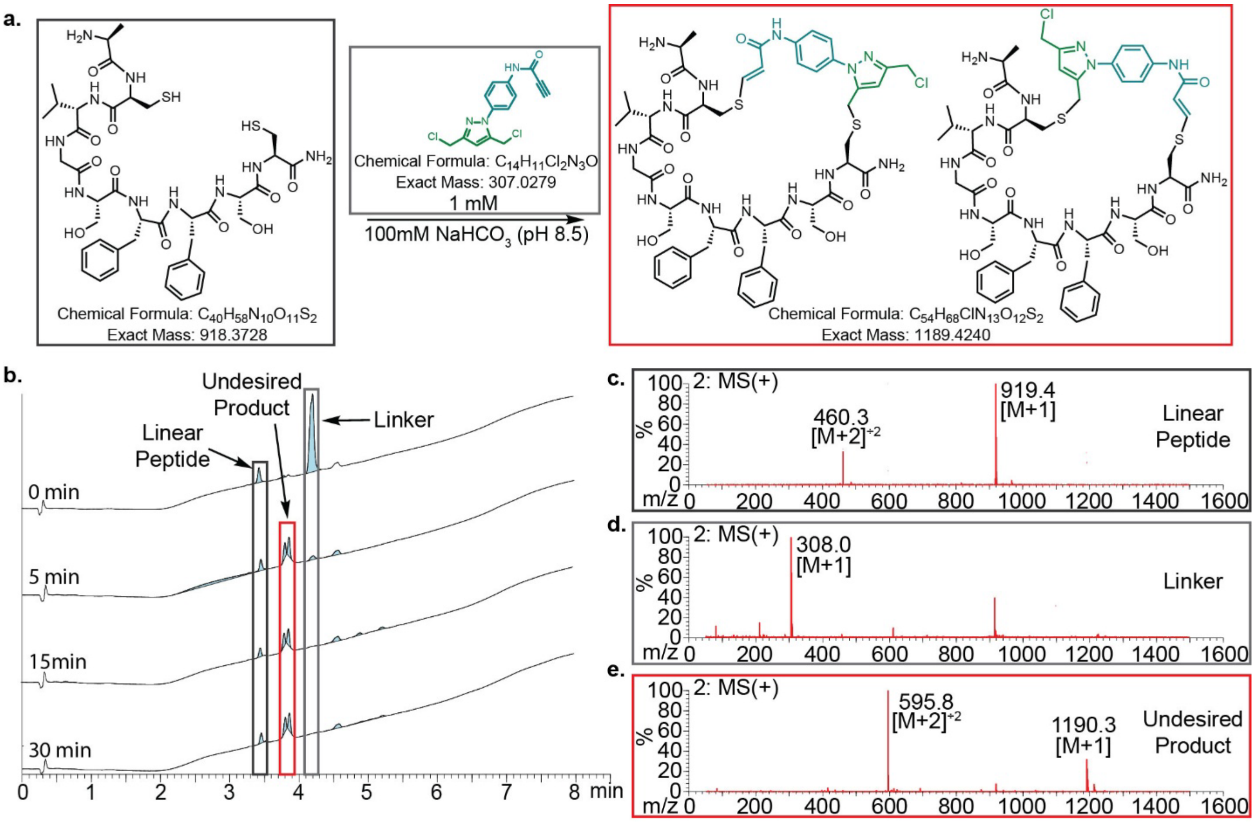

**Figure S1.1. PPA-DKPyr** intramolecular competition.
a) intermolecular competition reaction between **DCPyr-PPA** and ACVGSFFSC. b) LC of the reaction at 0, 5, 15, and 30 min. After 30 min, 100% of the product formed with the peptide is the undesired crosslinked product. c) MS Spectrum of the linear peptide starting material. d) MS spectrum of the **DCPyr-PPA** linker. e) MS spectrum of the crosslinked product (undesired).

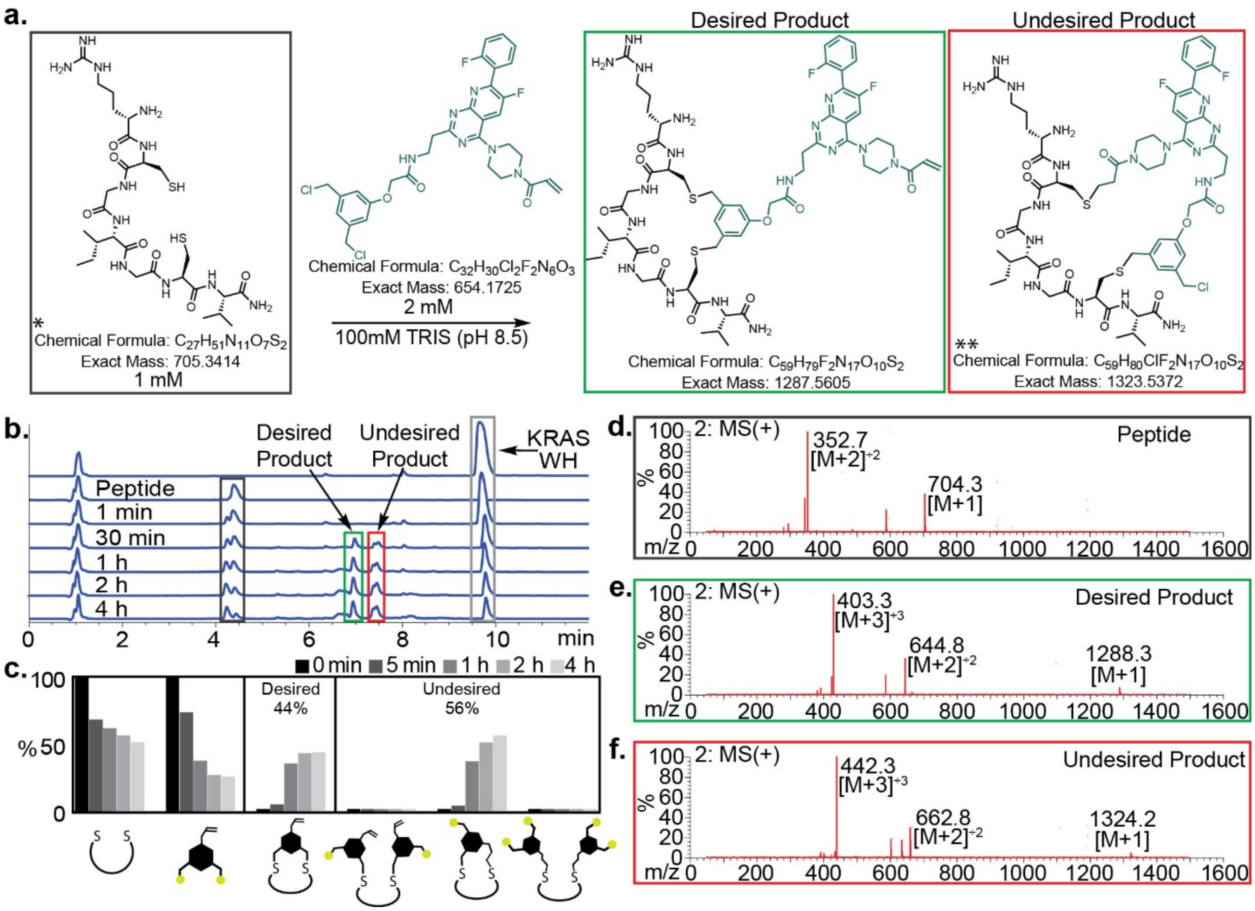

**Figure S1.2. MCX-AcA** intramolecular competition.

a) intramolecular competition reaction between **MCX-AcA** linker (a KRAS inhibitor (Sotorasib) derivative) and RCGIGCV (**30a**). b) HPLC trace of the peptide, linker, and reaction at 0 min, 1 min, 30 min, 1 h, 2h, and 4 h. c) Bar graph showing the consumption of starting materials and production of products over time. After 4 h, the ratio of desired:undesired product is 44%:56%. d) MS Spectrum of the disulfide peptide starting material. e) MS spectrum of the **MCX** linked product (desired). f) MS spectrum of the **MCX-AcA** crosslinked product (undesired).

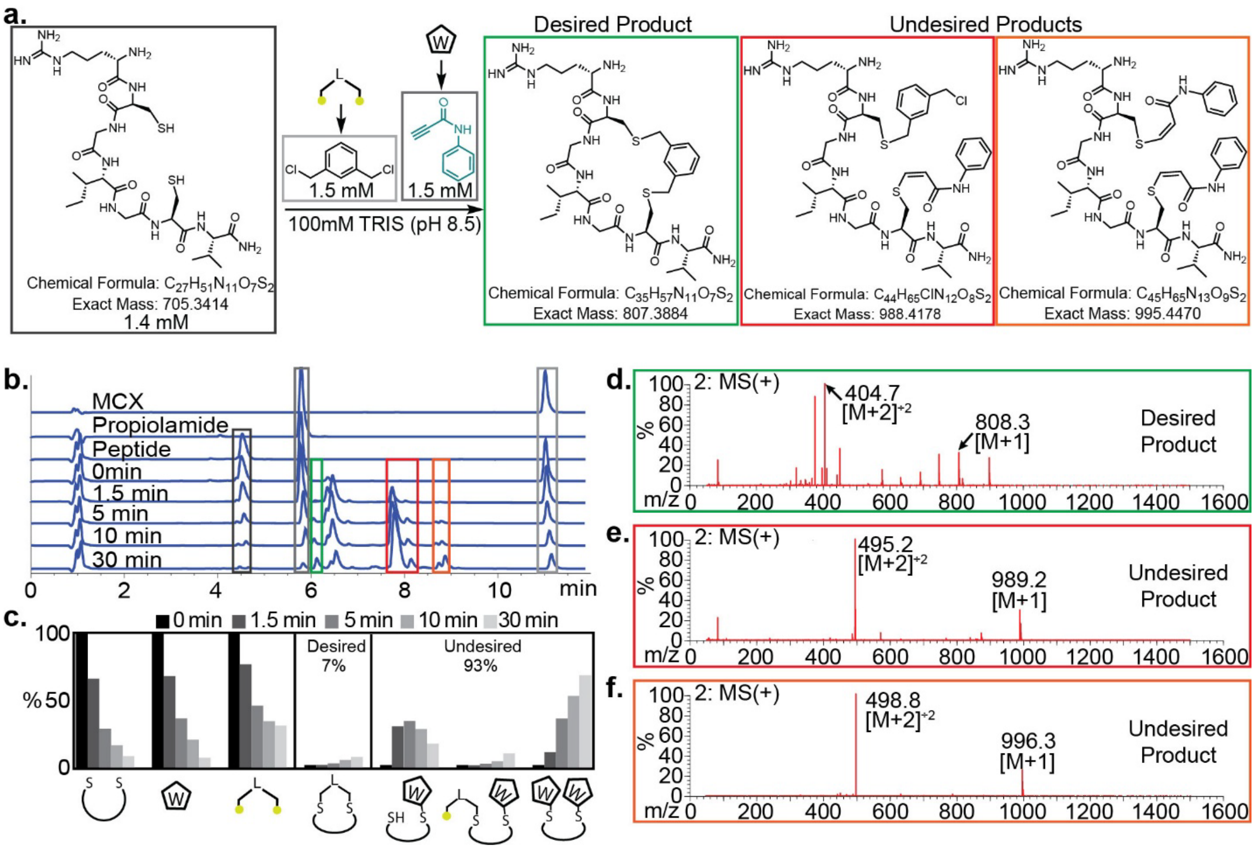

**Figure S1.3.** **MCX-PPA** intermolecular competition.

a) Intermolecular competition reaction between **MCX** linker, **PPA** electrophile and **30a**. b) HPLC trace of the peptide, **MCX**, **PPA**, and reaction at 0 min, 1.5 min, 5 min, 10 min, and 30 min. After 30 min, the ratio of desired: undesired product is 7%: 93%.c) Bar graph showing the consumption of starting materials and production of products over time. d) MS Spectrum of the **MCX** linked product (desired product). e) MS spectrum of the crosslinked **MCX-PPA** product (undesired). f) MS spectrum of the disubstituted-**PPA** product (undesired).

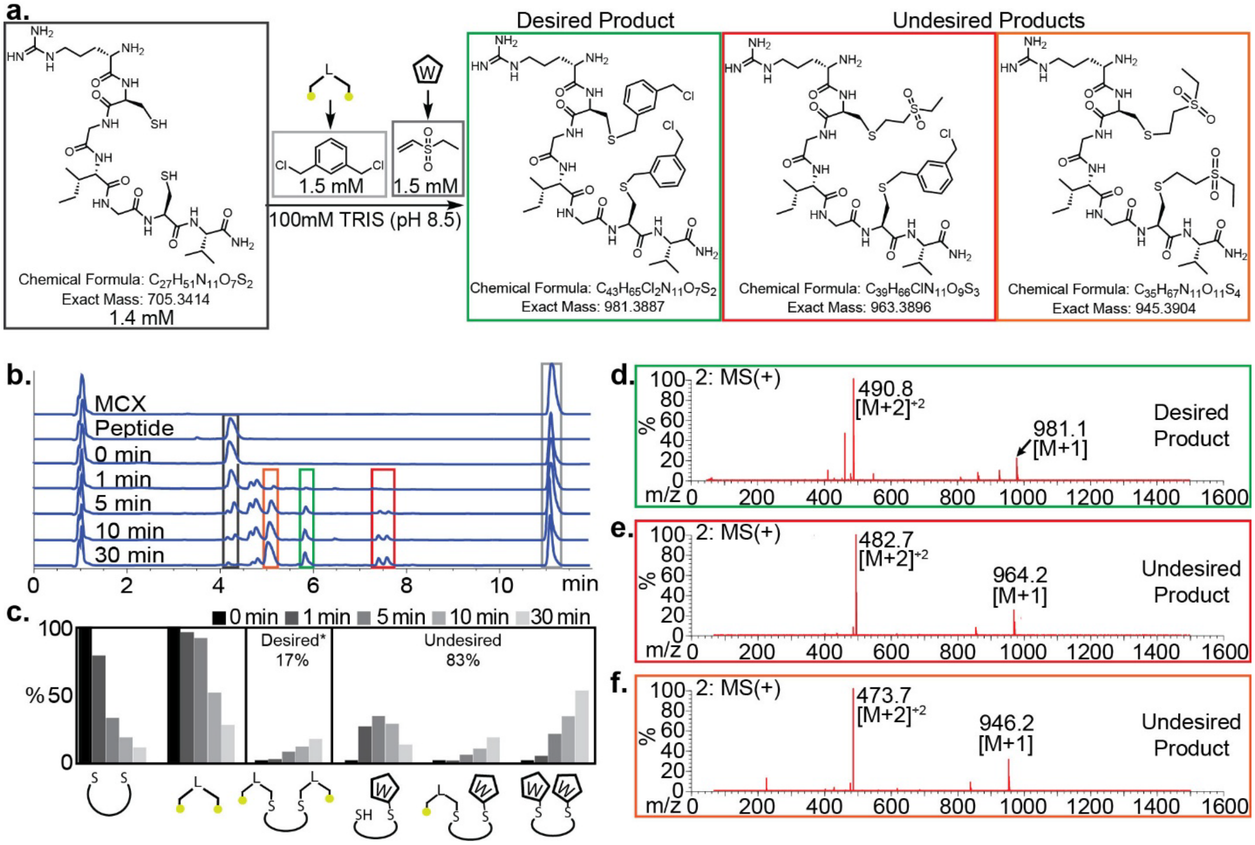

**Figure S1.4. MCX-VS** intermolecular competition.

a) Intrermolecular competition reaction between **MCX** linker and **VS** warhead and **30a**. b) HPLC trace of the peptide, **MCX**, and reaction at 0 min, 1 min, 5 min, 10 min, and 30 min. c) Bar graph showing the consumption of starting materials and production of products and intermediates over time. After 30 min, the ratio of desired: undesired product is 17%: 83%. d) MS Spectrum of the **MCX** linked product (representative of the desired product). e) MS spectrum of the crosslinked **MCX-VS** product (undesired). f) MS spectrum of the disubstituted-**VS** product (undesired).

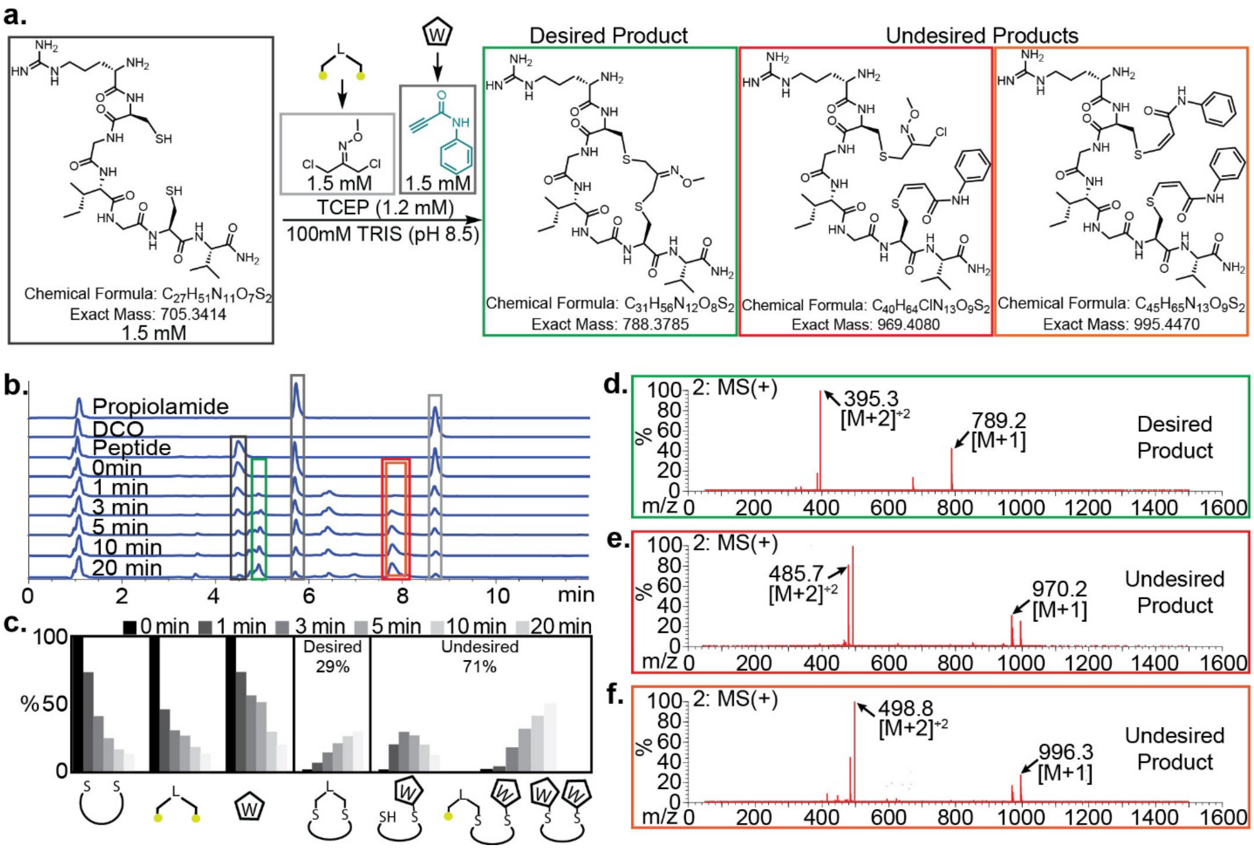

**Figure S1.5. DCO-PPA** intermolecular competition.

a) Intrermolecular competition reaction between **DCO** linker and **PPA** warhead and 30a. b) HPLC trace of the peptide, **DCO**, **PPA**, and reaction at 0 min, 1 min, 3 min, 5 min, 10 min, and 20 min. c) Bar graph showing the consumption of starting materials and production of products and intermediates over time. After 20 min, the ratio of desired: undesired product is 29%: 71%. d) MS Spectrum of the **DCO** linked product (desired product). e) MS spectrum of the crosslinked **DCO-PPA** product (undesired product). f) MS spectrum of the disubstituted- **PPA** product (undesired product).

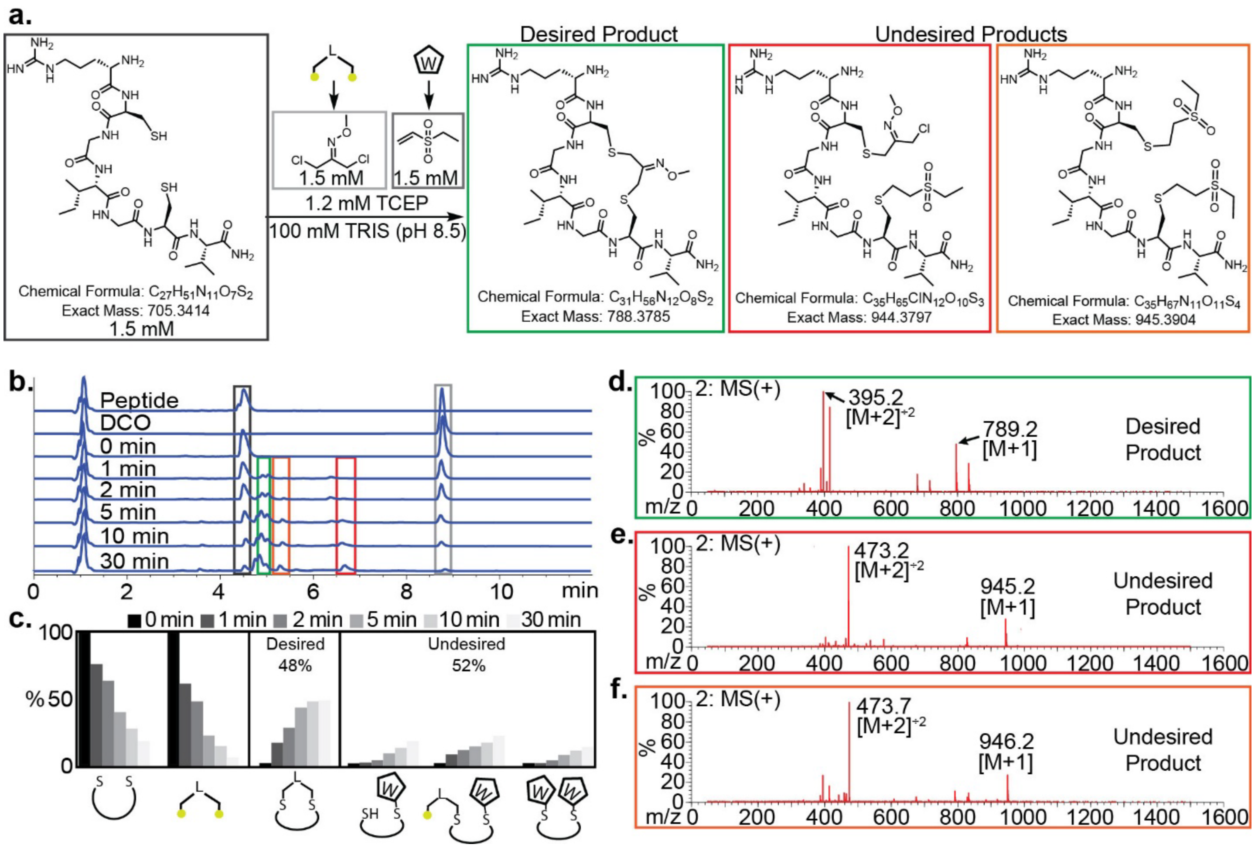

**Figure S1.6. DCO-VS** intermolecular competition.

a) Intrermolecular competition reaction between **DCO** linker and **VS** warhead and 30a. b) HPLC trace of the peptide, **DCO**, and reaction at 0 min, 1 min, 2 min, 5 min, 10 min, and 30 min. c) Bar graph showing the consumption of starting materials and production of products and intermediates over time. After 30 min, the ratio of desired: undesired product is 48%: 52%.d) MS Spectrum of the **DCO** linked product (desired product). e) MS spectrum of the crosslinked **DCO-VS** product (undesired product). f) MS spectrum of the disubstituted-**VS** product (undesired product).

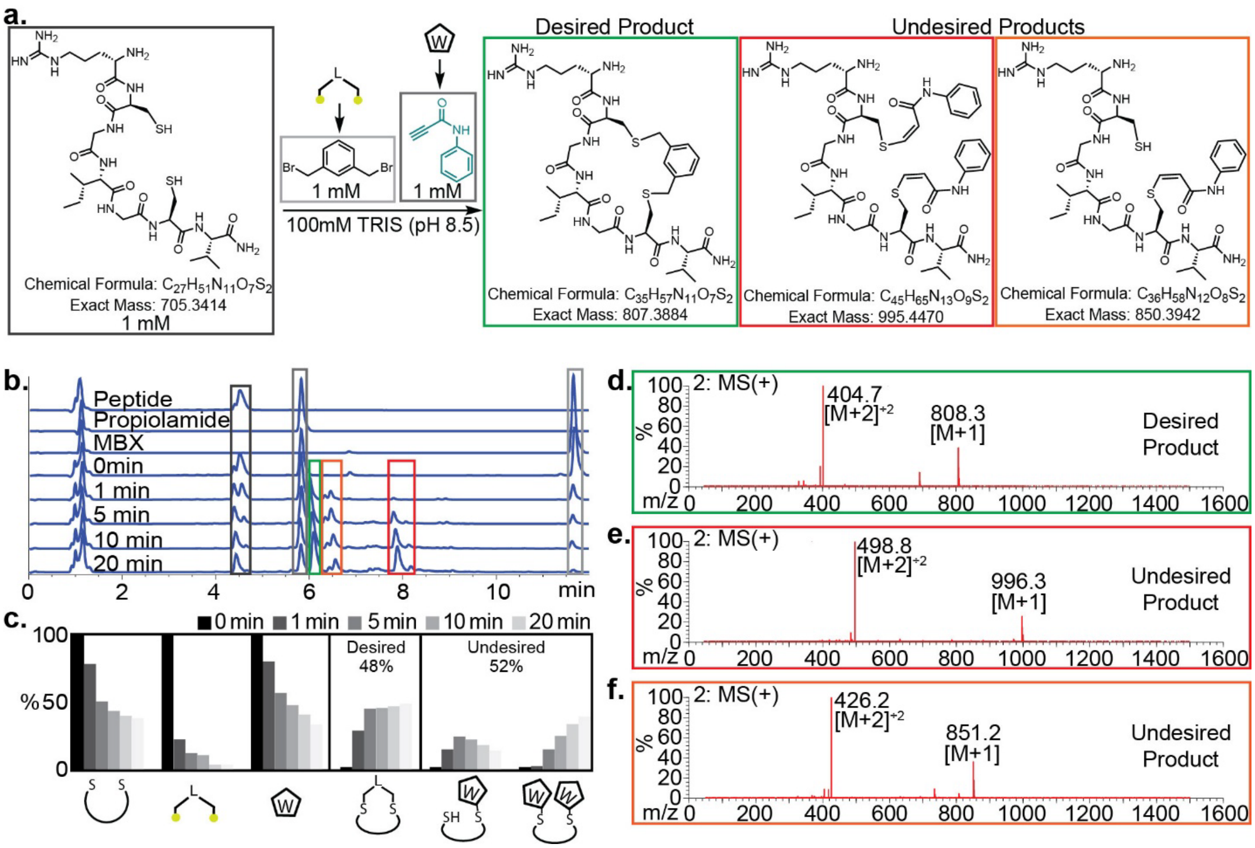

**Figure S1.7.** **MBX-PPA** intermolecular competition.

a) Intrermolecular competition reaction between **MBX** linker and **PPA** warhead and 30a. b) HPLC trace of the peptide, **MBX**, and reaction at 0 min, 1 min, 5 min, 10 min, and 20 min. c) Bar graph showing the consumption of starting materials and production of products and intermediates over time. After 20 min, the ratio of desired: undesired product is 48%: 52%.d) MS Spectrum of the **MBX** linked product (desired product). e) MS spectrum of the diosubstituted-**PPA** product (undesired product). f) MS spectrum of the monosubstituted-**PPA** product (undesired product).

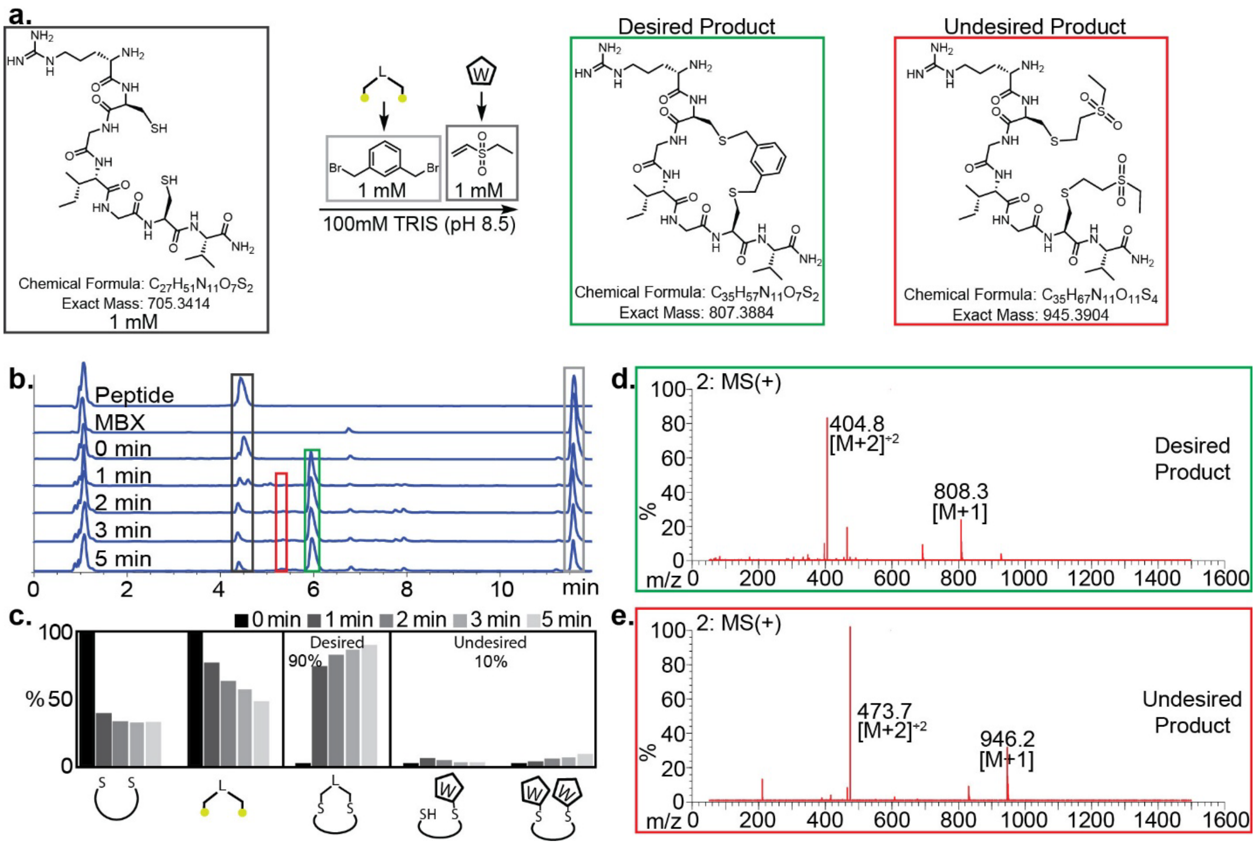

**Figure S1.8. MBX-VS** intermolecular competition.

a) Intrermolecular competition reaction between **MBX** linker and **VS** warhead and 30a. b) HPLC trace of the peptide, **MBX**, and reaction at 0 min, 1 min, 2 min, 3 min, and 4 min. c) Bar graph showing the consumption of starting materials and production of products and intermediates over time. After 20 min, the ratio of desired: undesired product is 90%: 10%.d) MS Spectrum of the **MBX** linked product (desired product). e) MS spectrum of the disubstituted-**VS** product (undesired product).

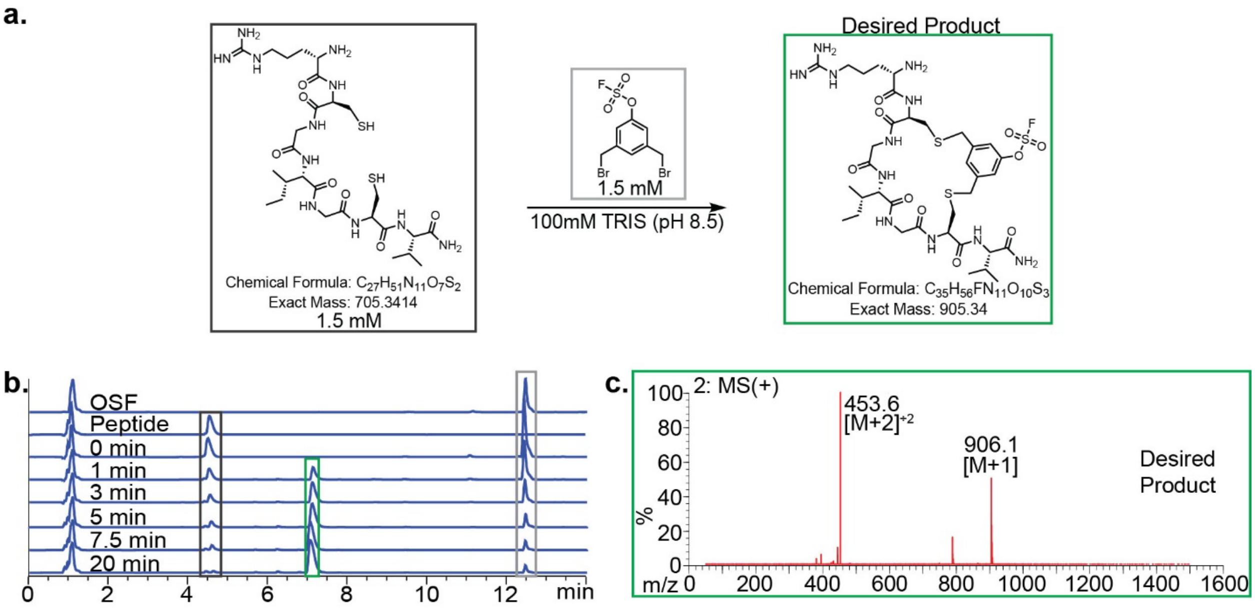

**Figure S1.9. MBX-OSF** Intramolecular Competition

a) Intrramolecular competition reaction between **MBX-OSF** linker and 30a. b) HPLC trace of the peptide, **MBX-OSF**, and reaction at 0 min, 1 min, 3 min, 5 min, 7.5 min, and 20 min. c) MS Spectrum of the **MBX** linked product (desired product product), 100% of the product formed is the desired product.

**PKM2 Expression Protocol**

The pET28a hPKM2 wild-type expression plasmid was obtained from addgene (Plasmid #122690). PKM2 was expressed in BL21(DE3) E. coli cells. A 50 mL starter culture was inoculated from a single colony and grown at 37 °C for 12 hours. The starter culture was diluted 1:40 into 1-L LB media supplemented with 2 mL $MgCl_2$ and grown at 37 °C with shaking (180 rpm) until reaching an $OD_{600}$ of 0.7. Cultures were then moved to room temperature, induced with 0.5 mM IPTG, and shaken at 225 rpm for 6 hours. Cells were harvested by centrifugation (6000 × g, 15 min, 4 °C), resuspended in 60 mL lysis buffer (50 mM Tris-HCl pH 8.5, 10 mM $MgCl_2$, 300 mM NaCl, 5 mM imidazole, 10% glycerol) per liter of culture, and lysed by sonication (30 sec on/30 sec off, 5 min total). Lysates were clarified by centrifugation (20,000 × g, 45 min, 4°C), and the supernatant was supplemented with β-mercaptoethanol (30 μL per 30 mL lysate) before incubation with Ni-NTA beads (Thermo Scientific cat no. R90115) at 4 °C for 2 hours with shaking. Beads were washed three times with wash buffer (50 mM Tris-HCl pH 8.5, 10 mM $MgCl_2$, 300 mM NaCl, 30 mM imidazole, 10% glycerol), and protein was eluted with elution buffer (50 mM Tris-HCl pH 8.5, 10 mM $MgCl_2$, 250 mM NaCl, 250 mM imidazole, 10% glycerol). Eluted fractions were dialyzed twice against 2 L of dialysis buffer (50 mM Tris-HCl pH 7.5, 10 mM $MgCl_2$, 25 mM NaCl, 20% glycerol), overnight and for 4 hours.

**Preparation of PPA-modified ANCMC phage-displayed library**

To remove glycerol from the phage stock: 100 μL of $10^{13}$ PFU/mL of ANCMC phage library in 50% glycerol/Milli-Q water was cleaned 3 times with an equilibrated Zeba spin column. Zeba column was spun at 3000 xG for 3 min each wash and washed with 300 μL of PBS between each phage wash. A solution of approx. $10^{11}$ PFU/mL of ANCMC phage library was then prepared in PBS from the cleaned stock. 88 μL of the prepared solution was transferred to a 1.7 mL centrifuge tube with 1 μL of 200 mM TCEP and 10 μL 100 mM Sodium bicarbonate buffer at pH 8.5 and left to incubate for 30 min at rt. 1 μL of 200 mM **DKL** in DMF was then added to the reaction mixture and incubated for another 30 min at rt. The reaction mixture was then loaded into an equilibrated Zeba column and the flow through was collected by centrifugation at 3000 xG for 3 min. This wash was repeated 2 more times to obtain a clean **DKL** modified phage library. Zeba column was washed 3 times with 300 μL of PBS between each flow through. **DKL** cyclized phage were then modified with biotin hydrazide (BH) (Figure S1.10b) to quantify **DKL** modification through biotin capture and tittering of the captured phage (Figure S1.10c).

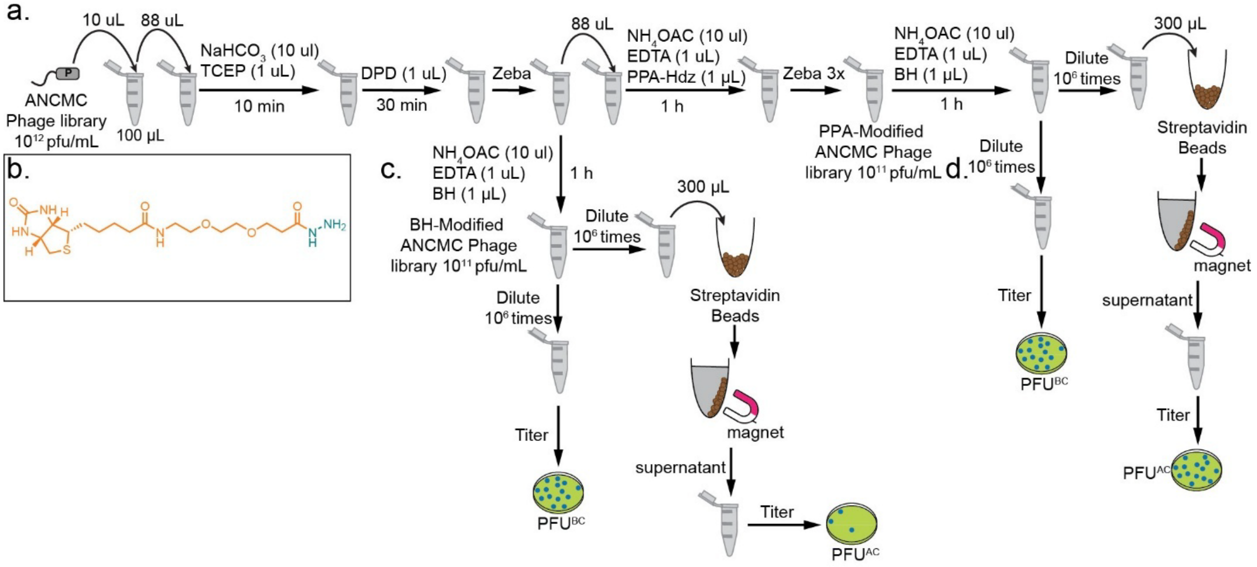

**Figure S1.10.** Confirmation of Modification Through Biotin Capture.
**a**) DKL and Knorr-pyrazole modification of ANCMC library. **b**) Biotin Hydrazide (BH) used to quantify phage modification by biotin capture. **c**) Modification of DKL modified phage with BH followed by biotin capture by streptavidin beads and tittering of captured phage. **d**) Modification of PPA modified phage with BH followed by biotin capture by streptavidin beads and tittering of captured phage

For the **PPA** modification, 88 µL of the **DPD** modified phage library was transferred to a 1.7 mL centrifuge tube with 10 µL 100 mM Ammonium acetate buffer (pH 4.5), 1 µL of 500 mM EDTA (pH 8.0), and 1 µL of 200 mM **PPA** hydrazine in DMF. Reaction was left to incubate for 1 hour at rt. After 1 hour, reaction mix was loaded into an equilibrated Zeba column and the flow through was collected by centrifugation at 3000 xG for 3 min. This wash was repeated 3 times, cleaning the Zeba column with 300 µL of PBS between each flow through. The **PPA** modified phage library was then diluted to 1 mL before each round of panning (Figure S1.10a). PPA functionalized phage were then modified with biotin hydrazide (BH) (Figure S1.10b) to quantify PPA modification through biotin capture and tittering of the captured phage (Figure S1.10d).

**Panning strategy: panning on plate , on beads and in solution**

**Round 1:**

PKM2 was coated on the surface of High Protein Binding 96-well plates (Corning PN:3369) overnight at 4 °C in PBS. In parallel, BSA and Cysteine blocked BSA (CB-BSA) coated wells were coated on wells following the same procedure. Coated wells were then blocked with 2% CB-BSA for 45 min at rt. 100 μL of 3.1 x $10^{10}$ PFU/mL **PPA** modified library was incubated for 2 h at 37 °C. The **PPA** modified library supernatant was then removed and the wells were washed with PBS+0.1% Tween (10 washes, 5 M GdnHCl (1 wash), PBS (10 washes), and then with TEV protease buffer (1 wash) to cleave the phage. With. Eluted phage were then amplified with 2-step PCR using M13 primers and sequenced using Next Generation Sequencing (NGS).

**Round 2:**

A suspension of 50 μL of magnetic streptavidin beads was transferred to a 1.7 mL centrifuge tube and washed 3 times with 1 mL of PBS. The beads were then resuspended in 1 mL of 50 nM biotinylated PKM2 in PBS+2% CB-BSA and incubated for 1 h at rt. In parallel, 50 μL of magnetic streptavidin beads were washed and coated with 2 % biotinylated BSA or CB-BSA. After coating the beads, 100 μL (2.57 x $10^{10}$ PFU/mL) of the **PPA** modified phage library (amplified from round 1 output) was incubated with the coated beads for 1 h at 37 °C. Supernatant was then removed and the beads washed with PBS+0.1% Triton X-100 (5 washes), 5 M GdnHCl (3 washes), PBS (12 washes), 5 M GdnHCl (1 wash) and TEV protease buffer (1 wash) to elute phage. The recovered phage solution was amplified with 2 step PCR and submitted for NGS analysis.

**Round 3:**

For depletion, a suspension of 50 μL of magnetic streptavidin beads was transferred to a 1.7 mL centrifuge tube and washed 3 times with 1 mL of PBS. Beads were then coated with 2% biotinylated CB-BSA in PBS for 1 h at rt. The supernatant was removed and 100 μL of 2 x $10^{10}$ PFU/mL amplified and **PPA** modified library was incubated for 1 h at rt °C. The depleted **PPA** modified library supernatant was then transferred to 1.7 mL centrifuge tubes containing 1 mL of 50 nM biotinylated PKM2 and incubated for 105 min at 37 °C. In parallel, depleted **PPA** modified library supernatants were also incubated with 1 mL of 5 nM biotinylated PKM2 and 1 mL of 2% biotinylated CB-BSA. After incubation, beads were washed with 0.1% Triton X-100 in PBS (1 wash), 5M GdnHCl (3 washes), PBS (6 washes), and then with Milli-Q water (1 wash). Phage was then eluted by boiling, amplified with 2-step PCR using M13 primers, and submitted for NGS analysis.

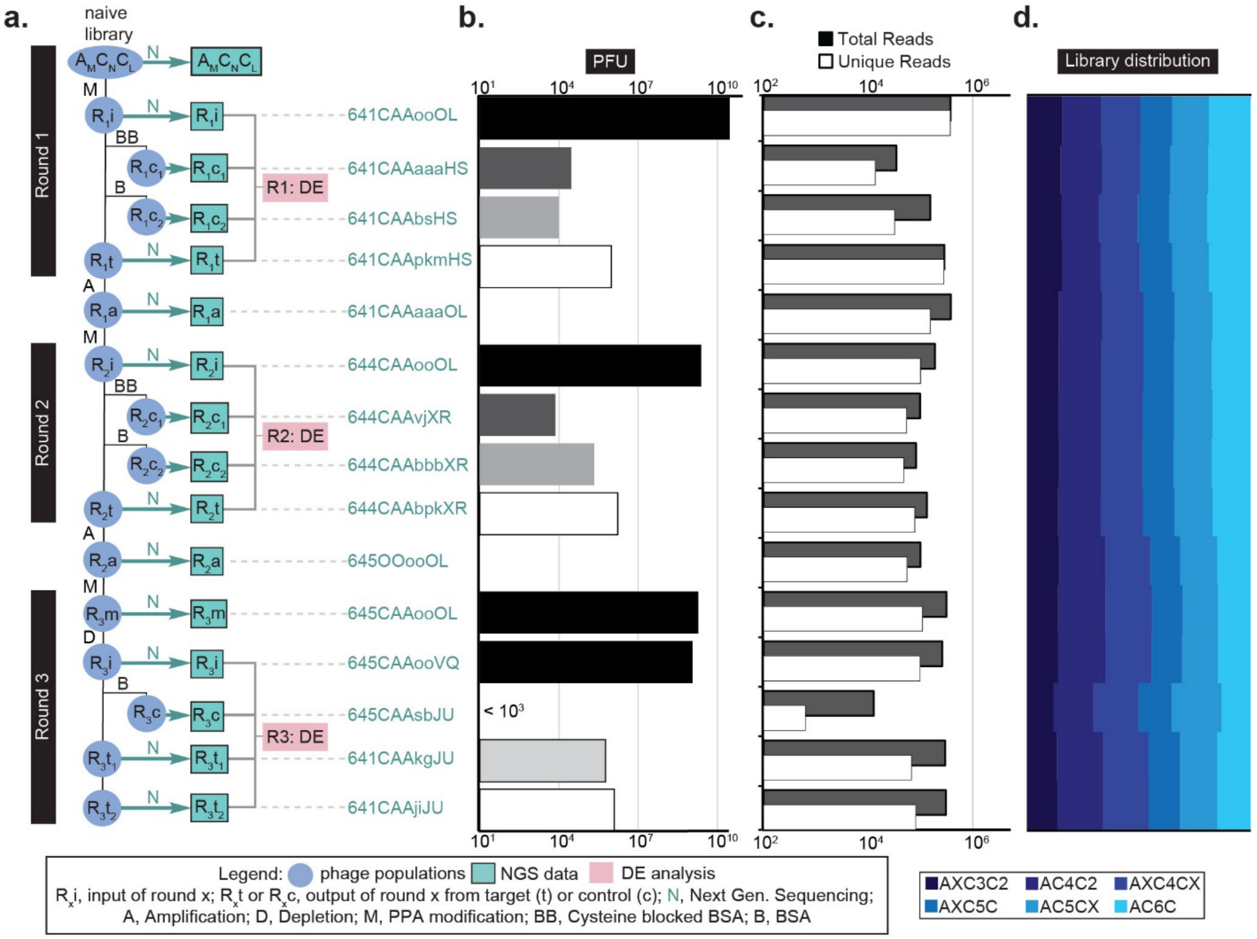

**Figure S1.11.** Summary of NGS data for multi-round panning (R1-R3).

**a**) Flowchart of 3-Round panning and names of sequencing files on 48HD cloud. **b**) PFU following iterative rounds of sequencing and depletion. **c**) Sequencing depth for each NGS output. **d**) Library sequence distribution

**PCR amplification protocol to prepare Phage DNA for Illumina deep sequencing**

Two-step semi-nested PCR was used to improve the sensibility (Figure S1.12). The method was adapted from our previous developed protocol.[1, 2] DNA template (phage solution) was amplified in Phusion® HF buffer with Phusion® High-Fidelity DNA Polymerase (NEB, #M0530S).

**$1^{st}$ step:**

A typical 50 µL reaction mixture contained:

1. 5x Phusion buffer 10 µL
2. 10 mM dNTPs 1 µL
3. DMSO 2.5 µL
4. Phusion® Polymeras 0.5 µL
5. Forward primer (TTTTGGAGATTTTCAACGTG, 10 µM) 1 µL
6. Reverse primer (CCCTCATAGTTAGCGTAACG, 10 µM) 1 µL
7. DNA Template solution 10 µL
8. Nuclease free water 24 µL

Cycling was performed using the following thermocycler settings:

a) 98 °C 3 min
b) 98 °C for 10 s
c) 50 °C 20 s
d) 72 °C 20 s
e) repeat b)-d) for 30 cycles
f) 12 °C 1 min
g) 4 °C hold

**$2^{st}$ step:**

A typical 50 µL reaction mixture contained:

1. 5x Phusion buffer 10 µL
2. 10 mM dNTPs 1 µL
3. DMSO 2.5 µL
4. Phusion® Polymeras 0.5 µL
5. Forward primer (10 µM) 1 µL
6. Reverse primer (10 µM) 1 µL
7. PCR product from $1^{st}$ step 2 µL
8. Nuclease free water 32 µL

Cycling was performed using the following thermocycler settings:

a) 98 °C 3 min
b) 98 °C for 10 s
c) 50 °C 20 s
d) 72 °C 20 s
e) repeat b)-d) for 20 cycles
f) 12 °C 1 min
g) 4 °C hold

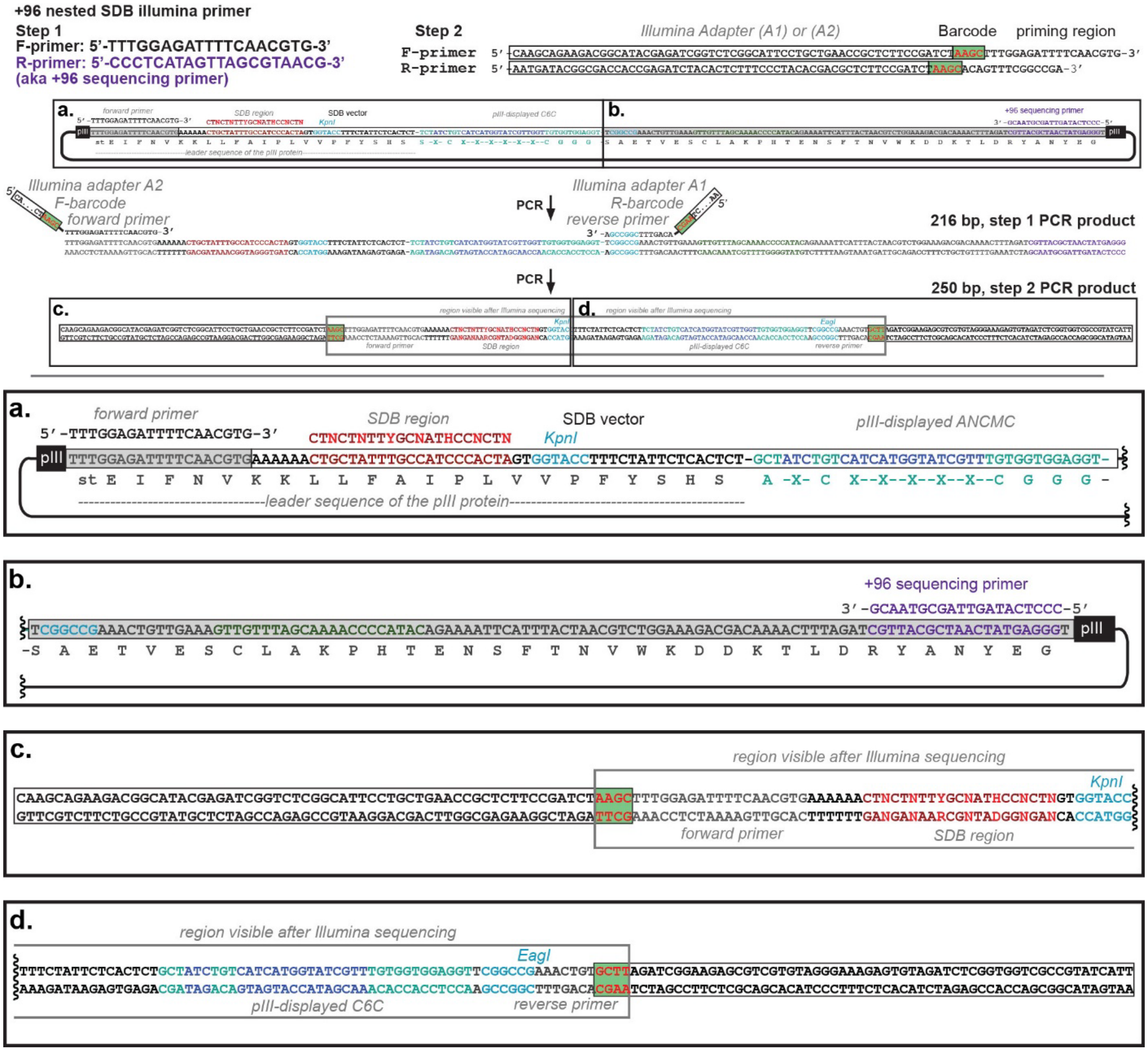

**Figure S1.12.** PCR amplification protocol for Illumina deep sequencing.
**a**) Zoom-in of section (a) Phage DNA prior to PCR showing the forward primer binding region and SDB region, and the ANCMC sequence to be displayed on pIII. **b**) Zoom-in of section (b) Phage DNA prior to PCR amplification showing the reverse primer (+96 primer) binding region. **c**) Zoom-in of section (c) Phage DNA after 2-step PCR amplification showing the primer barcode region, forward primer, SDB region, and the ANCMC sequence to be displayed on pIII. **d**) Zoom-in of section (d) Phage DNA after 2-step PCR amplification showing the reverse primer and primer barcode region.

**llumina Sequencing of samples before and after panning**

The DNA was produced by PCR as described in general PCR amplification protocol for Illumina deep sequencing with one exception: in amplification of libraries before panning (input), the template volume (phage solution) was 2 μL. All products were quantified by 2% (w/v) agarose gel in Tris-Borate-EDTA buffer at 100 volts for ~35 min using a low molecular weight DNA ladder as standard (NEB, cat# N3233S). PCR products that contain different indexing barcodes were pooled, allowing 10 ng of each product in the mixture. The mixture was purified by eGel, quantified by quBit and sequenced using the Illumina NextSeq paired-end 500/550 High Output Kit v2.5 (2×75 Cycles). Data were automatically uploaded to BaseSpace™ Sequence Hub. Processing of the data is described in the section describing processing of Illumina data.

**Processing of Illumina data**

The Gzip compressed FASTQ files were downloaded from BaseSpace™ Sequence Hub. The files were converted into tables of DNA sequences and their counts per experiment. Briefly, FASTQ files were parsed based on unique multiplexing barcodes within the reads discarding any reads that contained a low-quality score. Mapping the forward (F) and reverse (R) barcoding regions, mapping of F and R priming regions allowing no more than one base substitution each and F-R read alignment allowing no mismatches between F and R reads yielded DNA sequences located between the priming regions as described in previous publications.[2] The files with DNA reads, raw counts, and mapped peptide modifications were uploaded to http://48hd.cloud/ server (Table S1.1). Each experiment has a unique alphanumeric name and unique static URL (Figure S1.11 and Table S1.1).

**Table S1.1.** URL for R1-R3 PKM2 panning deep sequencing results.

<table>
<tr><th></th><th>PKM2</th><th>BSA</th><th>C-Blocked BSA</th></tr>
<tr><td>R1</td><td>https://48hd.cloud/file/9431</td><td>https://48hd.cloud/file/9329</td><td>https://48hd.cloud/file/9430</td></tr>
<tr><td>R2</td><td>https://48hd.cloud/file/9335</td><td>https://48hd.cloud/file/9334</td><td>https://48hd.cloud/file/9432</td></tr>
<tr><td rowspan="2">R3</td><td>https://48hd.cloud/file/9377</td><td rowspan="2">N/A</td><td rowspan="2">https://48hd.cloud/file/9380</td></tr>
<tr><td>https://48hd.cloud/file/9376</td></tr>
</table>

**Peptide synthesis**

Peptides were synthesized on a PreludeX peptide synthesizer (Gyros Protein Technologies) by standard Fmoc solid chemistry using Rink Amide AM resin. Exception: in peptides with C-terminal propargyl-glycine (**31** and **33**), the first amino-acid was loaded manually. Fmoc-protected amino acids, HBTU, Rink Amide AM resin were purchased from ChemPrep, Wellington FL USA. Peptides were cleaved from the resin by using a TFA/EDT/TIPS/Water (89.9/2.28/4.54/2.28 v/v) or TFA/thioanisole/1,2-ethanedithiol/anisole (90/5/3/2 v/v) for peptides with C-terminal propargyl-glycine. Cleaved peptides were precipitated and washed with ice-cold diethyl ether and purified by HPLC and lyophilized into the product.

**General chemistry methods**

Chemical reagents and solvents were purchased from Sigma-Aldrich or Fisher Scientific unless noted otherwise. Reagents for peptide synthesis were purchased from ChemPep; model peptides were synthesized using standard Fmoc solid phase synthesis as described above. Reactions were monitored by TLC which was carried out on silica gel 60 F254 (Merck) plates and visualized by UV-light (λ=254nm) and/or by spraying potassium permanganate or anisaldehyde followed by heating. Flash column chromatography was performed using silica gel 60 (40-63 μm). The subsequent evaporation of solvents was performed using IKA RV10 rotary evaporator. Proton ($^1$H NMR) and Carbon ($^{13}$C NMR) nuclear magnetic resonance spectra were recorded on an Agilent/Varian VNMRS two channel 500 MHz or Agilent/Varian Inova two-channel 400 MHz spectrometer. The chemical shifts are given in part per million (ppm) on the delta scale. The solvent peak was used as reference values. For $^1$H NMR: $CDCl_3$= 7.24 ppm and for $^{13}$C NMR: $CDCl_3$= 77.16 ppm. The following abbreviations have been used: s, singlet; d, doublet; t, triplet; m, multiplet. LCMS data was obtained on Agilent Technologies 6130 LCMS. A gradient of solvent A (MQ water) and solvent B ($MeCN/H_2O$ 95/5) was run at a flow rate of 0.5 mL/min (0-4.0 min 5% B; 4.0-5.0 min 5%/60% B; 5.0-5.50 min 60%/100% B; 5.5-7.5 100% B 7.5-11 min 100%/5% B). Analytical and preparative HPLC was conducted using Waters 1525 Binary pump equipped with a Waters Symmetry prep 19×50 mm C18 Columns and Waters 2489 UV detector. Removal of aqueous solvents was performed using Labconco Freezone 2.5w system.

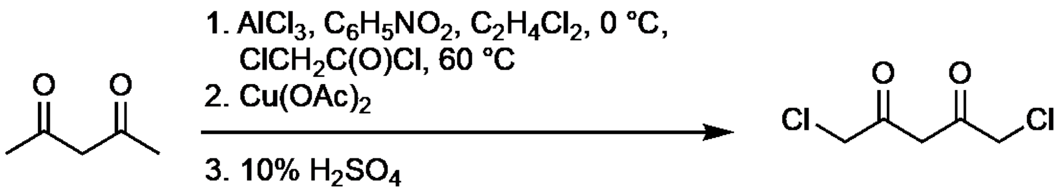

**Scheme. S1.1.** Synthesis of **DKL**.

2,4-pentadione was converted to 1,5-dichloro-2,4-pentadione (Supplementary Scheme 1) using previously described synthetic methods.[1] A three necked round bottom flask was charged with $AlCl_3$ (8 g, 60 mmol), nitrobenzene (10 mL), and dichloroethane (12 mL) under $N_2$. The mixture was stirred and 2,4-pentadione (6.15 mL, 60 mmol) was added dropwise. The new mixture was cooled to 0 ºC, before adding chloroacetyl chloride (9.5 mL, 120 mmol) dropwise over the course of 30 min. The reaction was heated to 60 ºC while stirring for 4 h and the reaction mixture was poured to a flask containing 10 mL of HCl in 80g of ice and the was stirred overnight. The aqueous layer was separated, and the products were extracted into diethyl ether. The combined organic layers were washed with brine and $H_2O$ before mixing with 100 mL of saturated $Cu(OAc)_2$ with shaking and the blue green precipitate was collected through filtration. The resulting crude product (12g, 3 mmol) was dissolved in diethyl ether and a solution of 10% $H_2SO_4$ (10 mL) by stirring. The mixture was then extracted with ethyl acetate. The combined organic layers were washed with brine and water, before drying over $Na_2SO_4$. The dried organic layer was concentrated in a rotary evaporator, and the product was separated from the crude material through distillation and isolated as a greenish brown solid (4.05 g, 43%). The characterization of 1,5-dichloro-2,4-pentadione was adapted from a previous report.[3] $^1$H NMR (400 MHz): δ 4.17 (4 H, s), 3.91 (2 H, s); $^{13}$C NMR (100 MHz); 196.2, 50.4, 48.2. $[M+H]^-$ calculated for $C_5H_6Cl_2O_2$ 166.9823, observed 166.9672.[3]

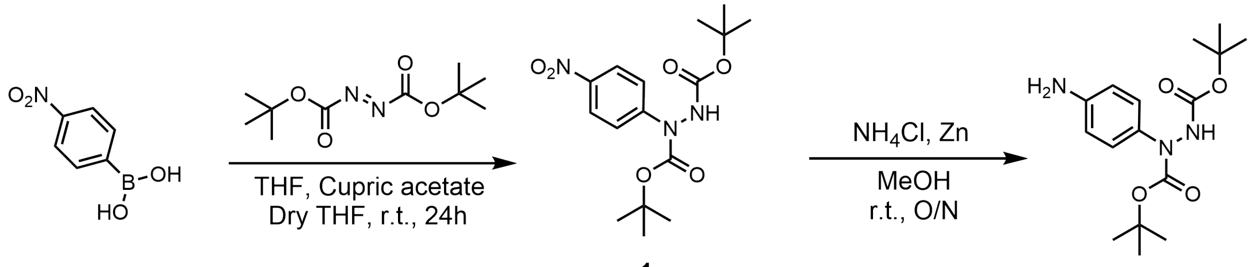

**Scheme. S1.2.** Synthesis of *N,N'*-(4-bis-Boc-hydrazinyl)-aniline (**2**)

To a dry three neck flask containing a solution of 4-nitrophenylboronic acid (4.2 g, 25 mmol) in 25 mL of anhydrous THF, di-tert-butyl azodicarboxylate was added while under stirring (0.5 equiv., 2.9 g, 12.5 mmol). A catalytic amount of $Cu(OAc)_2$ was added (0.05 equiv., 225 mg, 1.25 mmol) and the mixture was allowed to react for 24 hours under inert atmosphere. The reaction mixture was then purified by dry loaded silica flash column chromatography using gradient elution of DCM:EtOAc. The solvent was then evaporated at reduced pressure to afford the *N,N'*-(4-bis-Boc-hydrazinyl)-nitrophenyl (**1**) intermediate as a yellow solid (3.7 g, 84%) (Supplementary Scheme 2).

The intermediate (1.46 g, 4.1 mmol) was dissolved in 15 mL of MeOH. $NH_4Cl$ (5 equiv., 1.11 g, 20.5 mmol) and Zn(0) (5 equiv., 1.35 g, 20.5 mmol) was added and the mixture was stirred overnight at room temperature. The reaction mixture was then diluted with 15 mL EtOAc and the organic layer was washed three times with 10% $Na_2CO_3$ then dried with $MgSO_4$. The solvent was then evaporated under reduced pressure to afford *N,N'*-(4-bis-Boc-hydrazinyl)-aniline (**2**) as a bright yellow solid (1.14 g, 91%) (Supplementary Scheme 2).

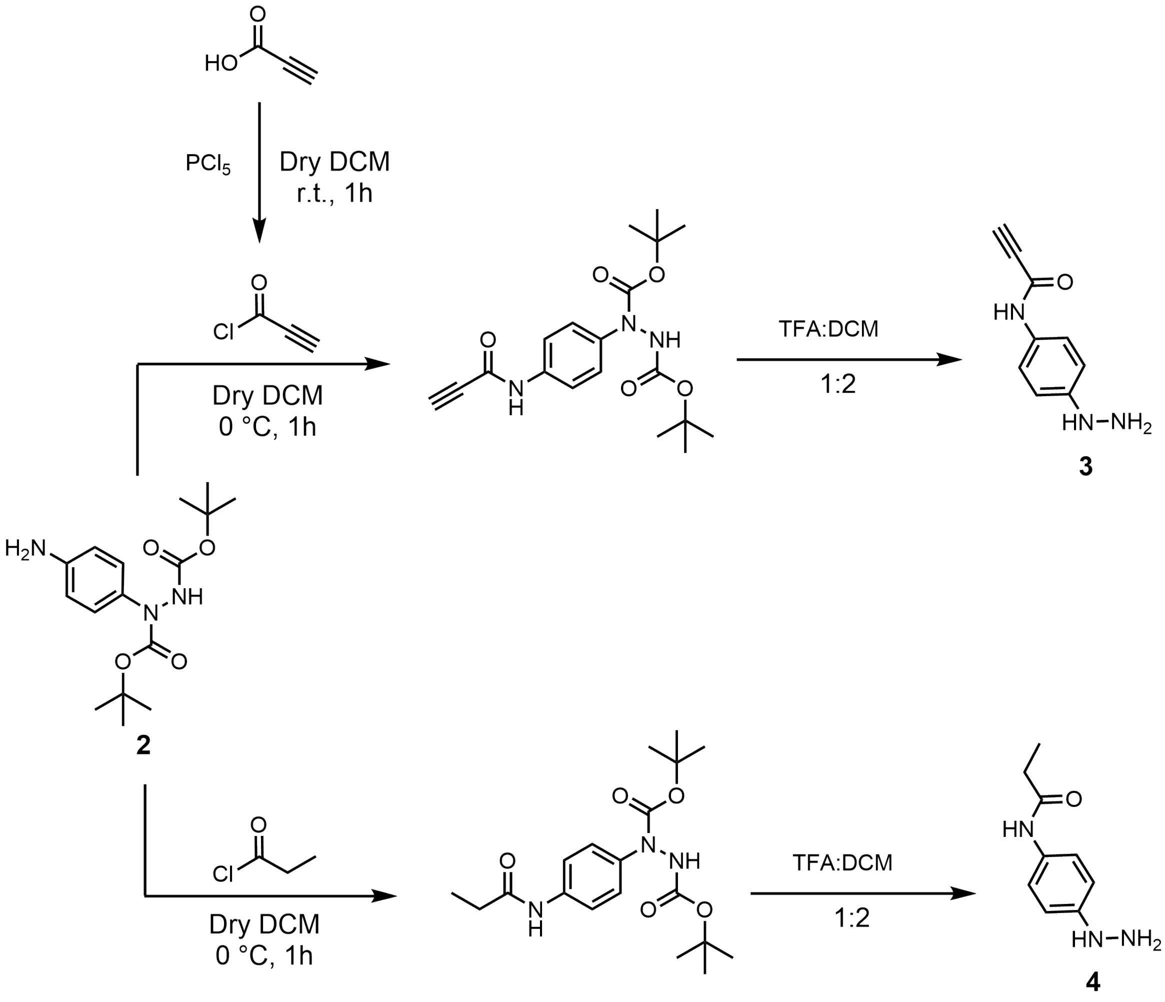

**Scheme. S1.3.** Synthesis of hydrazinyl Active (**3**) and Control warheads (**4**)

This is a representative procedure of the synthesis of *N*-(4-bis-Boc-Hydrazinylphenyl) propiolamide. The synthesis of the propanamide variant can be performed by the substitution of propynoyl chloride for propioyl chloride. The propynoyl chloride reagent is prepared *in-situ* from propiolic acid. To a dry flask containing 10 mL of dry DCM, propiolic acid (105 mg, 1.50 mmol) was added. $PCl_5$ (2 equiv., 0.62 g, 3 mmol) was added slowly and the mixture was stirred for one hour under inert atmosphere. The solvent was then evaporated under reduced pressure to afford a yellow liquid which was quantitatively transferred to the next step (Supplementary Scheme 3).

A dry three-neck flask containing a solution of **2** (181 mg, 0.57 mmol) in 10 mL of anhydrous DCM was cooled to 0 °C. Propynoyl chloride (2 equiv., 105 mg, 1.14 mmol) was added slowly. Upon completion of addition, the reaction mixture was stirred for one hour under an inert atmosphere and diluted with 20 mL DCM and 50 mL of sat. $NaHCO_3$. The organic layer was washed three times with sat. $NaHCO_3$, washed three times with brine, then dried with $MgSO_4$. The mixture was then purified by dry-loaded silica flash column chromatography using gradient elution of hexanes:EtOAc. The solvent was then evaporated at reduced pressure to afford the *N*-(4-bis-Boc-Hydrazinylphenyl)propiolamide intermediate as a yellow solid (175 mg, 82%). The intermediate (175 mg, 0.47 mmol) was transferred to a flask and 1.5 mL of a TFA:DCM 1:2 solution was added and the reaction mixture was stirred overnight. The solvent was removed by evaporation under reduced pressure and the remaining TFA was lyophilized overnight to afford *N*-(4-hydrazinylphenyl)propiolamide (**3**) (80 mg, 99%) (Supplementary Scheme 3). Compound **3** $^1H$ NMR (500 MHz) ($CD_3CN$: δ 1.93): δ 8.73 (1 H, s), 7.45 (2 H, d), 6.91 (2 H, d), 3.69 (3 H, br), 3.30 (1 H, s) (Supplementary Fig. 70). Compound **4** $^1H$ NMR (500 MHz) (d-DMSO: δ 2.5): δ 9.92 (2 H, br), 9.79 (1 H, s), 7.95 (1 H, br), 7.49 (2 H, d), 6.89 (2 H, d), 2.27 (2 H, q), 1.05 (3 H, t) (Supplementary Fig. 71).

**List of hits from phage-display screening**

**Table S1.2.** Sequences of the nominated peptides for activity profiling.

| Number | Sequence | a | b | c | d |
|---|---|---|---|---|---|
| | | SH (μM) | Diketone (DKL) (μM) | Alkyne (PPA) (μM) | Ethyl (PEA) (μM) |
| **AXC3C2** | | | | | |
| 5 | ASCLFNCPL | **NA** | **5b** | **5c** (49.95) | **NA** |
| 6 | AICTWNCIP | **NA** | **6b** | **6c** (412.8) | **NA** |
| 7 | ASCLFKCKS | **NA** | **7b** | **7c** (413.7) | **NA** |
| 8 | AECISFCRN | **NA** | **8b** | **8c** (72.37) | **NA** |
| 9 | ARCGMVCNQ | **NA** | **9b** | **9c** (197.5) | **NA** |
| **AXC4CX** | | | | | |
| 10 | AWCGVRTC | **NA** | **10b** | **10c** (97.97) | **NA** |
| 11 | ASCFFNTCH | **NA** | **11b** | **11c** (685.1) | **NA** |
| 12 | AFCSYDTCF | **NA** | **12b** | **12c** (480.6) | **NA** |
| 13 | ATCAWRNCT | **NA** | **13b** | **13c** (551.0) | **NA** |
| 14 | ASCFFSACK | **NA** | **14b** | **14c** (293.8) | **NA** |
| 15 | ANCPNYKCR | **15a** (>1000) | **15b** | **15c** (4.6 ± 0.6) | **15d** (35.16 ± 1.8) |
| **AC4C2** | | | | | |
| 16 | ACHTSICWL | **NA** | **16b** | **16c** (229.8) | **NA** |
| 17 | ACWKMNCLH | **NA** | **17b** | **17c** (77.54) | **NA** |
| 18 | ACRGFLCNY | **18a** (465.5) | **18b** | **18c** (1.54 ± 1.47) | **18d** (11.09 ± 1.5) |
| **AC5CX** | | | | | |
| 19 | ACHFDAFCT | **NA** | **19b** | **19c** (277.8) | **NA** |
| 20 | ACNFSTFCR | **20a** (228.7) | **20b** | **20c** (5.8 ± 1.3) | **20d** (180.3 ± 26) |
| 21 | ACVPYFRCI | **21a** (>1000) | **21b** | **21c** (8.2 ± 1.3) | **21d** (7.01 ± 0.44) |
| 22 | ACWQSWTCY | **NA** | **22b** | **22c** (67.45) | **NA** |
| **AC6C** | | | | | |
| 23 | ACASRFHEC | **NA** | **23b** | **23c** (104.0) | **NA** |
| 24 | ACVGSFFSC | **NA** | **24b** | **24c** (75.66) | **NA** |
| 25 | ACNFNKSSC | **NA** | **25b** | **25c** (163.5) | **NA** |
| 26 | ACLPFYSRC | **26a** (709.1) | **26b** | **26c** (6.08 ± 1.8) | **26d** (377 ± 161) |
| 27 | ACFFYSKNC | **27a** (>1000) | **27b** | **27c** (6.7 ± 1.2) | **27d** (582 ± 56) |
| 28 | ACTDRFFKC | **NA** | **28b** | **28c** (117.9) | **NA** |
| 29 | ACFHSSFFC | **NA** | **29b** | **29c** (N/A) | **NA** |

**Table S1.3.** Sequences synthesized for LCMS or competition studies.

| Number | Sequence | **a** | **b** | **c** | **d** |
|---|---|---|---|---|---|
| | | SH | Diketone (DKL) | Alkyne (PPA) | Ethyl (PEA) |
| **Competition study** | | | | | |
| 30 | RCGIGCV | **30a** | **NA** | **NA** | **NA** |
| **Protein-Peptide Adduct LCMS** | | | | | |
| 31 | ACFFYSKNCGGG(Z) | **NA** | **31b** | **31c** | **NA** |
| 32 | ACFFASKNC | **NA** | **32b** | **32c** | **NA** |
| 33 | ACFFASKNCGGG(Z) | **NA** | **33b** | **33c** | **NA** |

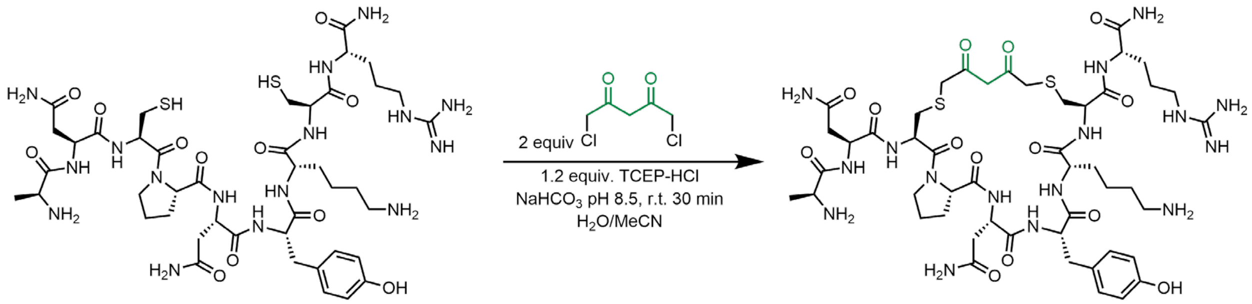

**Scheme. S1.4.** Cyclization of linear peptides by **DKL**.

This is a representative procedure for peptide ANCPNYKCR (**15a**) (Supplementary Scheme 4). To a peptide solution in water (5 mg in 2.5 mL, 5 µmol, 1 eq), a 500 mM stock of TCEP in water (0.012 mL, 6 µmol, 1.2 eq) and 2 mM 1,5-dichloro-2,4-pentadione solution in MeCN (5 µL, 10 µmol, 2 eq) were added. The reaction was initiated by increasing the pH of the solution by adding $NaHCO_3$ buffer (2.5 mL, 200 mM, pH 8.5). The reaction was incubated at room temperature for 30 min. After 30 min, the reaction mixture was filtered by a 0.2 micron filter and injected into preparative HPLC (C18, 0-2 min 2% MeCN, 2-26 min 2-50% MeCN, 26-27 min 50-100% MeCN, 27-28 min 100% MeCN). The collected fraction was lyophilized and peptide ANCPNYKCR-**DKL** (**15b**) was obtained as a fluffy light-yellow powder (4.1 mg, 75%). Apparent purity (95%) was estimated by LC-MS with UV-Vis detector at 214 nm.

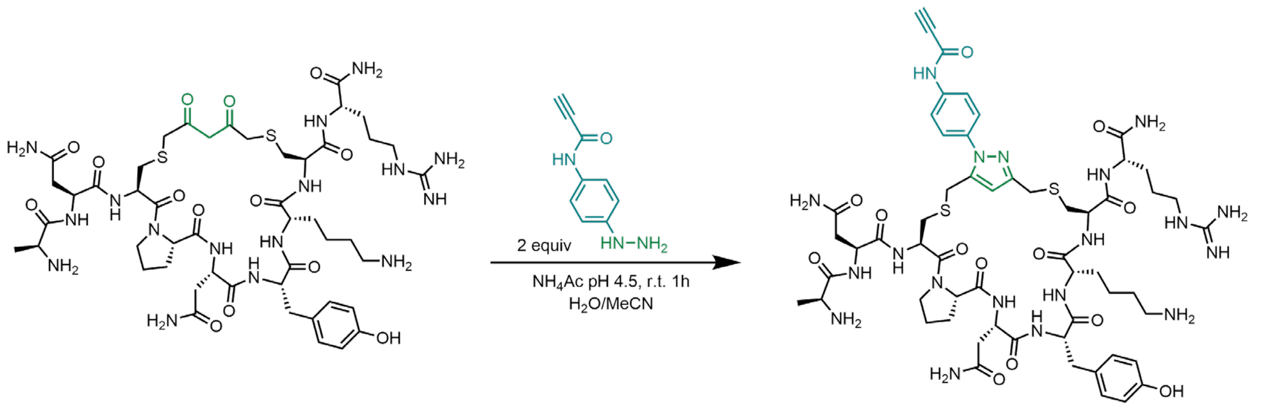

**Scheme. S1.5.** **PPA** functionalization of **DKL**-cyclized peptides.

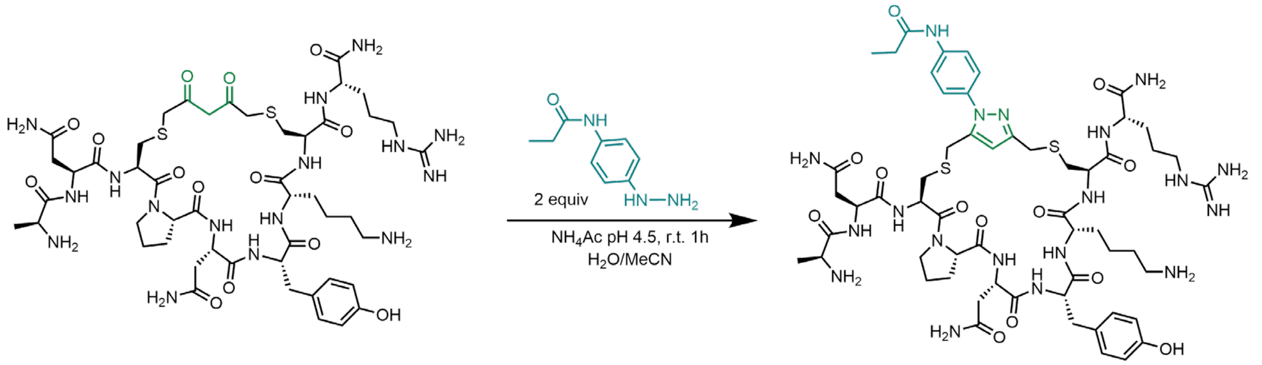

**Scheme. S1.6.** **PEA** functionalization of **DKL**-cyclized peptides**.**

This is a representative procedure for peptide ANCPNYKCR-**DKL** (**15b**) (Supplementary Scheme 5). The HPLC purified diketone macrocyclic peptide from the previous step was dissolved in water (3 mg in 240 µL, 3 µmol, 1 eq) followed by the addition of compound **3** (or compound **4** (Supplementary Scheme 6)) solution in MeCN (0.2 M, 30 µL, 60 µmol, 20 eq). The reaction was initiated by adding ammonium acetate buffer (1 M, 30 µL). After 1 hour, the reaction mixture was filtered by a 0.2 micron filter and injected into preparative HPLC (C18, 0-2 min 2% MeCN, 2-26 min 2-50% MeCN, 26-27 min 50-100% MeCN, 27-28 min 100% MeCN). The collected fraction was lyophilized and peptide ANCPNYKCR-**PPA** (**15c**) or ANCPNYKCR-**PEA** (**15d**) was obtained as a fluffy white powder (1.2 mg, 40%) in a racemic mixture of both regioisomers. Apparent purity (95%) was estimated by LC-MS with UV-Vis detector at 214 nm.

### PKM2 Activity Assay

Compounds **5c-29c** were incubated with hPKM2 (500 nM) for 2 h at 37 °C in 50 μL of buffer provided with the Pyruvate Kinase Activity Assay kit (Abcam, ab83432). $IC_{50}$'s were measured following the provided fluorometric protocol from the kit. Fluorescence was measured for 45 min. after reaction mix was added at each concentration of inhibitor. Same procedure was used for measuring the activity of select disulfide (**SH**) peptides and peptides bearing the control warhead (**PEA**).

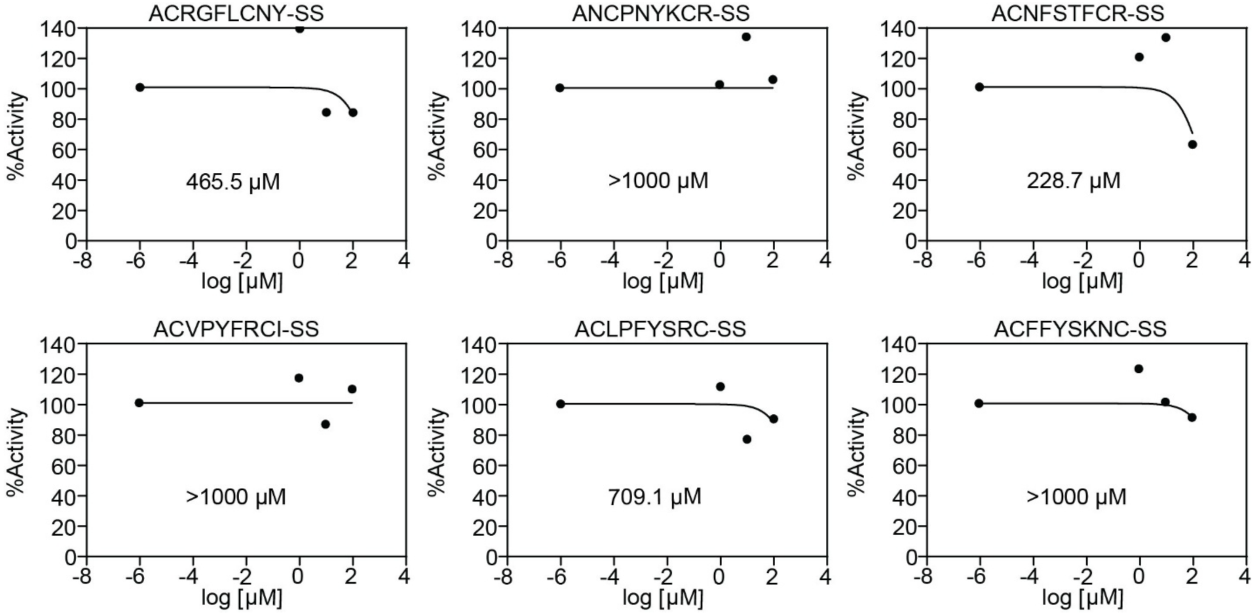

**Figure S1.13.** $IC_{50}$ curves for 6 nominated disulfide (unmodified) sequences.
Disulfide $IC_{50}$ curves for **15a**, **18a**, **20a**, **21a**, **26a**, and **27a**.

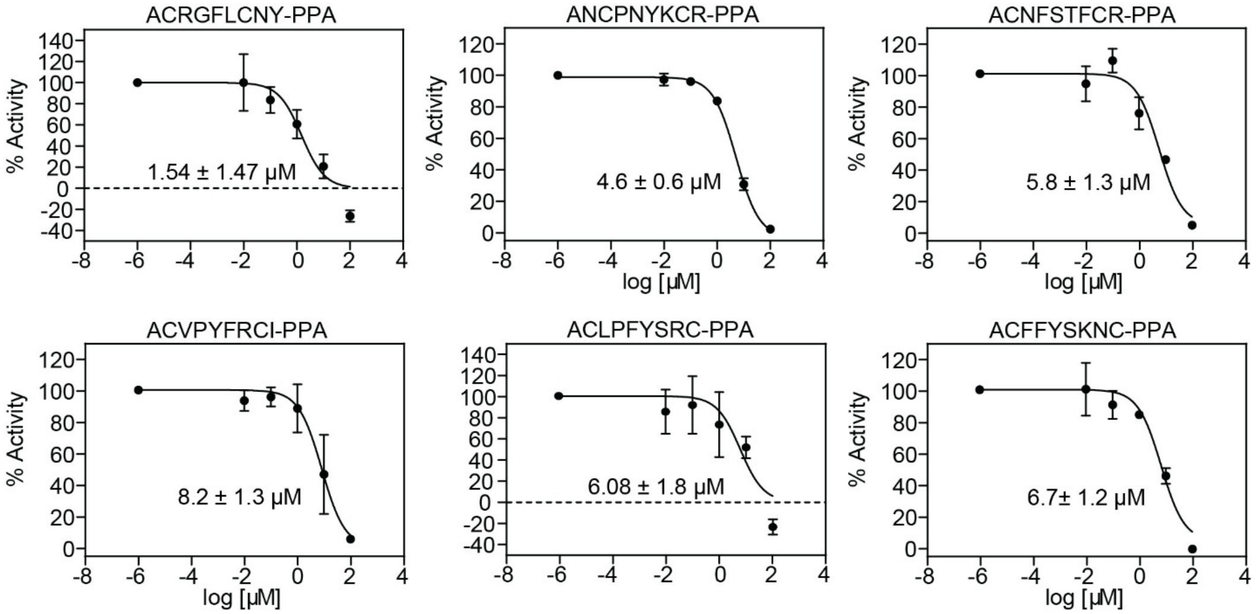

**Figure S1.14.** $IC_{50}$ curves for 6 nominated **PPA** modified sequences.
Disulfide $IC_{50}$ curves for **15c**, **18c**, **20c**, **21c**, **26c**, and **27c**.

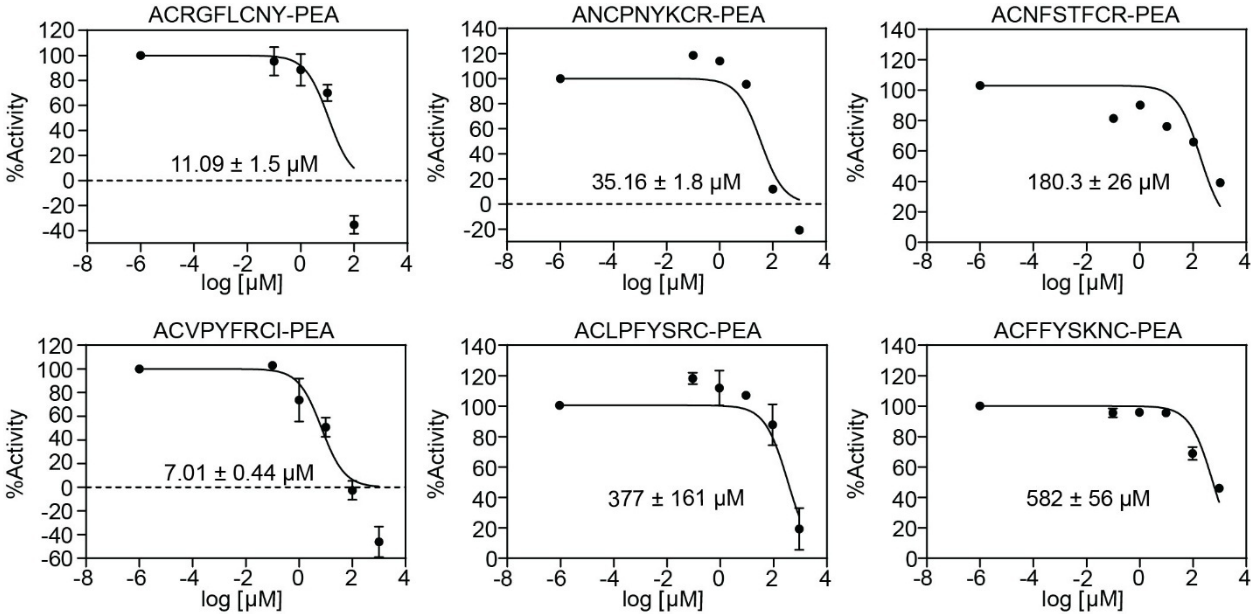

**Figure S1.15.** $IC_{50}$ curves for 6 nominated **PEA** modified sequences.
Disulfide $IC_{50}$ curves for **15d**, **18d**, **20d**, **21d**, **26d**, and **27d**.

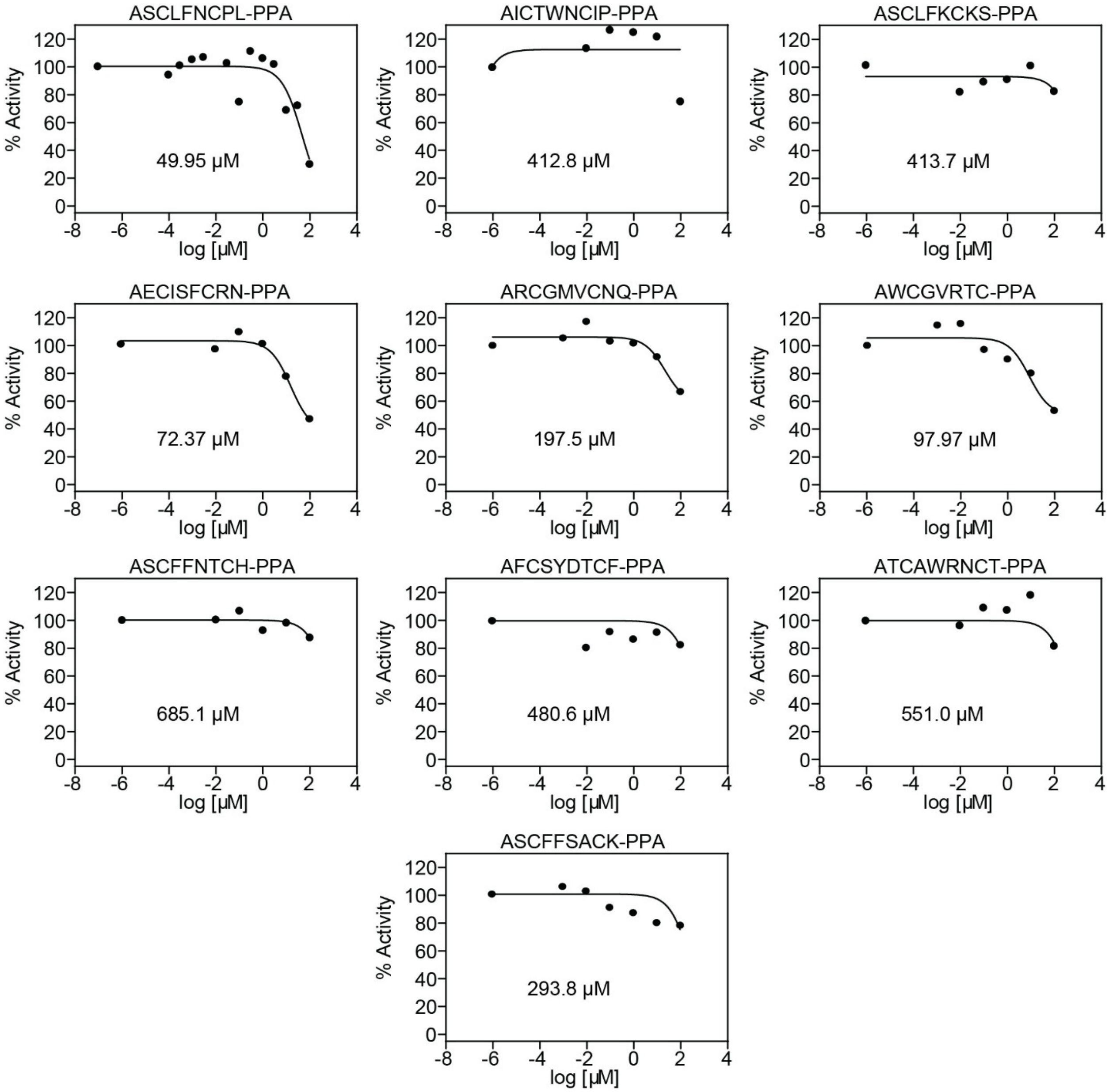

**Figure S1.16.** $IC_{50}$ curves for remaining **PPA** modified peptides.
**PPA**-Modified $IC_{50}$ curves for **5c** - **14c**.

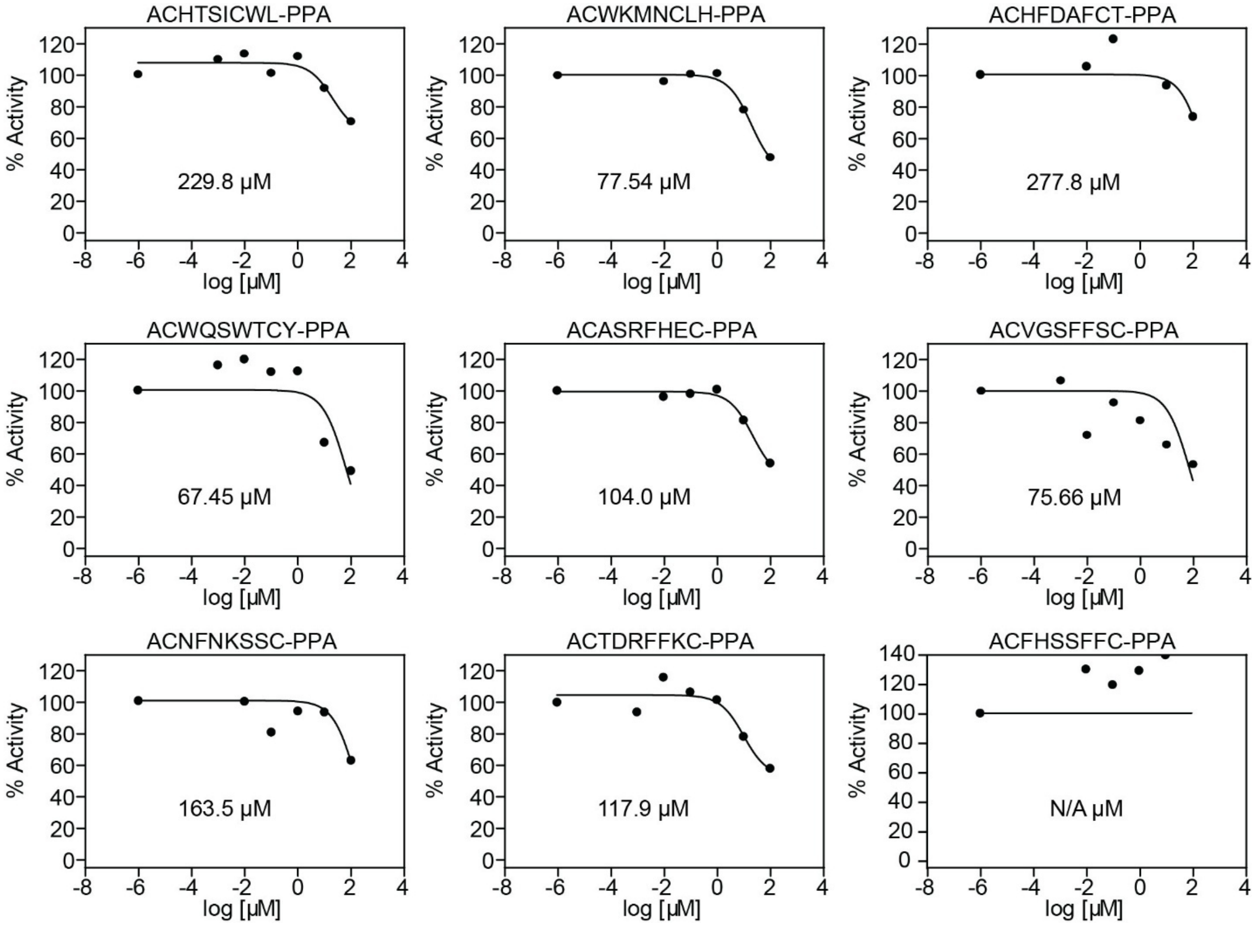

**Figure S1.17.** $IC_{50}$ curves for **PPA** modified peptides.
**PPA** Modified IC50 curves for **16c**, **17c**, **19c**, **22c-25c**, **28c**, and **29c**.

### Molecular Docking Calculations

#### Macrocycle Structural Models.

The structure of the ACFFYSKNC peptide was prepared by mutating an existing nine amino acid template with a coil backbone conformation. The covalent linker (warhead) fragment was built separately with GaussView 6.1.1 software.[4] The peptide was parameterized within the CHARMM36 force field,[5] solvated in TIP3P water, and its structure was minimized using NAMD 2.3.[6] To induce cyclization, a collective variable with a harmonic potential was applied to the sulfur atoms of the two cysteine residues (Cys2 and Cys9), reducing their separation to ~6.5 Å in a short molecular dynamics simulation. The peptide and warhead segments were then covalently joined using the psfgen plugin in VMD,[7] producing the final ACFFYSKNC-based macrocycle containing the warhead. The resulting ligand structure, which had reasonable bond lengths and angles, was used as input for docking studies.

#### Macrocycle Docking.

Structures of macrocycle-PKM2 complexes were calculated with AutoDock Vina,[8] using the PKM2 crystal structure (PDB ID: 6nu5)[9] and the peptide-warhead structures described above. Docking was performed with both the crystallographic dimer and the isolated monomer (one protein chain extracted from the crystal structure). For each structure, the grid box was centered on one of the ten surface-exposed cysteine residues of PKM2 (Cys31, Cys49, Cys152, Cys165, Cys317, Cys326, Cys358, Cys423, Cys424, Cys474 in chain A). The default exhaustiveness parameter (8) was used. For each docking run, we obtained the poses of the complex with the most favorable docking scores and calculated the distance between the cysteine sulfur atom and the center of mass of the warhead's C≡C bond. Only a small fraction of poses had this distance below 14 Å (Fig. 6b).

**Labelling protocol for LCMS study**

For individual labeling experiments, hPKM2 was diluted to 10 μM in PBS and incubated with 100 μM inhibitor (from a 10 mM DMSO stock) or DMSO control at 37 °C for 2 hours. After incubation, samples were desalted using Zeba desalting columns (Thermo Scientific cat no. 89882) and diluted to 1 μM in PBS. Samples were injected onto an Agilent 1200 HPLC equipped with a Zorbax SB-C3 column (1.8 μm, 2.1 × 150 mm, 300 Å pore size) coupled to an Agilent 6125B Single Quad Mass Spectrometer. For LC-MS analysis, samples were run at 0.6 mL/min with 95% solvent A (0.1% formic acid in water) and 5% solvent B (0.1% formic acid in acetonitrile) for 2 minutes, followed by a 6-minute linear gradient from 80% solvent A to 20% solvent A, then held for 1 minute at 95% solvent A. Protein masses were deconvoluted using the Agilent Bioanalysis Software package. For mixed labeling experiments, the experiment was the same was described above with the following change: 25 μM of each inhibitor for a total inhibitor concentration of 100 μM was incubated with 10 μM of protein. Protocol followed using compound **27c, 31c, 32c,** and **33c** (Supplementary Fig.18).

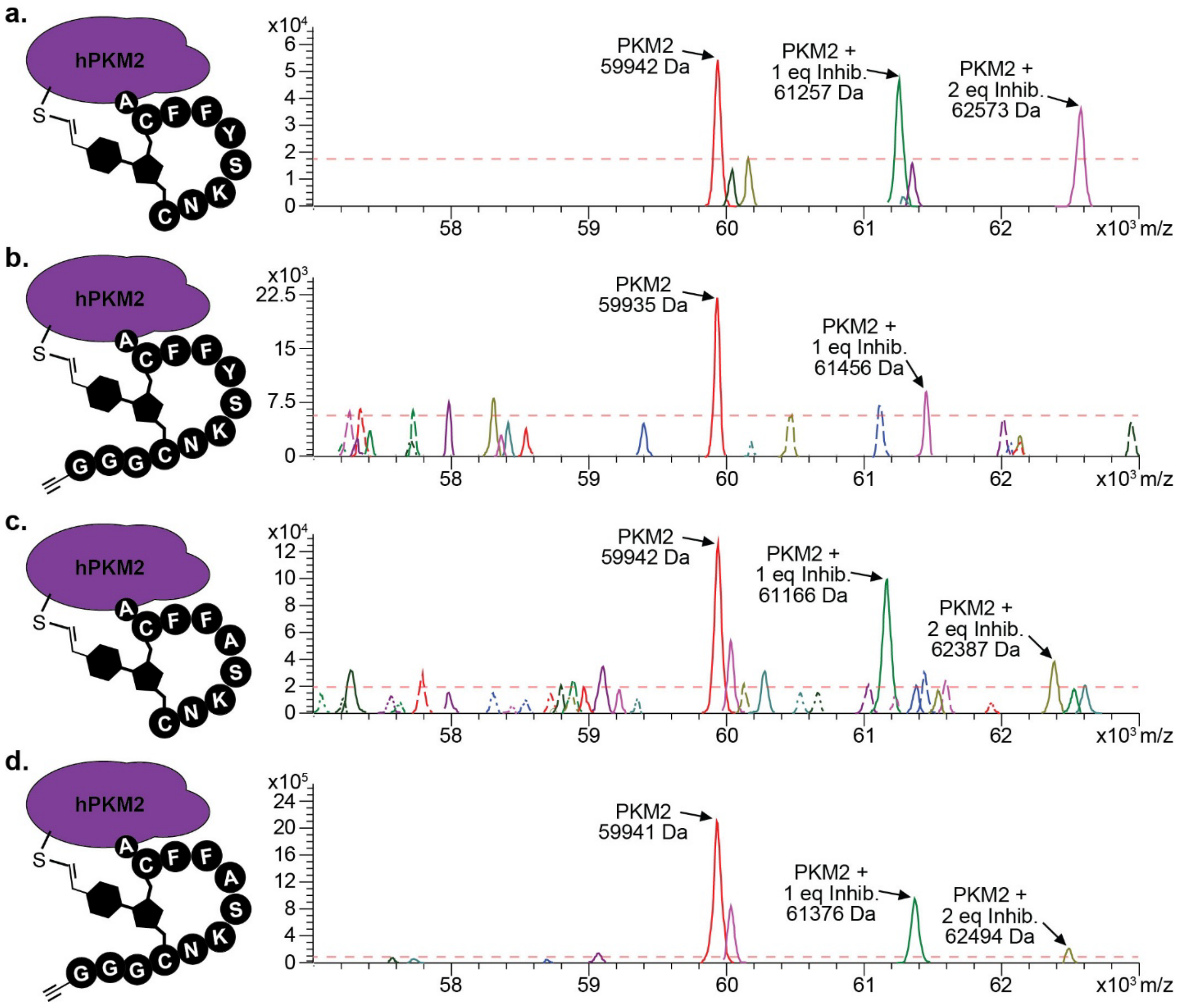

**Figure S1.18.** Monoisotopic MS peaks of PKM2 with **27c** and analogues.
**a**) MS of sarent sequence (**27c**) shows mass peaks for PKM2, PKM2 + 1 eq. of **27c**, and PKM2 + 2 eq of **27c**. **b**) MS of the -GGG(Z) modified sequence (**31c**) shows mass peaks for PKM2 and PKM2 + 1 eq. of **31c**. **c**) MS of the Tyrosine alanine scan (**32c**) shows mass peaks for PKM2, PKM2 + 1 eq. of **32c**, and PKM2 + 2 eq of **32c**. **d**) MS of the -GGG(Z) modified Tyrosine alanine scan (**33c**) shows mass peaks for PKM2, PKM2 + 1 eq. of **33c**, and PKM2 + 2 eq of **33c**.
