## Supporting information 2 for "A Two-Step Synthesis of Covalent Genetically-Encoded Libraries of Peptide-Derived Macrocycles (cGELs) enables use of electrophiles with diverse reactivity"

#### Summary of syntheses

##### ASCLFNCPL-DKL

Starting material mass: mg
Final product mass: mg

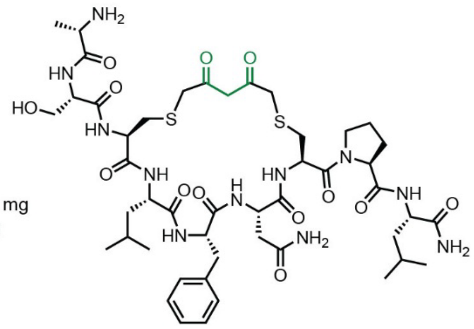

Chemical Formula: $C_{47}H_{71}N_{11}O_{13}S_2$
Exact Mass: 1061.4674
Molecular Weight: 1062.2690

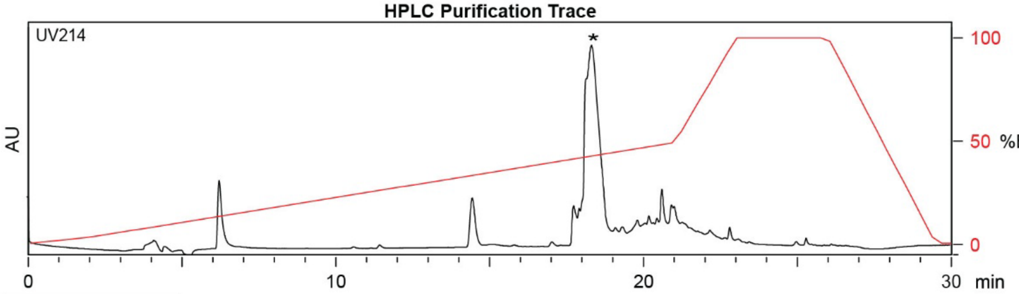

| Time (min) | Solvent B (%) |
|---|---|
| 0 | 2 |
| 2 | 2 |
| 22 | 50 |
| 24 | 100 |
| 27 | 100 |
| 28 | 2 |
| 30 | 2 |

Flow Rate: 13 mL / min
Phenomenex Kinetex EVO C18 Prep Column
(100 Å, 5 µm, 21.5 mm X 250 mm)

Solvent A: $H_2O$ + 0.1% (v/v) TFA
Solvent B: MeCN + 0.1% (v/v) TFA

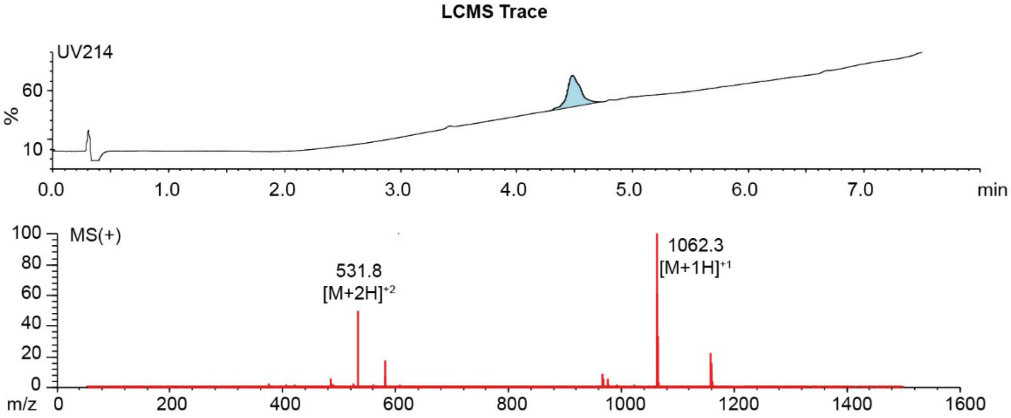

**Figure S2.1.** Synthesis summary of ASCLFNCPL-**DKL** (**5b**).
Expected: [M+1] = 1062.4674, $[M+2]^{+2}$ = 531.7337. Observed: [M+1] = 1062.3, $[M+2]^{+2}$ =531.8.

**ASCLFNCPL-PPA**

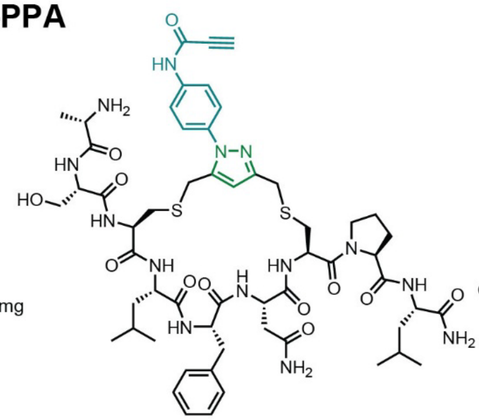

Starting material mass: mg
Final product mass: mg

Chemical Formula: $C_{56}H_{76}N_{14}O_{12}S_2$
Exact Mass: 1200.5209
Molecular Weight: 1201.4300

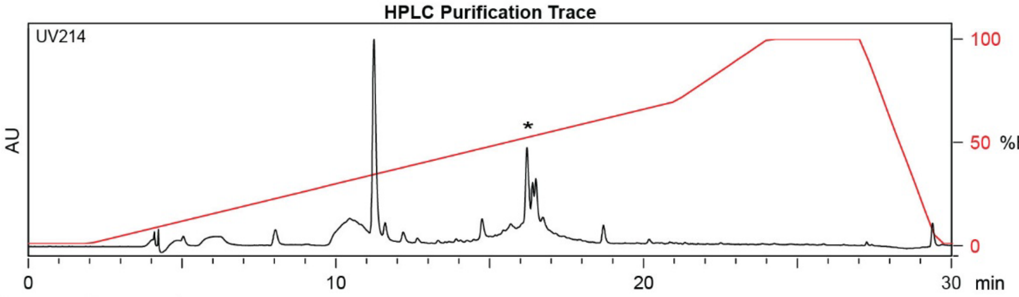

| Time (min) | Solvent B (%) |
|---|---|
| 0 | 2 |
| 2 | 2 |
| 21 | 70 |
| 24 | 100 |
| 27 | 100 |
| 29.5 | 2 |
| 30 | 2 |

Flow Rate: 13 mL / min
Phenomenex Kinetex EVO C18 Prep Column
(100 Å, 5 µm, 21.5 mm X 250 mm)

Solvent A: $H_2O$ + 0.1% (v/v) TFA
Solvent B: MeCN + 0.1% (v/v) TFA

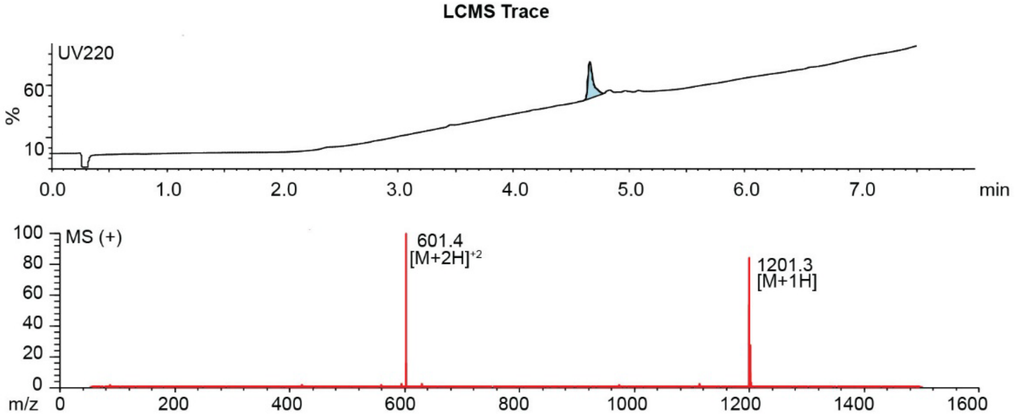

**Figure S2.2.** Synthesis summary of ASCLFNCPL-**PPA** (**5c**).
Expected: [M+1] = 1201.5209, $[M+2]^{+2}$ = 601.2605. Observed: [M+1] = 1201.3, $[M+2]^{+2}$ = 601.4.
Note: Two regioisomeric forms of the pyrazole are possible; the structure shown represents one isomer.

#### AICTWNCIP-DKL

Starting material mass: mg
Final product mass: mg

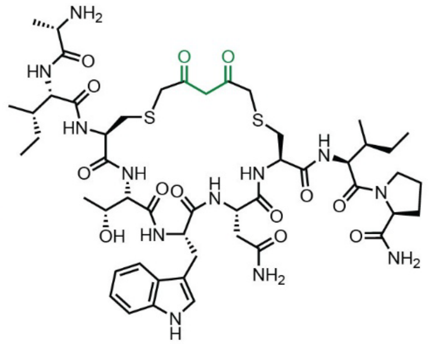

Chemical Formula: $C_{50}H_{74}N_{12}O_{13}S_2$
Exact Mass: 1114.4940
Molecular Weight: 1115.3330

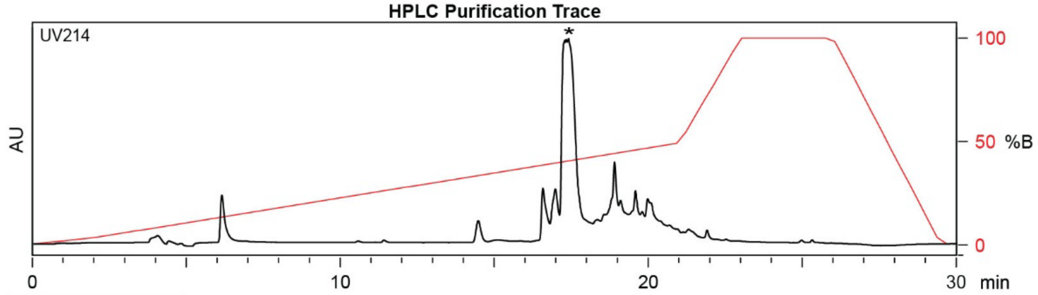

| Time (min) | Solvent B (%) |
|---|---|
| 0 | 2 |
| 2 | 2 |
| 22 | 50 |
| 24 | 100 |
| 27 | 100 |
| 28 | 2 |
| 30 | 2 |

Flow Rate: 13 mL / min
Phenomenex Kinetex EVO C18 Prep Column
(100 Å, 5 µm, 21.5 mm X 250 mm)

Solvent A: $H_2O$ + 0.1% (v/v) TFA
Solvent B: MeCN + 0.1% (v/v) TFA

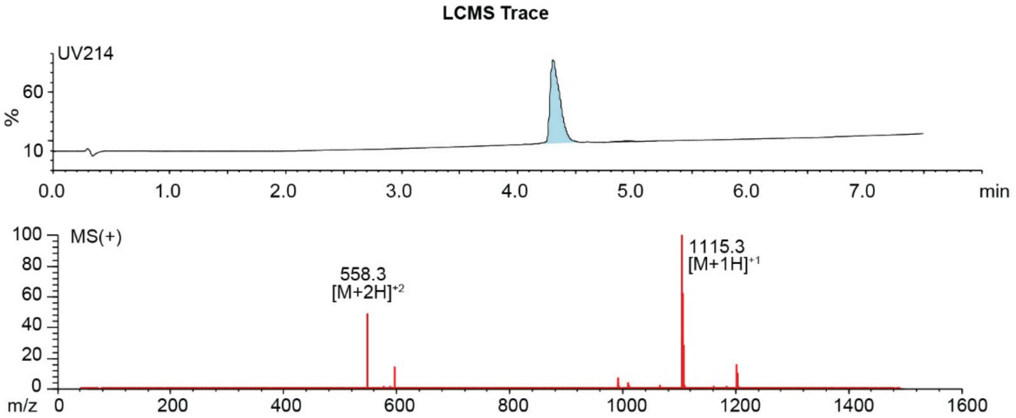

**Figure S2.3.** Synthesis summary of AICTWNCIP-**DKL** (**6b**).
Expected: [M+1] = 1115.4940, $[M+2]^{\div 2}$ = 558.2470. Observed: [M+1] = 1115.3, $[M+2]^{\div 2}$ = 558.3.

#### AICTWNCIP-PPA

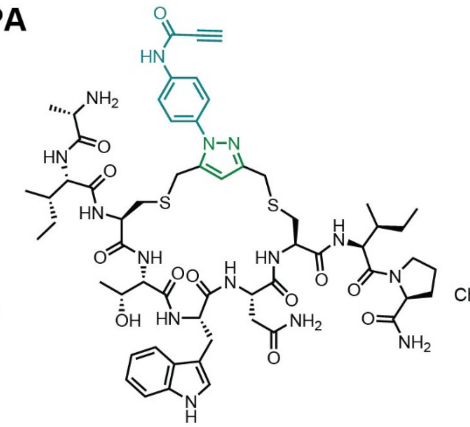

Starting material mass: mg
Final product mass: mg

Chemical Formula: $C_{59}H_{79}N_{15}O_{12}S_2$
Exact Mass: 1253.5474
Molecular Weight: 1254.4940

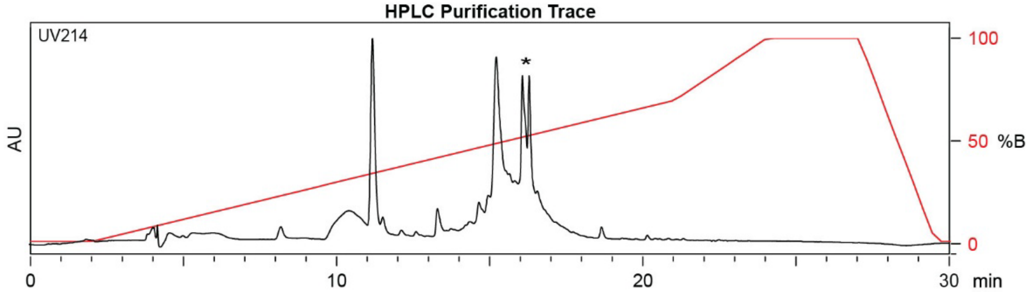

| Time (min) | Solvent B (%) |
|---|---|
| 0 | 2 |
| 2 | 2 |
| 21 | 70 |
| 24 | 100 |
| 27 | 100 |
| 29.5 | 2 |
| 30 | 2 |

Flow Rate: 13 mL / min
Phenomenex Kinetex EVO C18 Prep Column
(100 Å, 5 µm, 21.5 mm X 250 mm)

Solvent A: $H_2O$ + 0.1% (v/v) TFA
Solvent B: MeCN + 0.1% (v/v) TFA

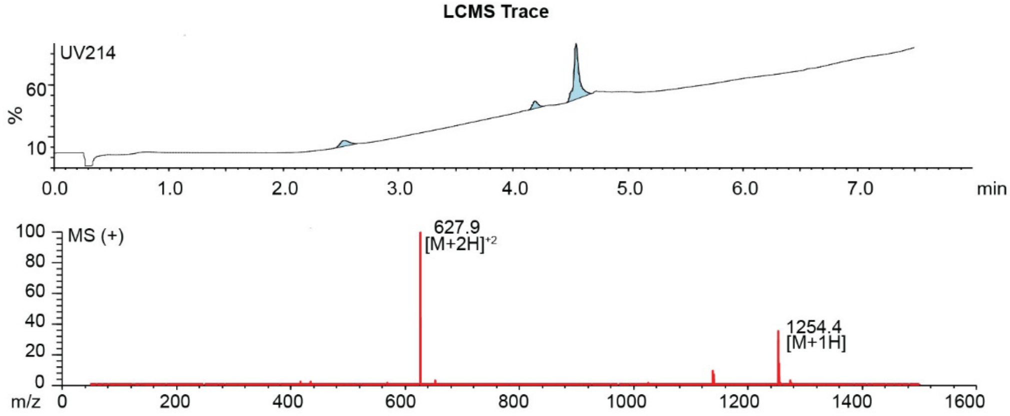

**Figure S2.4.** Synthesis summary of AICTWNCIP-**PPA** (**6c**).
Expected: [M+1] = 1254.5474, $[M+2]^{+2}$ = 627.7737. Observed: [M+1] = 1254.4, $[M+2]^{+2}$ = 627.9.
Note: Two regioisomeric forms of the pyrazole are possible; the structure shown represents one isomer.

#### ASCLFKCKS-DKL

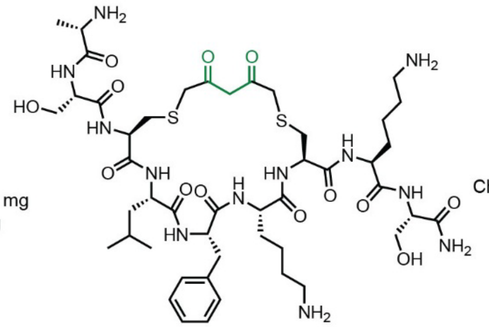

Starting material mass: mg
Final product mass: mg

Chemical Formula: $C_{47}H_{76}N_{12}O_{13}S_2$
Exact Mass: 1080.5096
Molecular Weight: 1081.3160

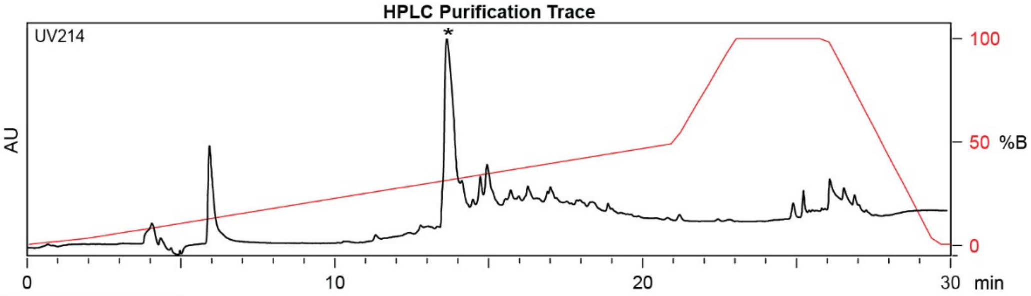

| Time (min) | Solvent B (%) |
|---|---|
| 0 | 2 |
| 2 | 2 |
| 22 | 50 |
| 24 | 100 |
| 27 | 100 |
| 28 | 2 |
| 30 | 2 |

Flow Rate: 13 mL / min
Phenomenex Kinetex EVO C18 Prep Column
(100 Å, 5 µm, 21.5 mm X 250 mm)

Solvent A: $H_2O$ + 0.1% (v/v) TFA
Solvent B: MeCN + 0.1% (v/v) TFA

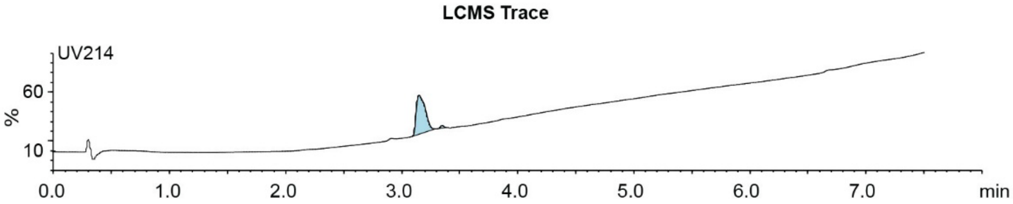

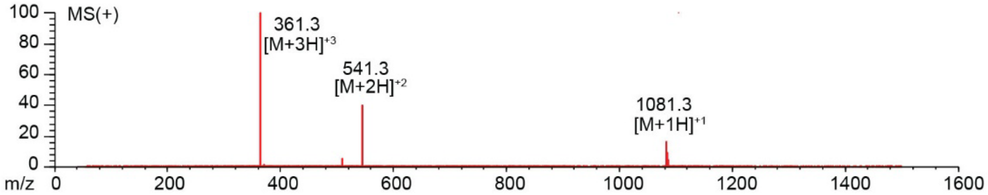

**Figure S2.5.** Synthesis summary of ASCLFKCKS-**DKL** (**7b**).
Expected: [M+1] = 1081.5096, $[M+2]^{\div 2}$ = 541.2548, $[M+3]^{\div 3}$ = 361.1698. Observed: [M+1] = 1081.3, $[M+2]^{\div 2}$ = 541.3, $[M+3]^{\div 3}$ = 361.3.

#### ASCLFKCKS-PPA

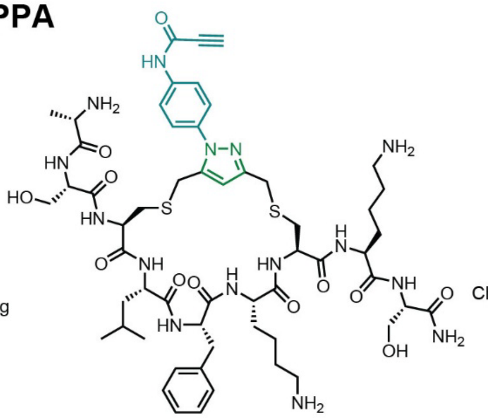

Starting material mass: mg
Final product mass: mg

Chemical Formula: $C_{56}H_{81}N_{15}O_{12}S_2$
Exact Mass: 1219.5631
Molecular Weight: 1220.4770

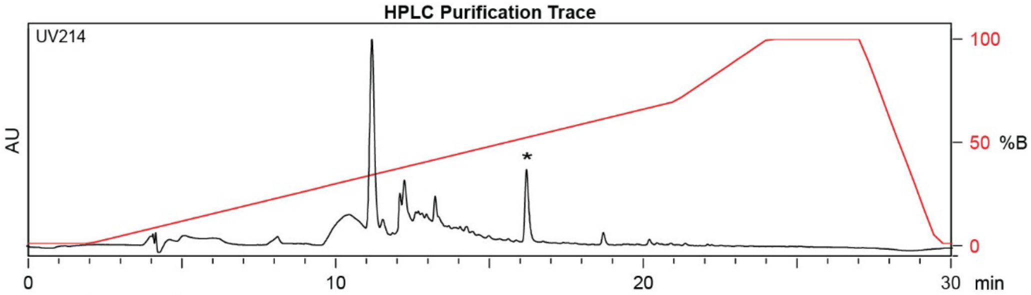

| Time (min) | Solvent B (%) |
|---|---|
| 0 | 2 |
| 2 | 2 |
| 21 | 70 |
| 24 | 100 |
| 27 | 100 |
| 29.5 | 2 |
| 30 | 2 |

Flow Rate: 13 mL / min
Phenomenex Kinetex EVO C18 Prep Column
(100 Å, 5 µm, 21.5 mm X 250 mm)

Solvent A: $H_2O$ + 0.1% (v/v) TFA
Solvent B: MeCN + 0.1% (v/v) TFA

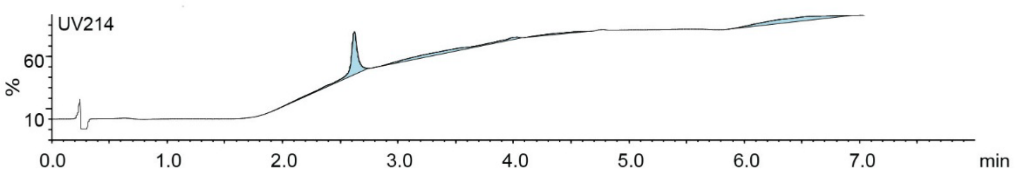

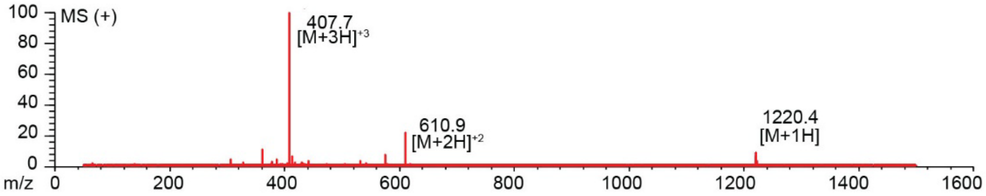

**Figure S2.6.** Synthesis summary of ASCLFKCKS-**PPA** (**7c**).
Expected: [M+1] = 1220.5631, $[M+2]^{+2}$ = 610.7816, $[M+3]^{+3}$ = 407.5210. Observed: [M+1] = 1220.4, $[M+2]^{+2}$ = 610.9, $[M+3]^{+3}$ = 407.7. Note: Two regioisomeric forms of the pyrazole are possible; the structure shown represents one isomer.

#### AECISFCRN-DKL

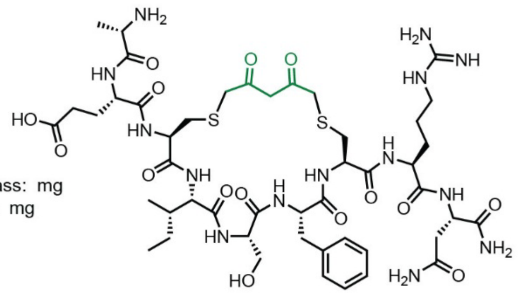

Starting material mass: mg
Final product mass: mg

Chemical Formula: $C_{47}H_{72}N_{14}O_{15}S_2$
Exact Mass: 1136.4743
Molecular Weight: 1137.2960

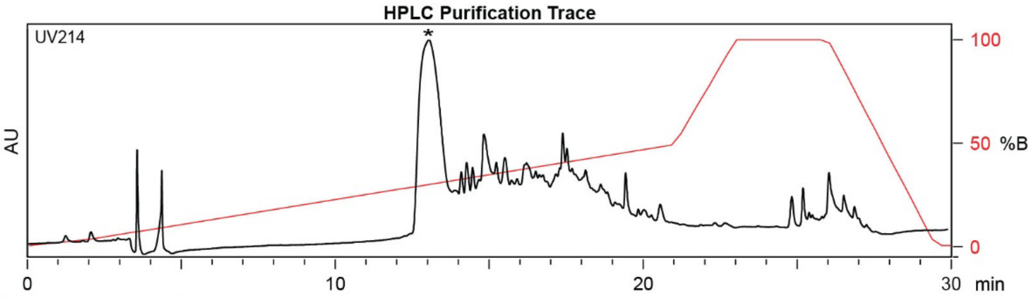

| Time (min) | Solvent B (%) |
|---|---|
| 0 | 2 |
| 2 | 2 |
| 22 | 50 |
| 24 | 100 |
| 27 | 100 |
| 28 | 2 |
| 30 | 2 |

Flow Rate: 13 mL / min
Phenomenex Kinetex EVO C18 Prep Column
(100 Å, 5 μm, 21.5 mm X 250 mm)

Solvent A: $H_2O$ + 0.1% (v/v) TFA
Solvent B: MeCN + 0.1% (v/v) TFA

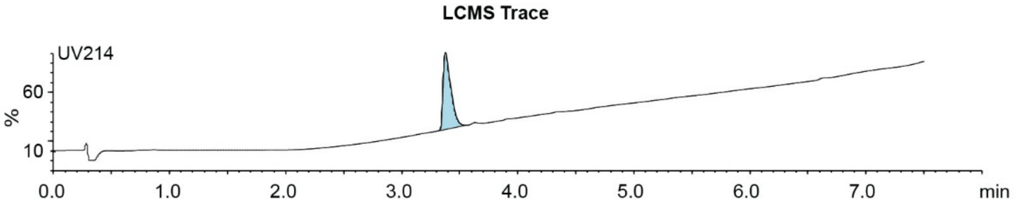

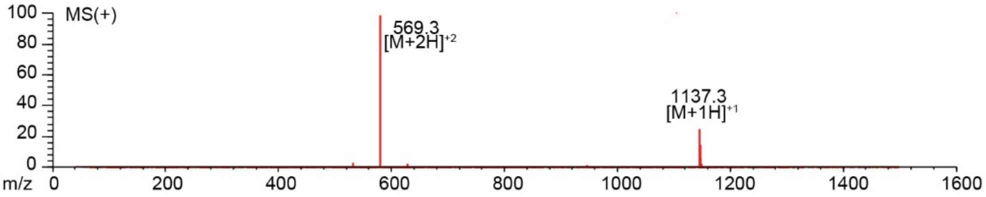

**Figure S2.7.** Synthesis summary of AECISFCRN-**DKL** (**8b**).
Expected: [M+1] = 1137.4743, $[M+2]^{\div 2}$ = 569.2372. Observed: [M+1] = 569.3, $[M+2]^{\div 2}$ = 1137.3.

#### AECISFCRN-PPA

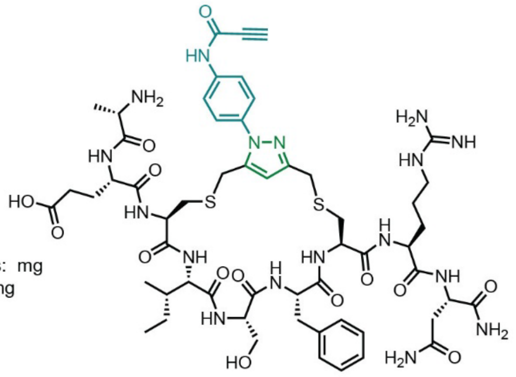

Starting material mass: mg
Final product mass: mg

Chemical Formula: $C_{56}H_{77}N_{17}O_{14}S_2$
Exact Mass: 1275.5277
Molecular Weight: 1276.4570

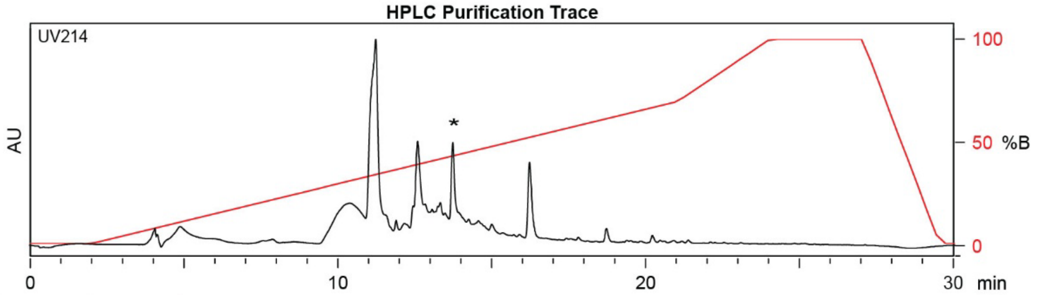

| Time (min) | Solvent B (%) |
|---|---|
| 0 | 2 |
| 2 | 2 |
| 21 | 70 |
| 24 | 100 |
| 27 | 100 |
| 29.5 | 2 |
| 30 | 2 |

Flow Rate: 13 mL / min
Phenomenex Kinetex EVO C18 Prep Column
(100 Å, 5 µm, 21.5 mm X 250 mm)

Solvent A: $H_2O$ + 0.1% (v/v) TFA
Solvent B: MeCN + 0.1% (v/v) TFA

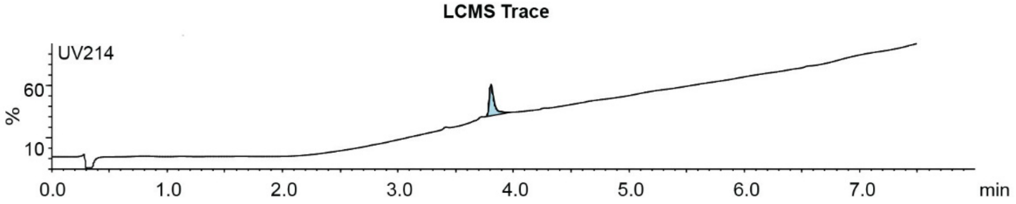

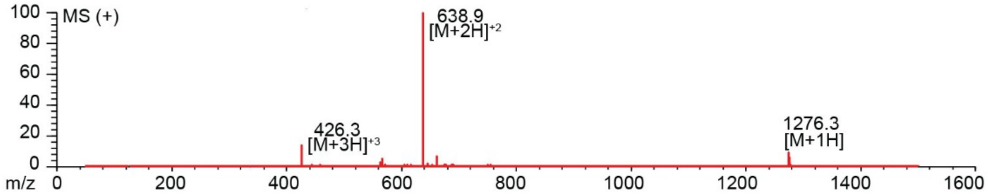

**Figure S2.8.** Synthesis summary of AECISFCRN-**PPA** (**8c**).
Expected: [M+1] = 1276.5277, $[M+2]^{+2}$ = 638.7638, $[M+3]^{+3}$ = 426.1759. Observed: [M+1] = 1276.3, $[M+2]^{+2}$ = 638.9, $[M+3]^{+3}$ = 426.3. Note: Two regioisomeric forms of the pyrazole are possible; the structure shown represents one isomer.

#### ARCGMVCNQ-DKL

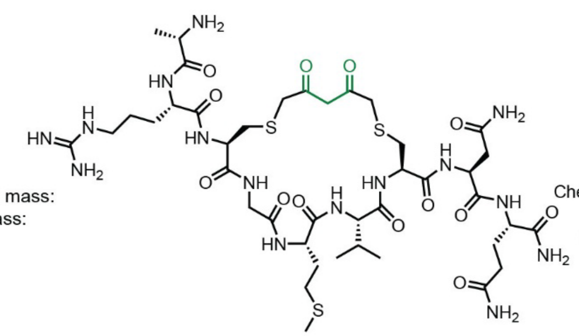

Starting material mass:
Final product mass:

Chemical Formula: $C_{41}H_{69}N_{15}O_{13}S_3$
Exact Mass: 1075.4361
Molecular Weight: 1076.2750

**HPLC Purification Trace**

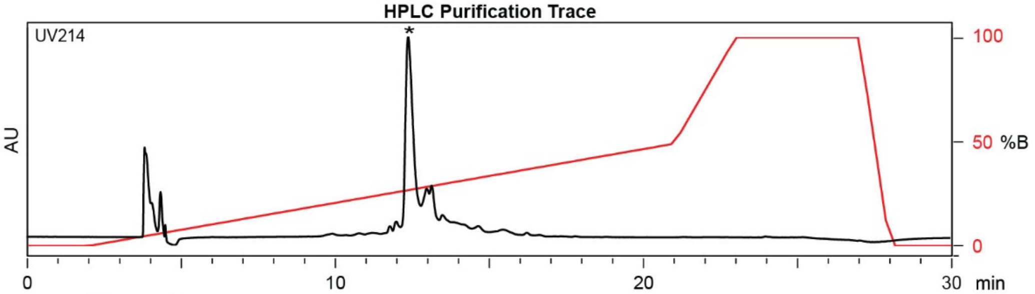

| Time (min) | Solvent B (%) |
|---|---|
| 0 | 2 |
| 2 | 2 |
| 22 | 50 |
| 24 | 100 |
| 27 | 100 |
| 28 | 2 |
| 30 | 2 |

Flow Rate: 13 mL / min
Phenomenex Kinetex EVO C18 Prep Column
(100 Å, 5 µm, 21.5 mm X 250 mm)

Solvent A: $H_2O$ + 0.1% (v/v) TFA
Solvent B: MeCN + 0.1% (v/v) TFA

**LCMS Trace**

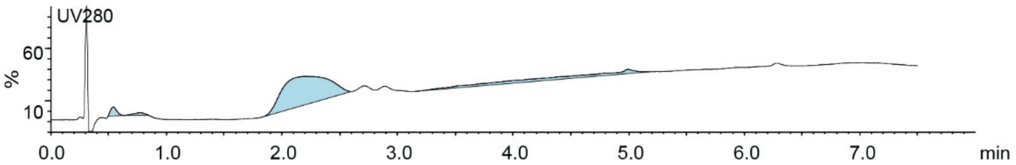

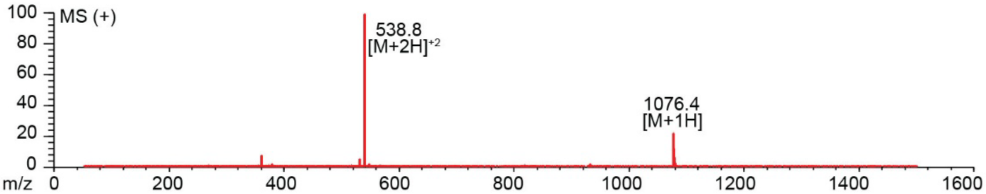

**Figure S2.9.** Synthesis summary of ARCGMVCNQ-**DKL** (**9b**).
Expected: [M+1] = 1076.4361, $[M+2]^{+2}$ = 538.7180. Observed: [M+1] = 1076.4, $[M+2]^{+2}$ = 538.8.

#### ARCGMVCNQ-PPA

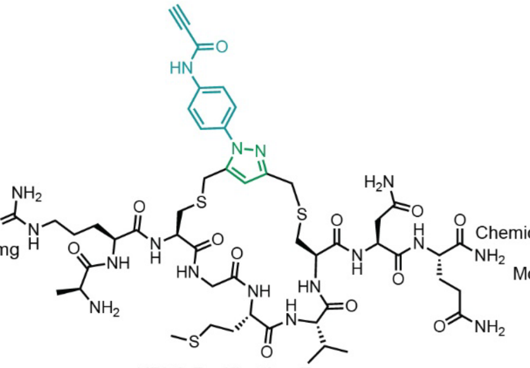

Starting material mass: 4.1 mg
Final product mass: 1.8 mg

Chemical Formula: $C_{50}H_{74}N_{18}O_{12}S_3$
Exact Mass: 1214.49
Molecular Weight: 1215.44

**HPLC Purification Trace**

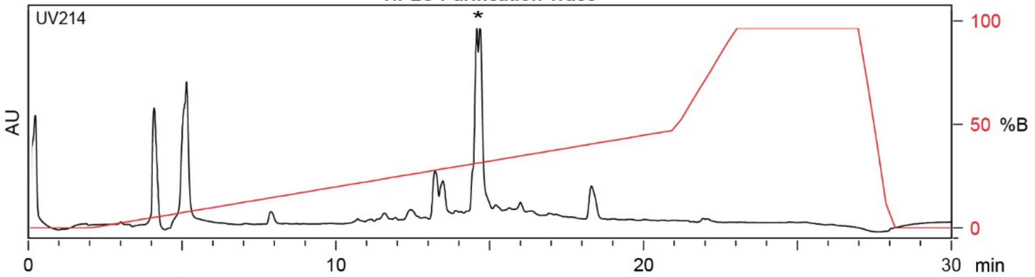

| Time (min) | Solvent B (%) |
|---|---|
| 0 | 2 |
| 2 | 2 |
| 21 | 50 |
| 24 | 100 |
| 27 | 100 |
| 29.5 | 2 |
| 30 | 2 |

Flow Rate: 13 mL / min
Phenomenex Kinetex EVO C18 Prep Column
(100 Å, 5 µm, 21.5 mm X 250 mm)

Solvent A: $H_2O$ + 0.1% (v/v) TFA
Solvent B: MeCN + 0.1% (v/v) TFA

**LCMS Trace**

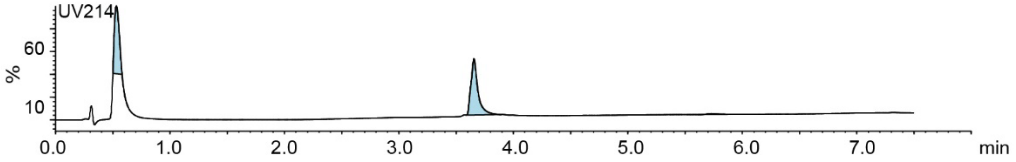

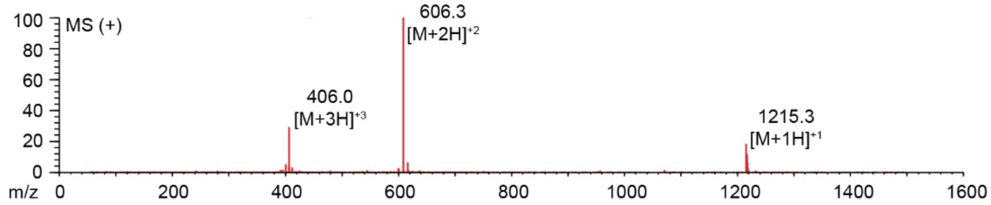

**Figure S2.10.** Synthesis summary of ARCGMVCNQ-**PPA** (**9c**).
Expected: [M+1] = 1215.49, $[M+2]^{+2}$ = 608.24, $[M+3]^{+3}$ = 405.83. Observed: [M+1] = 1215.3, $[M+2]^{+2}$ = 606.3, $[M+3]^{+3}$ = 406.0. Note: Two regioisomeric forms of the pyrazole are possible; the structure shown represents one isomer.

#### AWCGVRTC-DKL

Starting material mass:
Final product mass:

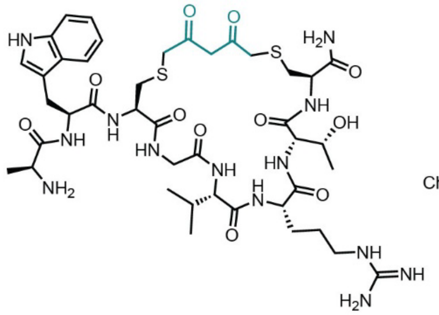

Chemical Formula: $C_{42}H_{63}N_{13}O_{11}S_2$
Exact Mass: 989.42
Molecular Weight: 990.17

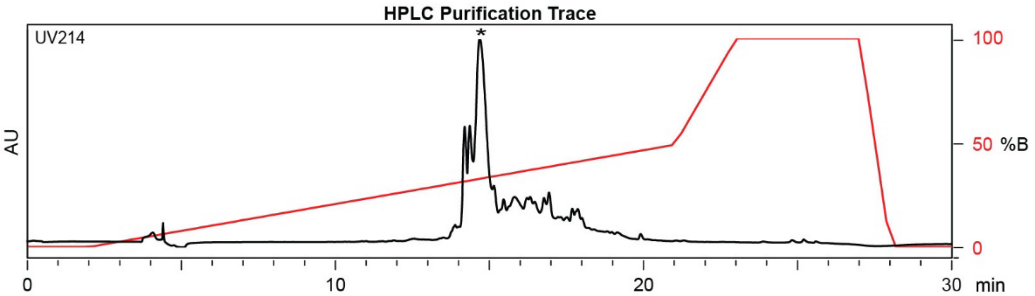

| Time (min) | Solvent B (%) |
|---|---|
| 0 | 2 |
| 2 | 2 |
| 22 | 50 |
| 24 | 100 |
| 27 | 100 |
| 28 | 2 |
| 30 | 2 |

Flow Rate: 13 mL / min
Phenomenex Kinetex EVO C18 Prep Column
(100 Å, 5 µm, 21.5 mm X 250 mm)

Solvent A: $H_2O$ + 0.1% (v/v) TFA
Solvent B: MeCN + 0.1% (v/v) TFA

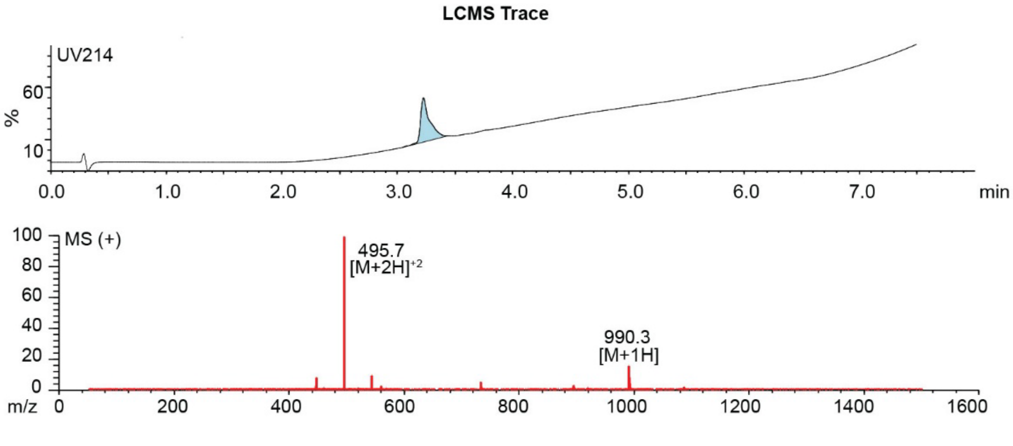

**Figure S2.11.** Synthesis summary of AWCGVRTC-**DKL** (**10b**).
Expected: [M+1] = 990.42, $[M+2]^{+2}$ = 495.71. Observed: [M+1] = 990.3, $[M+2]^{+2}$ = 495.7.

#### AWCGVRTC-PPA

Starting material mass: 3.5 mg
Final product mass: 1.2 mg

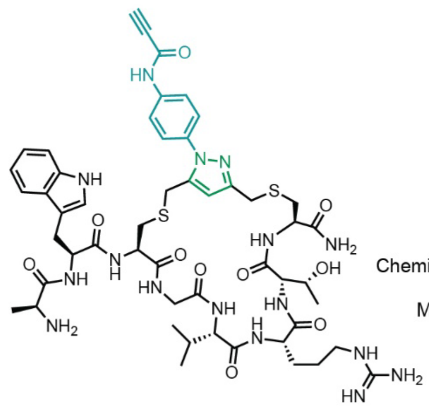

Chemical Formula: $C_{51}H_{68}N_{16}O_{10}S_2$
Exact Mass: 1128.47
Molecular Weight: 1129.33

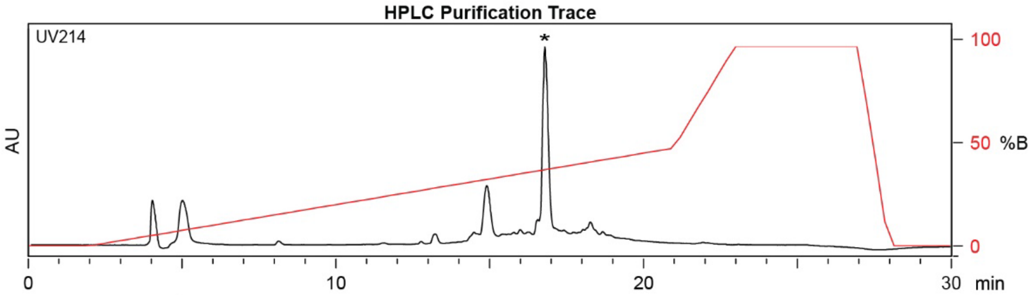

| Time (min) | Solvent B (%) |
|---|---|
| 0 | 2 |
| 2 | 2 |
| 21 | 50 |
| 24 | 100 |
| 27 | 100 |
| 29.5 | 2 |
| 30 | 2 |

Flow Rate: 13 mL / min
Phenomenex Kinetex EVO C18 Prep Column
(100 Å, 5 µm, 21.5 mm X 250 mm)

Solvent A: $H_2O$ + 0.1% (v/v) TFA
Solvent B: MeCN + 0.1% (v/v) TFA

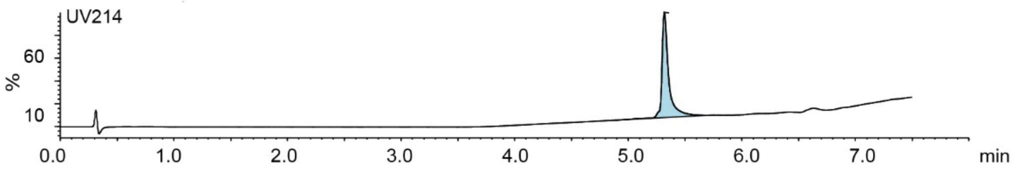

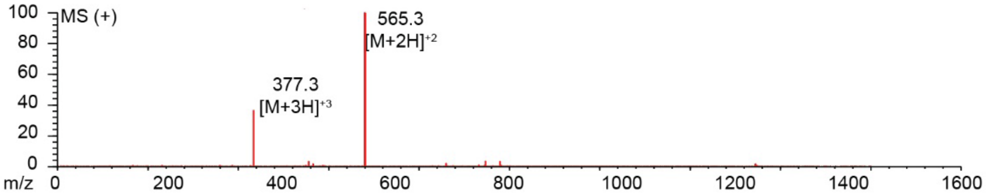

**Figure S2.12.** Synthesis summary of AWCGVRTC-**PPA** (**10c**).
Expected: [M+1] = 1129.47, $[M+2]^{\div 2}$ = 565.23, $[M+3]^{\div 3}$ = 377.16. Observed: $[M+2]^{\div 2}$ = 565.3, $[M+3]^{\div 3}$ =377.3. Note: Two regioisomeric forms of the pyrazole are possible; the structure shown represents one isomer.

#### ASCFFNTCH-DKL

Starting material mass: mg
Final product mass: mg

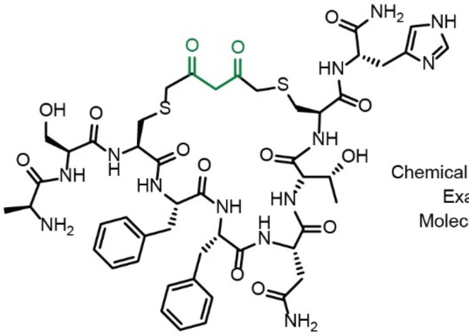

Chemical Formula: $C_{49}H_{65}N_{13}O_{14}S_2$
Exact Mass: 1123.4215
Molecular Weight: 1124.2560

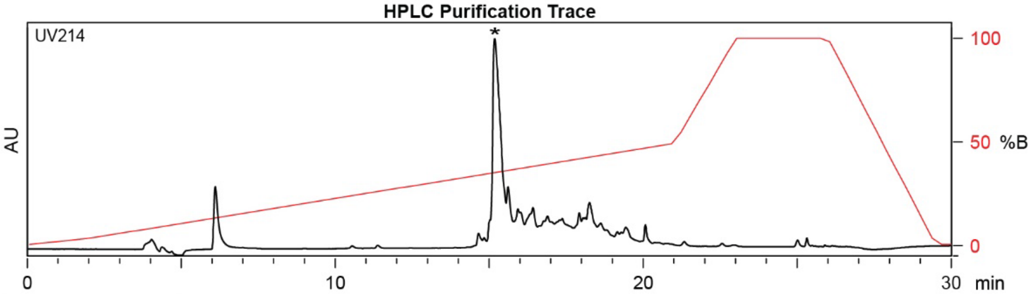

| Time (min) | Solvent B (%) |
|---|---|
| 0 | 2 |
| 2 | 2 |
| 22 | 50 |
| 24 | 100 |
| 27 | 100 |
| 28 | 2 |
| 30 | 2 |

Flow Rate: 13 mL / min
Phenomenex Kinetex EVO C18 Prep Column
(100 Å, 5 µm, 21.5 mm X 250 mm)

Solvent A: $H_2O$ + 0.1% (v/v) TFA
Solvent B: MeCN + 0.1% (v/v) TFA

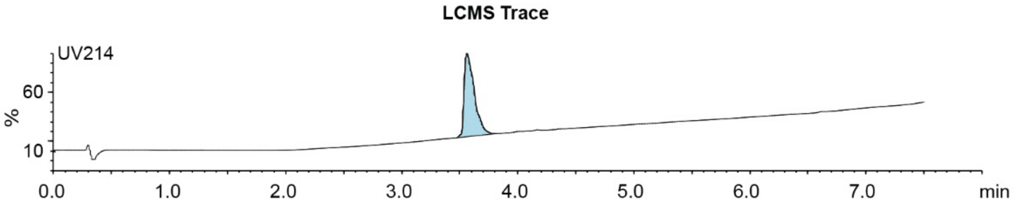

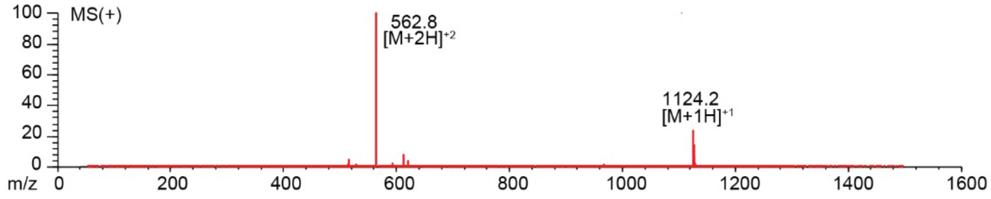

**Figure S2.13.** Synthesis summary of ASCFFNTCH-**DKL** (**11b**).
Expected: [M+1] = 1124.4215, $[M+2]^{+2}$ = 562.7108. Observed: [M+1] = 1124.2, $[M+2]^{+2}$ = 562.8.

#### ASCFFNTCH-PPA

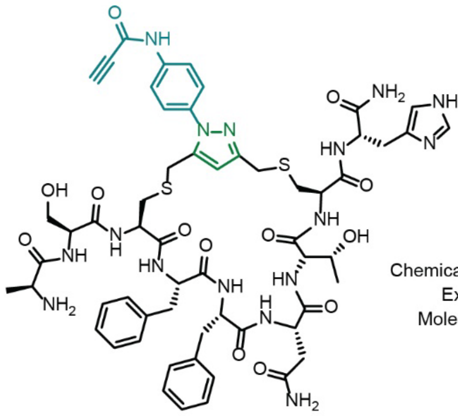

Starting material mass: mg
Final product mass: mg

Chemical Formula: $C_{58}H_{70}N_{16}O_{13}S_2$
Exact Mass: 1262.4750
Molecular Weight: 1263.4170

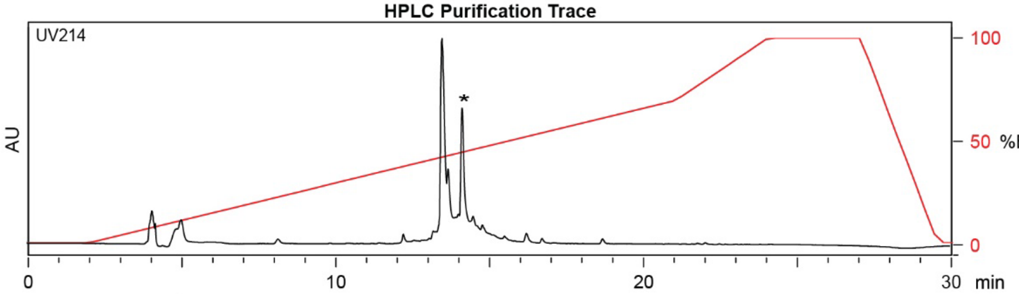

| Time (min) | Solvent B (%) |
|---|---|
| 0 | 2 |
| 2 | 2 |
| 21 | 70 |
| 24 | 100 |
| 27 | 100 |
| 29.5 | 2 |
| 30 | 2 |

Flow Rate: 13 mL / min
Phenomenex Kinetex EVO C18 Prep Column
(100 Å, 5 µm, 21.5 mm X 250 mm)

Solvent A: $H_2O$ + 0.1% (v/v) TFA
Solvent B: MeCN + 0.1% (v/v) TFA

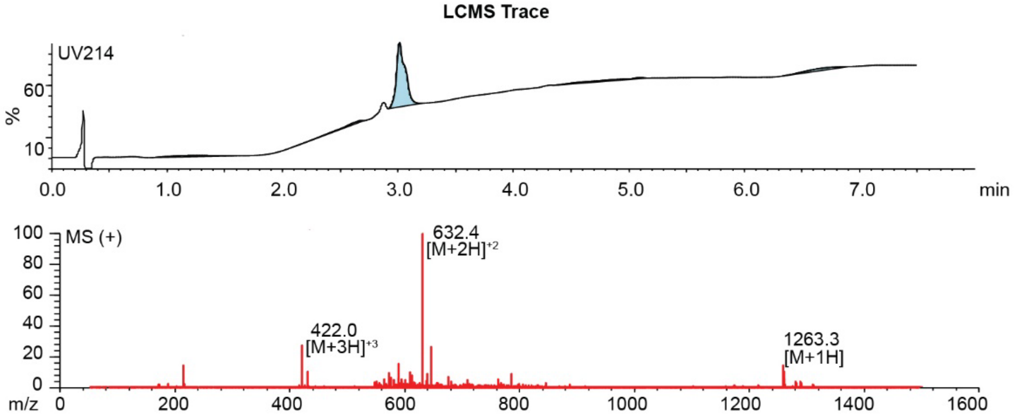

**Figure S2.14.** Synthesis summary of ASCFFNTCH-**PPA** (**11c**).
Expected: [M+1] = 1263.4750, $[M+2]^{+2}$ = 632.2375, $[M+3]^{+3}$ = 421.8250. Observed: [M+1] = 1263.3, $[M+2]^{+2}$ = 632.4, $[M+3]^{+3}$ = 422.0. Note: Two regioisomeric forms of the pyrazole are possible; the structure shown represents one isomer.

#### AFCSYDTCF-DKL

Starting material mass: mg
Final product mass: mg

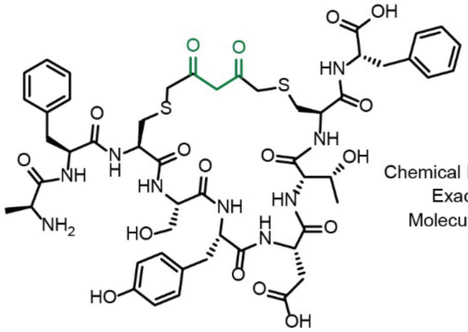

Chemical Formula: $C_{52}H_{65}N_{9}O_{17}S_{2}$
Exact Mass: 1151.3940
Molecular Weight: 1152.2580

**HPLC Purification Trace**

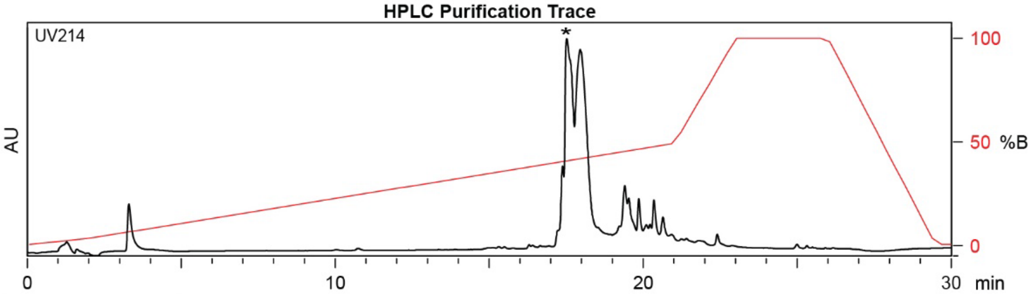

| Time (min) | Solvent B (%) |
|---|---|
| 0 | 2 |
| 2 | 2 |
| 22 | 50 |
| 24 | 100 |
| 27 | 100 |
| 28 | 2 |
| 30 | 2 |

Flow Rate: 13 mL / min
Phenomenex Kinetex EVO C18 Prep Column
(100 Å, 5 µm, 21.5 mm X 250 mm)

Solvent A: $H_2O$ + 0.1% (v/v) TFA
Solvent B: MeCN + 0.1% (v/v) TFA

**LCMS Trace**

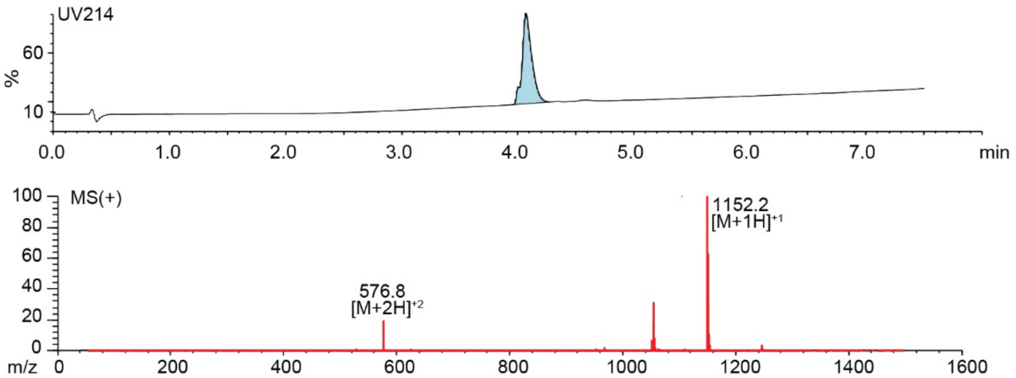

**Figure S2.15.** Synthesis summary of AFCSYDTCF-**DKL** (**12b**).
Expected: [M+1] = 1152.3940, $[M+2]^{+2}$ = 576.6970. Observed: [M+1] =1152.2 , $[M+2]^{+2}$ = 576.8.

#### AFCSYDTCF-PPA

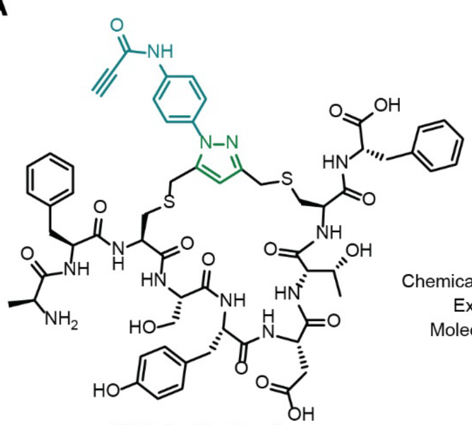

Starting material mass: mg
Final product mass: mg

Chemical Formula: $C_{61}H_{70}N_{12}O_{16}S_2$
Exact Mass: 1290.4474
Molecular Weight: 1291.4190

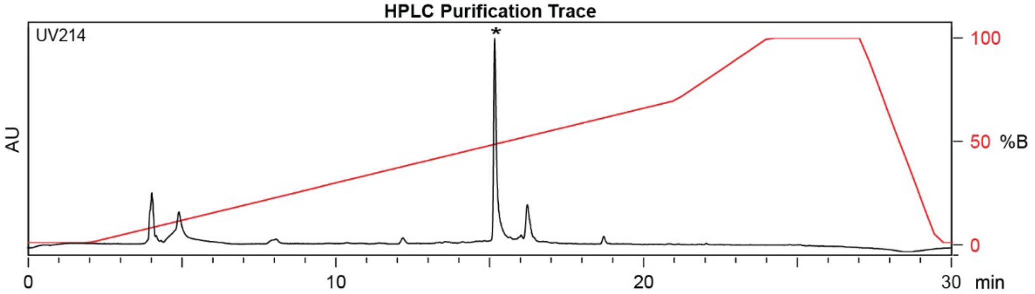

| Time (min) | Solvent B (%) |
|---|---|
| 0 | 2 |
| 2 | 2 |
| 21 | 70 |
| 24 | 100 |
| 27 | 100 |
| 29.5 | 2 |
| 30 | 2 |

Flow Rate: 13 mL / min
Phenomenex Kinetex EVO C18 Prep Column
(100 Å, 5 µm, 21.5 mm X 250 mm)

Solvent A: $H_2O$ + 0.1% (v/v) TFA
Solvent B: MeCN + 0.1% (v/v) TFA

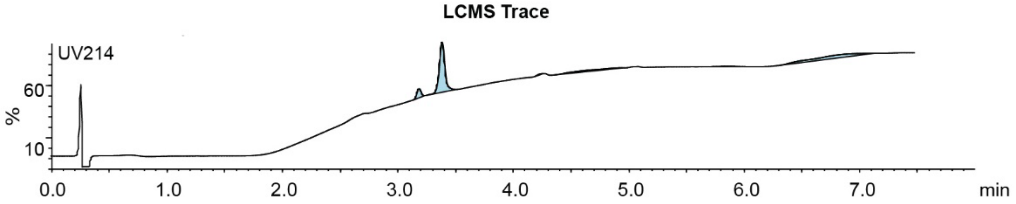

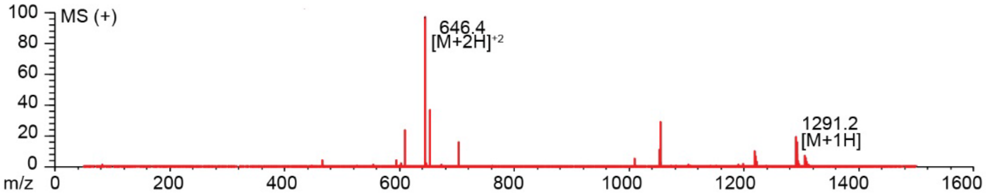

**Figure S2.16.** Synthesis summary of AFCSYDTCF-**PPA** (**12c**).
Expected: [M+1] = 1291.4474, $[M+2]^{+2}$ = 646.2237. Observed: [M+1] = 1291.2, $[M+2]^{+2}$ = 646.4.
Note: Two regioisomeric forms of the pyrazole are possible; the structure shown represents one plausible isomer.

#### ATCAWRNCT-DKL

Starting material mass: mg
Final product mass: mg

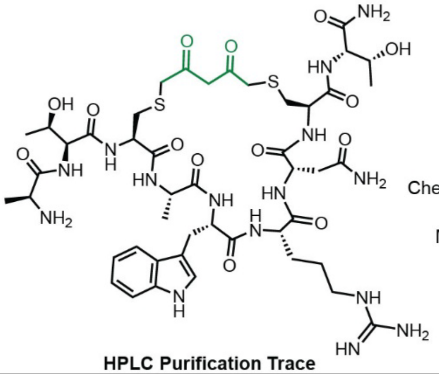

Chemical Formula: $C_{46}H_{69}N_{15}O_{14}S_2$
Exact Mass: 1119.4590
Molecular Weight: 1120.2690

**HPLC Purification Trace**

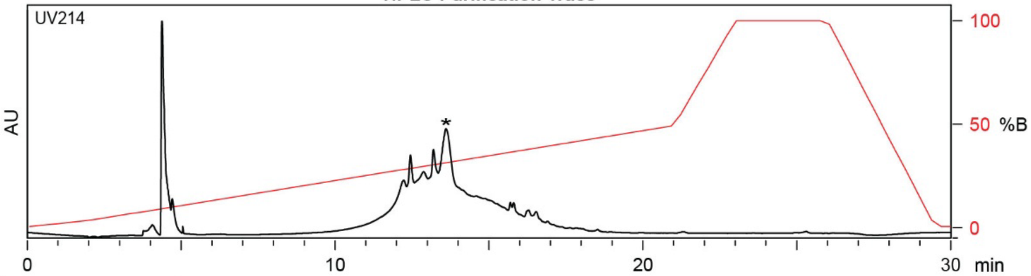

| Time (min) | Solvent B (%) |
|---|---|
| 0 | 2 |
| 2 | 2 |
| 22 | 50 |
| 24 | 100 |
| 27 | 100 |
| 28 | 2 |
| 30 | 2 |

Flow Rate: 13 mL / min
Phenomenex Kinetex EVO C18 Prep Column
(100 Å, 5 µm, 21.5 mm X 250 mm)

Solvent A: $H_2O$ + 0.1% (v/v) TFA
Solvent B: MeCN + 0.1% (v/v) TFA

**LCMS Trace**

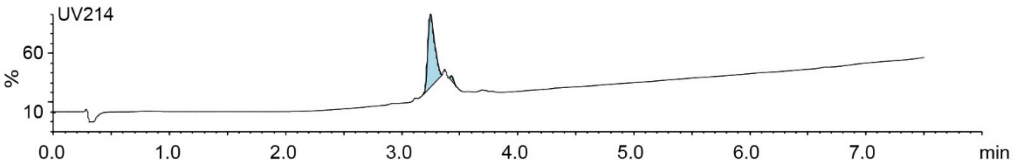

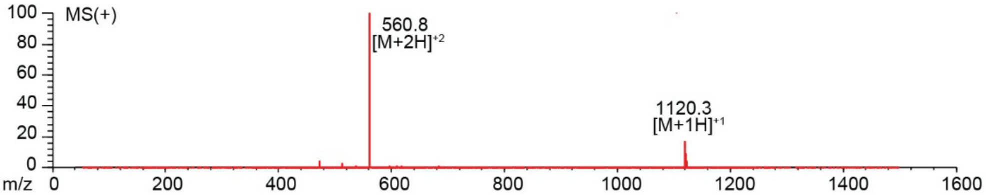

**Figure S2.17.** Synthesis summary of ATCAWRNCT-**DKL** (**13b**).
Expected: [M+1] = 1120.4590, $[M+2]^{\div 2}$ = 560.7295. Observed: [M+1] = 1120.3, $[M+2]^{\div 2}$ = 560.8.

#### ATCAWRNCT-PPA

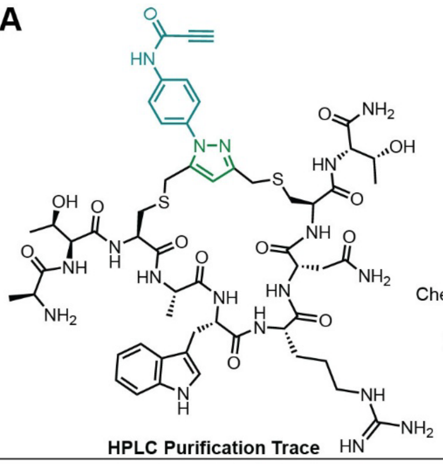

Starting material mass: mg
Final product mass: mg

Chemical Formula: $C_{55}H_{74}N_{18}O_{13}S_2$
Exact Mass: 1258.5124
Molecular Weight: 1259.4300

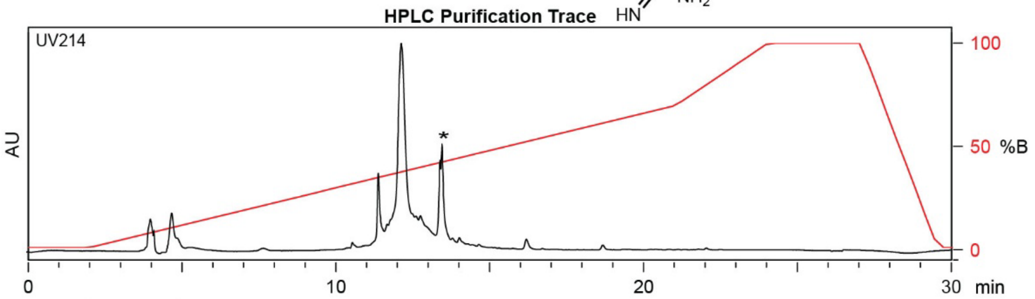

| Time (min) | Solvent B (%) |
|---|---|
| 0 | 2 |
| 2 | 2 |
| 21 | 70 |
| 24 | 100 |
| 27 | 100 |
| 29.5 | 2 |
| 30 | 2 |

Flow Rate: 13 mL / min
Phenomenex Kinetex EVO C18 Prep Column
(100 Å, 5 µm, 21.5 mm X 250 mm)

Solvent A: $H_2O$ + 0.1% (v/v) TFA
Solvent B: MeCN + 0.1% (v/v) TFA

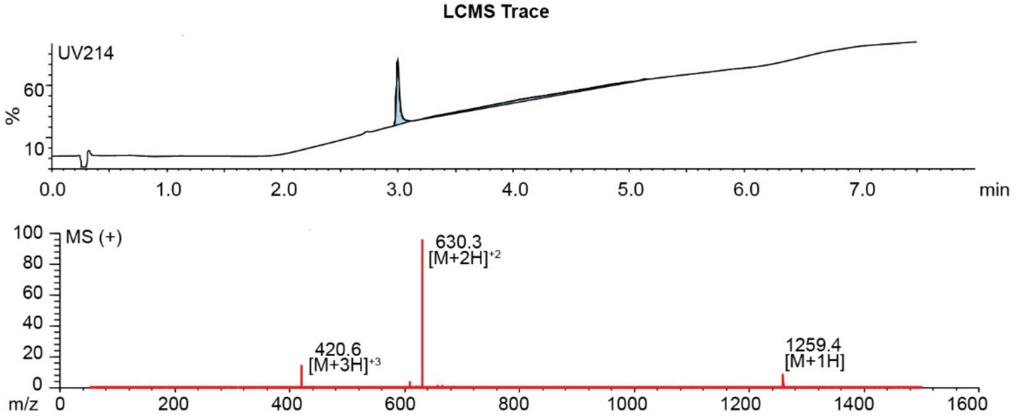

**Figure S2.18.** Synthesis summary of ATCAWRNCT-**PPA** (**13c**).

Expected: [M+1] = 1259.5124, $[M+2]^{+2}$ = 630.2562, $[M+3]^{+3}$ = 420.5041. Observed: [M+1] = 1259.4, $[M+2]^{+2}$ = 630.3, $[M+3]^{+3}$ = 420.6. Note: Two regioisomeric forms of the pyrazole are possible; the structure shown represents one isomer.

### ASCFFSACK-DKL

Starting material mass:
Final product mass:

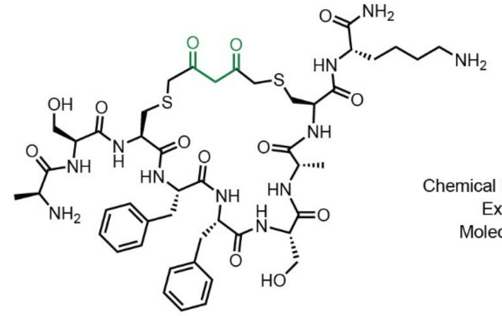

Chemical Formula: $C_{47}H_{67}N_{11}O_{13}S_2$
Exact Mass: 1057.44
Molecular Weight: 1058.24

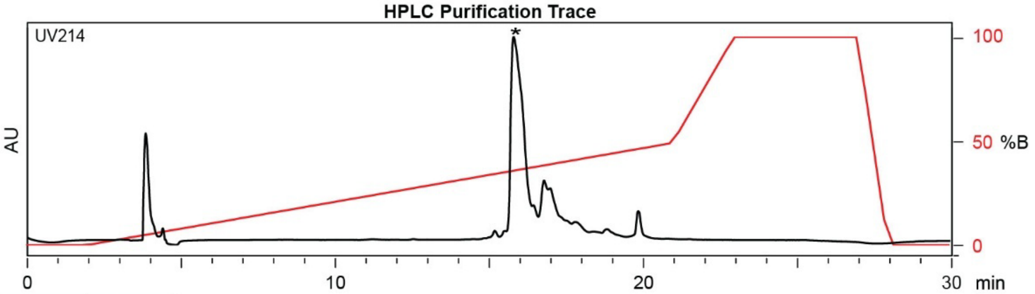

| Time (min) | Solvent B (%) |
|---|---|
| 0 | 2 |
| 2 | 2 |
| 22 | 50 |
| 24 | 100 |
| 27 | 100 |
| 28 | 2 |
| 30 | 2 |

Flow Rate: 13 mL / min
Phenomenex Kinetex EVO C18 Prep Column
(100 Å, 5 µm, 21.5 mm X 250 mm)

Solvent A: $H_2O$ + 0.1% (v/v) TFA
Solvent B: MeCN + 0.1% (v/v) TFA

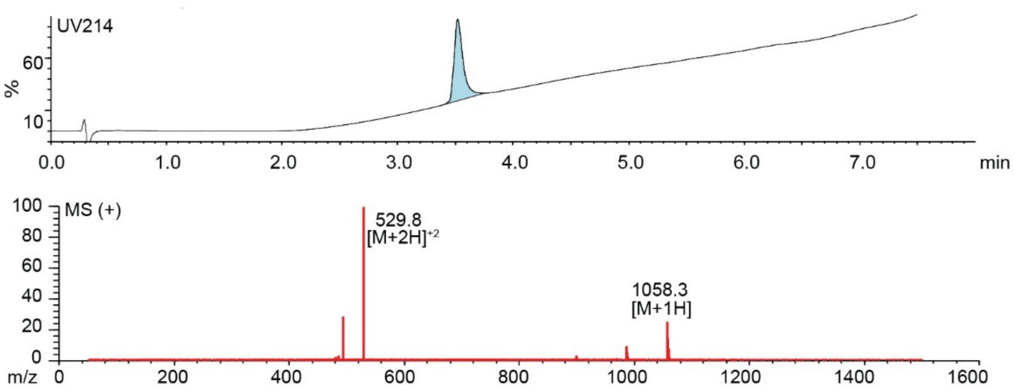

**Figure S2.19.** Synthesis summary of ASCFFSACK-**DKL** (**14b**).
Expected: [M+1] = 1058.44, $[M+2]^{+2}$ = 529.72. Observed: [M+1] = 1058.3, $[M+2]^{+2}$ = 529.8.

#### ASCFFSACK-PPA

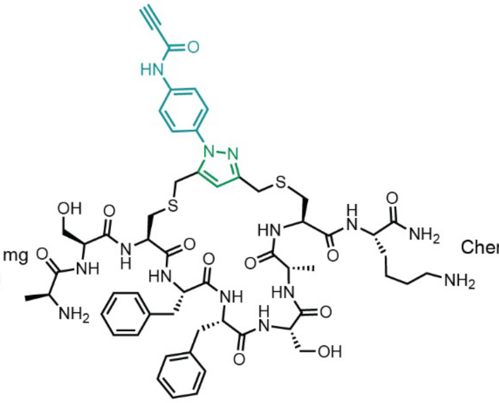

Starting material mass: 4.1 mg
Final product mass: 1.6 mg

Chemical Formula: $C_{56}H_{72}N_{14}O_{12}S_2$
Exact Mass: 1196.49
Molecular Weight: 1197.40

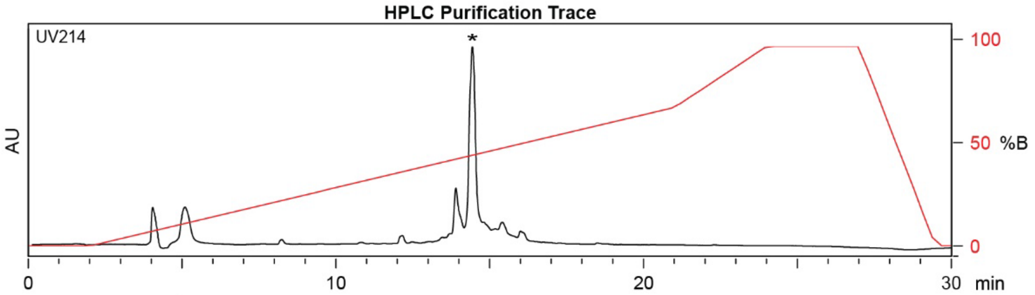

| Time (min) | Solvent B (%) |
|---|---|
| 0 | 2 |
| 2 | 2 |
| 21 | 50 |
| 24 | 100 |
| 27 | 100 |
| 29.5 | 2 |
| 30 | 2 |

Flow Rate: 13 mL / min
Phenomenex Kinetex EVO C18 Prep Column
(100 Å, 5 µm, 21.5 mm X 250 mm)

Solvent A: $H_2O$ + 0.1% (v/v) TFA
Solvent B: MeCN + 0.1% (v/v) TFA

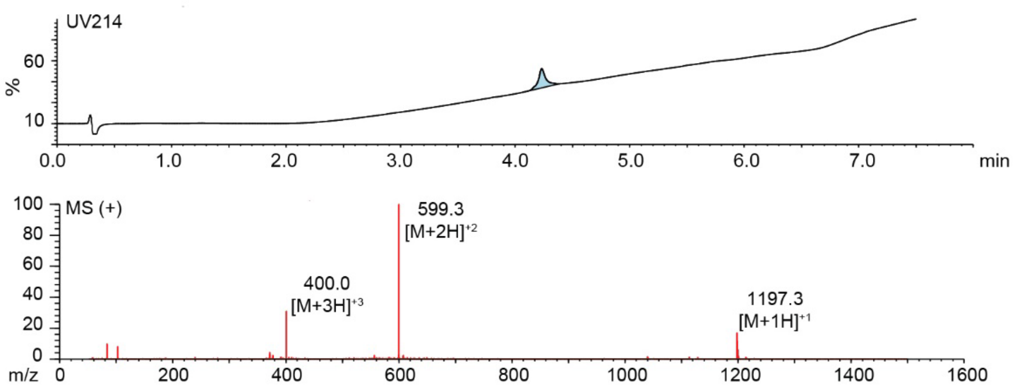

**Figure S2.20.** Synthesis summary of ASCFFSACK-**PPA** (**14c**).
Expected: [M+1] = 1197.49, $[M+2]^{+2}$ = 599.24, $[M+3]^{+3}$ = 399.83. Observed: [M+1] = 1197.3, $[M+2]^{+2}$ = 599.3, $[M+3]^{+3}$ = 400.0. Note: Two regioisomeric forms of the pyrazole are possible; the structure shown represents one isomer.

#### ANCPNYKCR

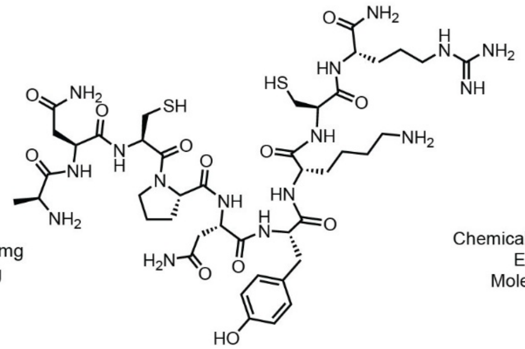

Starting material mass: N/A mg
Final product mass: 87.6 mg

Chemical Formula: $C_{43}H_{70}N_{16}O_{12}S_2$
Exact Mass: 1066.48
Molecular Weight: 1067.25

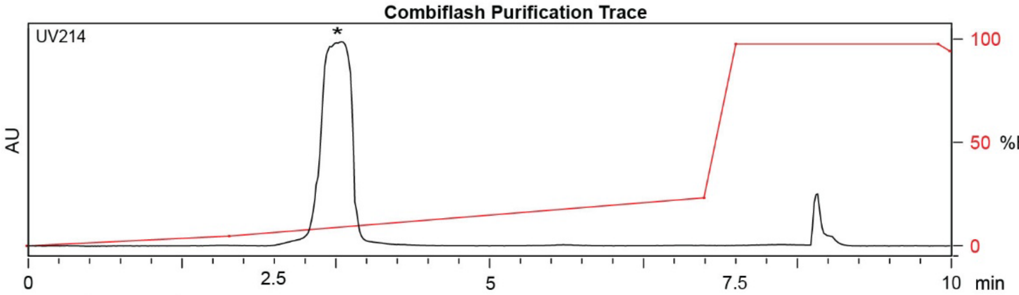

| Time (min) | Solvent B (%) |
| --- | --- |
| 0 | 0 |
| 1 | 5 |
| 5 | 20 |
| 7 | 25 |
| 7.5 | 100 |
| 9.8 | 100 |
| 100 | 95 |

Flow Rate: 35 mL / min
Redisep Rf Gold C18 Reverse Phase Column 30g
(100 Å, 20-40 µm, 300 ± 50 m²/g)

Solvent A: $H_2O$ + 0.1% (v/v) TFA
Solvent B: MeCN + 0.1% (v/v) TFA

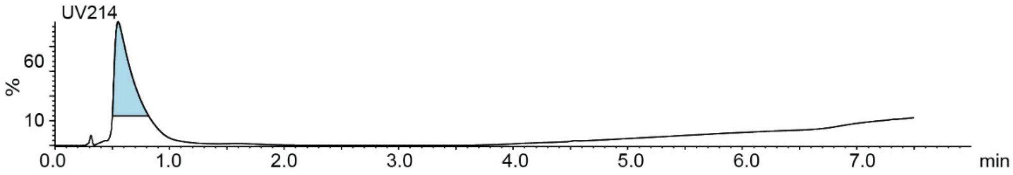

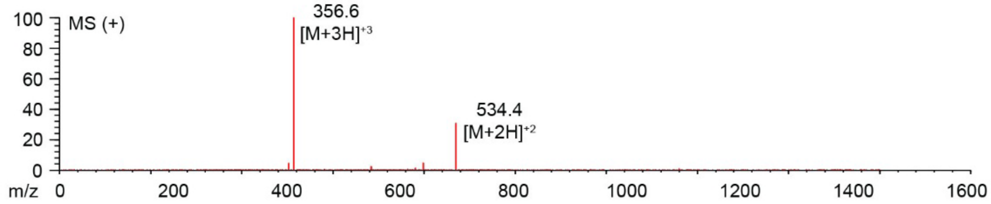

**Figure S2.21.** Synthesis summary of ANCPNYKCR (**15a**).
Expected: [M+1] = 1067.48, $[M+2]^{\div 2}$ = 534.24, $[M+3]^{\div 3}$ = 356.49. Observed: $[M+2]^{\div 2}$ = 534.4, $[M+3]^{\div 3}$ = 356.6.

#### ANCPNYKCR-DKL

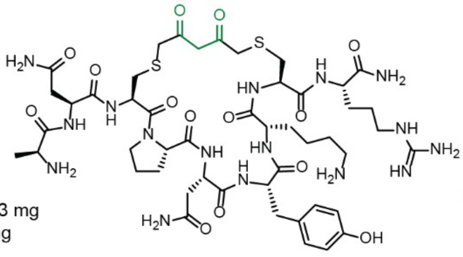

Starting material mass: 5.3 mg
Final product mass: 2.5 mg

Chemical Formula: $C_{48}H_{74}N_{16}O_{14}S_2$
Exact Mass: 1162.5012
Molecular Weight: 1163.3380

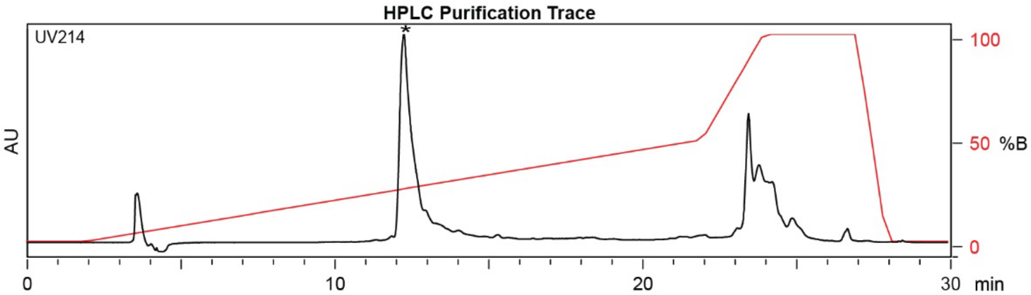

| Time (min) | Solvent B (%) |
|---|---|
| 0 | 2 |
| 2 | 2 |
| 22 | 50 |
| 24 | 100 |
| 27 | 100 |
| 28 | 2 |
| 30 | 2 |

Flow Rate: 13 mL / min
Phenomenex Kinetex EVO C18 Prep Column
(100 Å, 5 µm, 21.5 mm X 250 mm)

Solvent A: $H_2O$ + 0.1% (v/v) TFA
Solvent B: MeCN + 0.1% (v/v) TFA

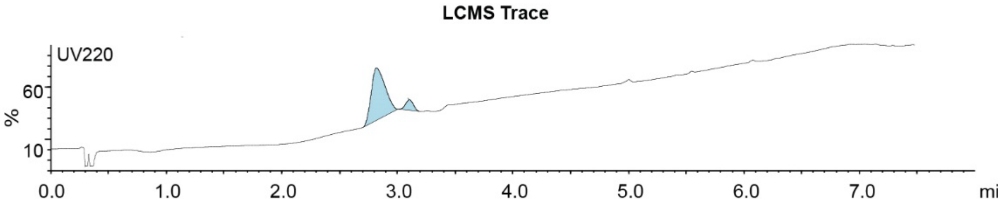

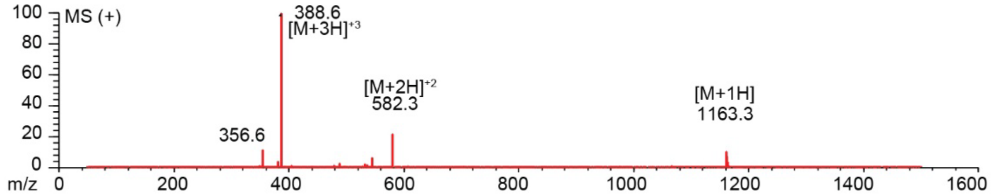

**Figure S2.22.** Synthesis summary of ANCPNYKCR -**DKL** (**15b**).
Expected: [M+1] = 1163.5012, $[M+2]^{+2}$ = 582.2506, $[M+3]^{+3}$ = 388.5004. Observed: [M+1] = 1163.3, $[M+2]^{+2}$ =582.3 , $[M+3]^{+3}$ = 388.6.

#### ANCPNYKCR-PPA

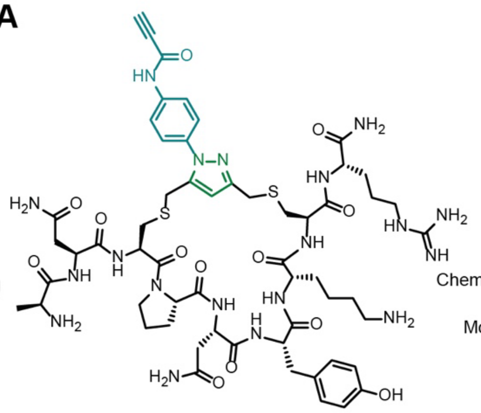

Starting material mass: 3.4 mg
Final product mass: 1.6 mg

Chemical Formula: $C_{57}H_{79}N_{19}O_{13}S_2$
Exact Mass: 1301.5546
Molecular Weight: 1302.4990

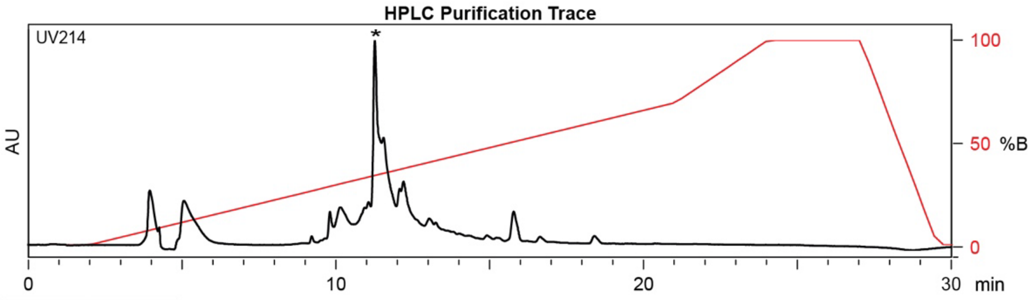

| Time (min) | Solvent B (%) |
|---|---|
| 0 | 2 |
| 2 | 2 |
| 21 | 70 |
| 24 | 100 |
| 27 | 100 |
| 29.5 | 2 |
| 30 | 2 |

Flow Rate: 13 mL / min
Phenomenex Kinetex EVO C18 Prep Column
(100 Å, 5 µm, 21.5 mm X 250 mm)

Solvent A: $H_2O$ + 0.1% (v/v) TFA
Solvent B: MeCN + 0.1% (v/v) TFA

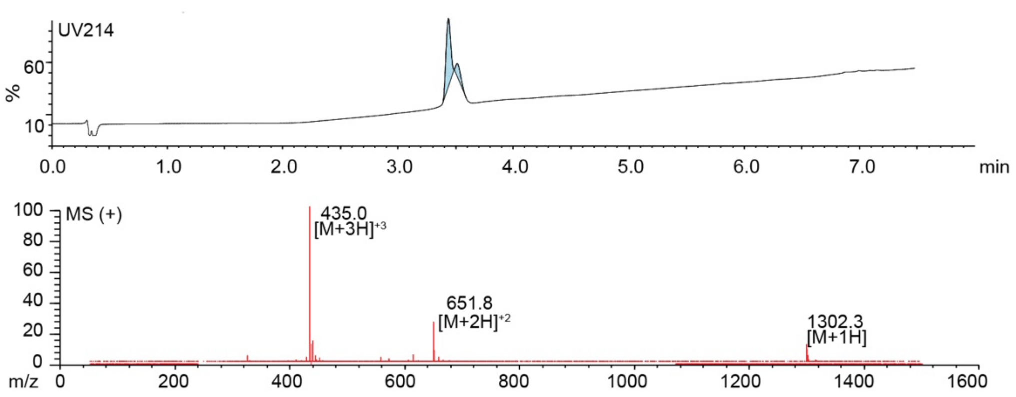

**Figure S2.23.** Synthesis summary of ANCPNYKCR-**PPA** (**15c**).
Expected: [M+1] = 1302.5546, $[M+2]^{+2}$ = 651.7773, $[M+3]^{+3}$ = 434.8515. Observed: [M+1] = 1302.3, $[M+2]^{+2}$ = 651.8, $[M+3]^{+3}$ = 435.0. Note: Two regioisomeric forms of the pyrazole are possible; the structure shown represents one isomer.

#### ANCPNYKCR-PEA

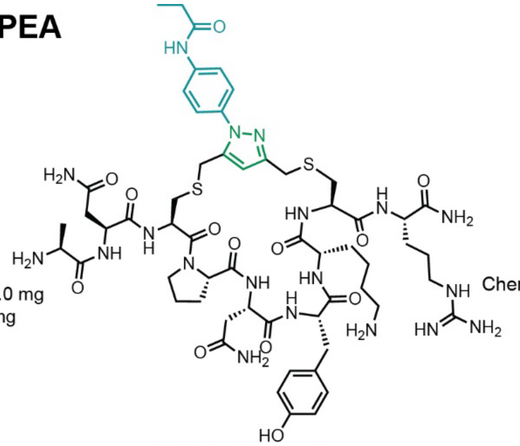

Starting material mass: 3.0 mg
Final product mass: 0.8 mg

Chemical Formula: $C_{57}H_{83}N_{19}O_{13}S_2$
Exact Mass: 1305.59
Molecular Weight: 1306.53

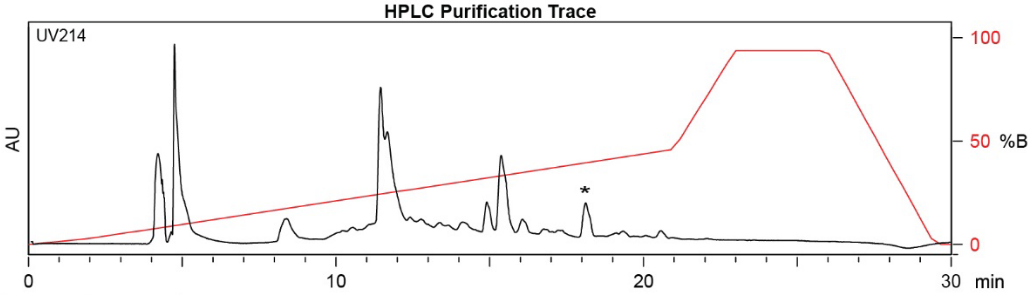

| Time (min) | Solvent B (%) |
|---|---|
| 0 | 2 |
| 2 | 2 |
| 21 | 50 |
| 24 | 100 |
| 27 | 100 |
| 29.5 | 2 |
| 30 | 2 |

Flow Rate: 13 mL / min
Phenomenex Kinetex EVO C18 Prep Column
(100 Å, 5 μm, 21.5 mm X 250 mm)

Solvent A: $H_2O$ + 0.1% (v/v) TFA
Solvent B: MeCN + 0.1% (v/v) TFA

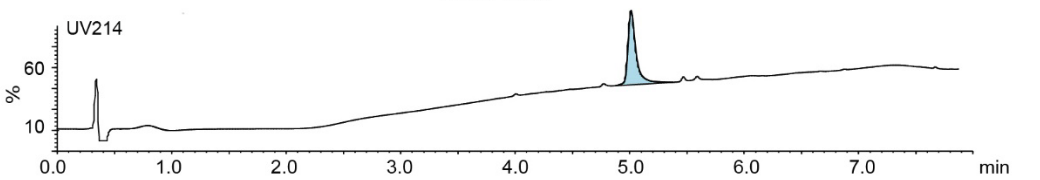

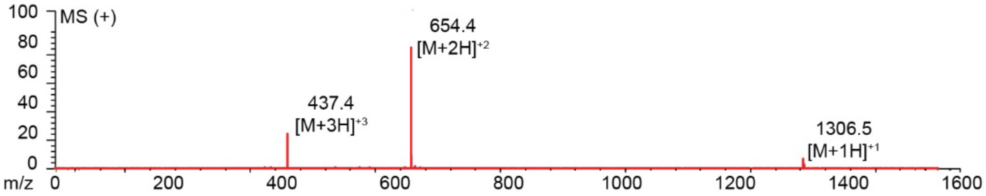

**Figure S2.24.** Synthesis summary of ANCPNYKCR-**PEA** (**15d**).
Expected: [M+1] = 1306.59, $[M+2]^{\div 2}$ = 653.80, $[M+3]^{\div 3}$ = 436.20. Observed: [M+1] = 1306.5, $[M+2]^{\div 2}$ = 654.4, $[M+3]^{\div 3}$ = 437.4. Note: Two regioisomeric forms of the pyrazole are possible; the structure shown represents one isomer.

#### ACHTSICWL-DKL

Starting material mass: mg
Final product mass: mg

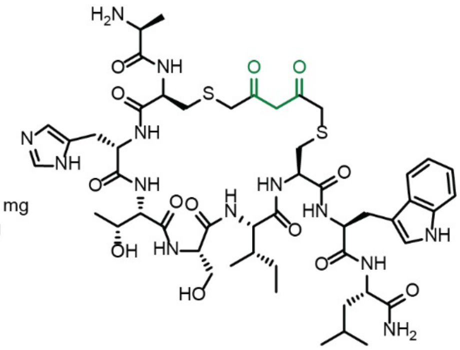

Chemical Formula: $C_{50}H_{73}N_{13}O_{13}S_2$
Exact Mass: 1127.4892
Molecular Weight: 1128.3320

HPLC Purification Trace

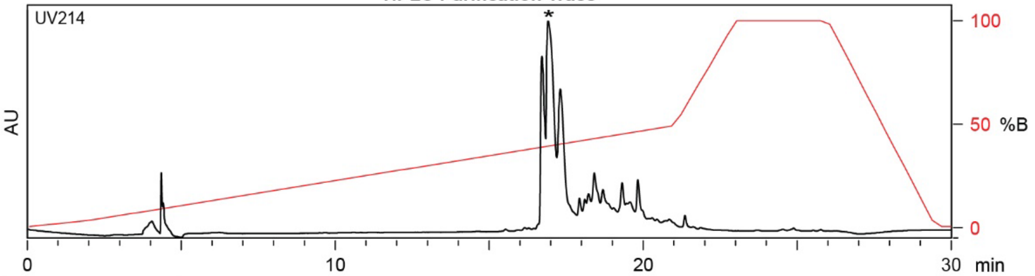

| Time (min) | Solvent B (%) |
|---|---|
| 0 | 2 |
| 2 | 2 |
| 22 | 50 |
| 24 | 100 |
| 27 | 100 |
| 28 | 2 |
| 30 | 2 |

Flow Rate: 13 mL / min
Phenomenex Kinetex EVO C18 Prep Column
(100 Å, 5 µm, 21.5 mm X 250 mm)

Solvent A: $H_2O$ + 0.1% (v/v) TFA
Solvent B: MeCN + 0.1% (v/v) TFA

LCMS Trace

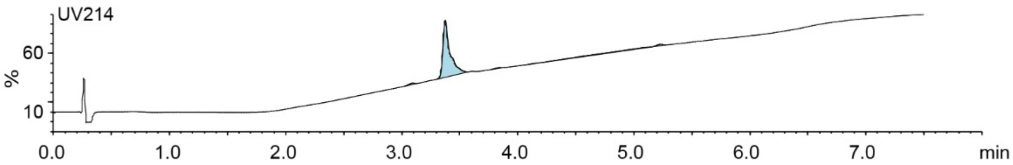

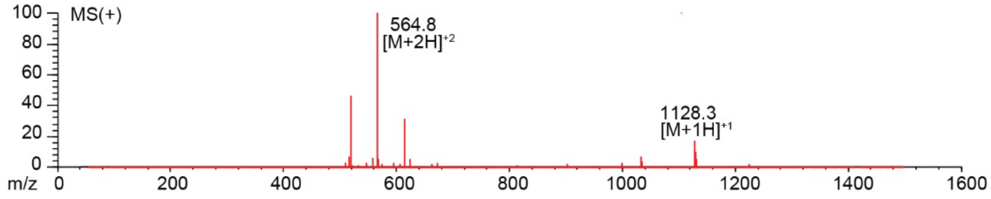

**Figure S2.25.** Synthesis summary of ACHTSICWL-**DKL** (**16b**).
Expected: [M+1] = 1128.4892, $[M+2]^{+2}$ = 564.7446. Observed: [M+1] = 1128.3, $[M+2]^{+2}$ = 564.8.

#### ACHTSICWL-PPA

Starting material mass: mg
Final product mass: mg

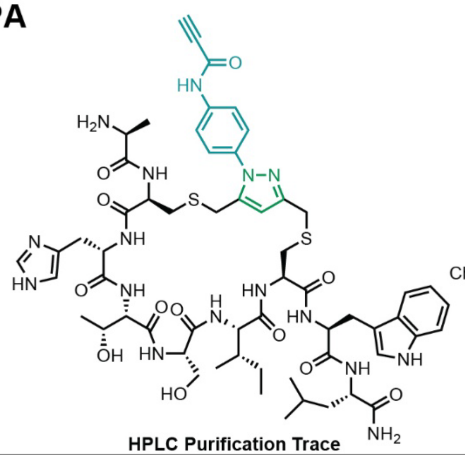

Chemical Formula: $C_{59}H_{78}N_{16}O_{12}S_2$
Exact Mass: 1266.54
Molecular Weight: 1267.49

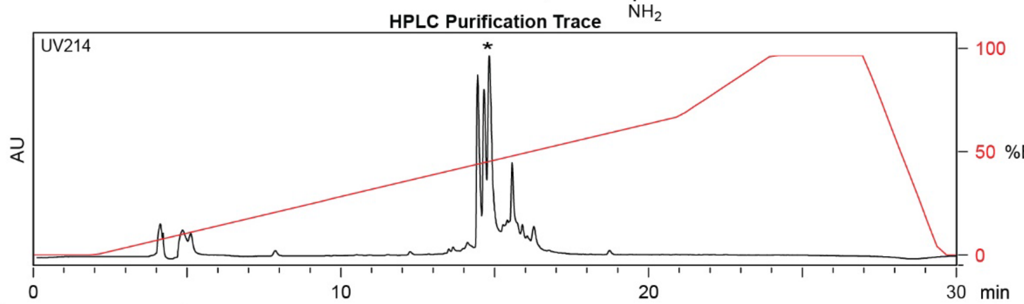

| Time (min) | Solvent B (%) |
|---|---|
| 0 | 2 |
| 2 | 2 |
| 21 | 70 |
| 24 | 100 |
| 27 | 100 |
| 29.5 | 2 |
| 30 | 2 |

Flow Rate: 13 mL / min
Phenomenex Kinetex EVO C18 Prep Column
(100 Å, 5 µm, 21.5 mm X 250 mm)

Solvent A: $H_2O$ + 0.1% (v/v) TFA
Solvent B: MeCN + 0.1% (v/v) TFA

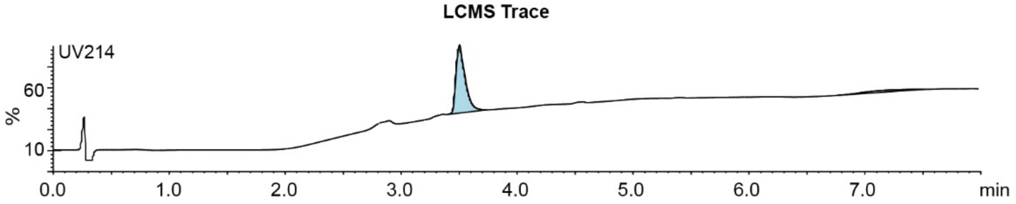

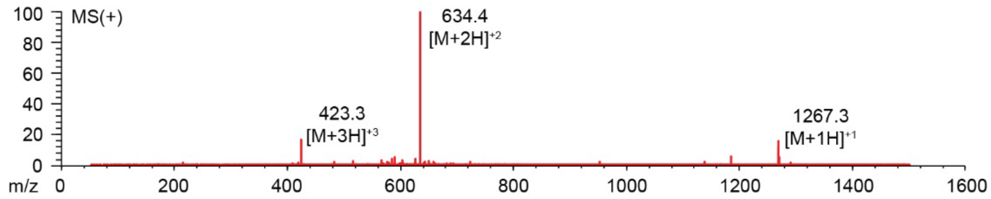

**Figure S2.26.** Synthesis summary of ACHTSICWL-**PPA** (**16c**).
Expected: [M+1] =1267.54 , $[M+2]^{+2}$ = 634.27, $[M+3]^{+3}$ = 423.18. Observed: [M+1] = 1267.3, $[M+2]^{+2}$ = 634.4, $[M+3]^{+3}$ = 423.3. Note: Two regioisomeric forms of the pyrazole are possible; the structure shown represents one isomer.

#### ACWKMNCLH-DKL

Starting material mass: mg
Final product mass: mg

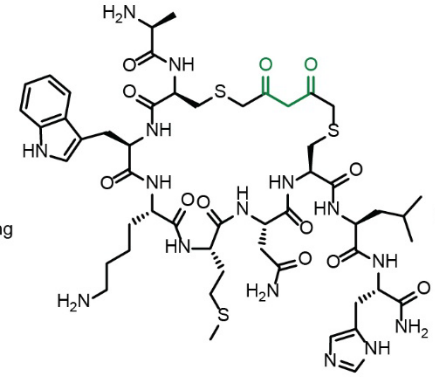

Chemical Formula: $C_{52}H_{77}N_{15}O_{12}S_3$
Exact Mass: 1199.5038
Molecular Weight: 1200.4610

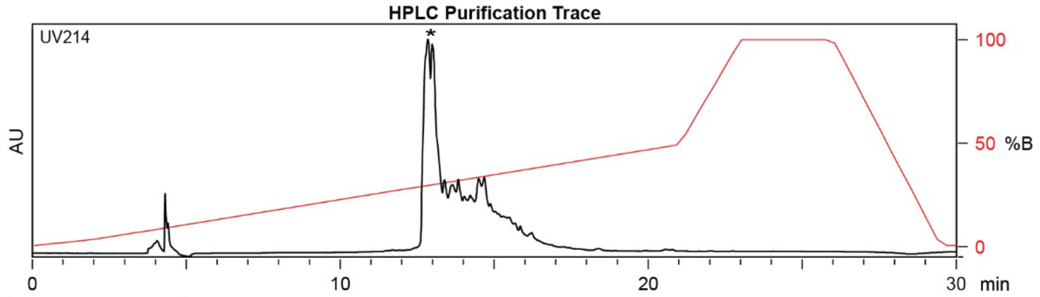

| Time (min) | Solvent B (%) |
|---|---|
| 0 | 2 |
| 2 | 2 |
| 22 | 50 |
| 24 | 100 |
| 27 | 100 |
| 28 | 2 |
| 30 | 2 |

Flow Rate: 13 mL / min
Phenomenex Kinetex EVO C18 Prep Column
(100 Å, 5 µm, 21.5 mm X 250 mm)

Solvent A: $H_2O$ + 0.1% (v/v) TFA
Solvent B: MeCN + 0.1% (v/v) TFA

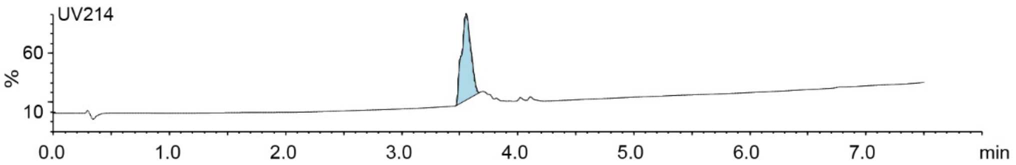

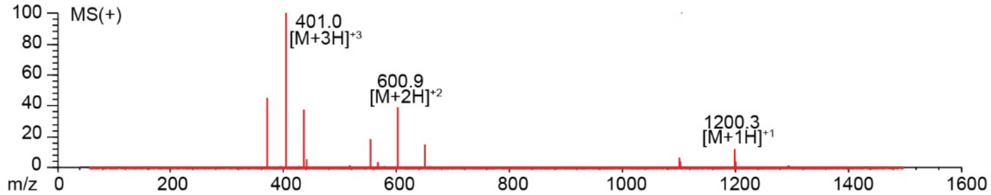

**Figure S2.27.** Synthesis summary of ACWKMNCLH-**DKL** (**17b**).
Expected: [M+1] = 1200.5038, $[M+2]^{+2}$ = 600.7519, $[M+3]^{+3}$ = 400.8346. Observed: [M+1] = 1200.3, $[M+2]^{+2}$ = 600.9, $[M+3]^{+3}$ = 401.0.

#### ACWKMNCLH-PPA

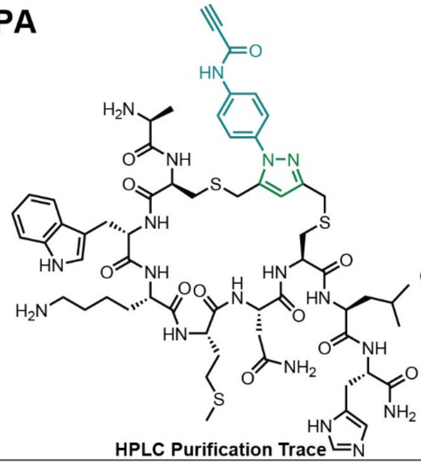

Starting material mass: mg
Final product mass: mg

Chemical Formula: $C_{61}H_{82}N_{18}O_{11}S_3$
Exact Mass: 1338.5573
Molecular Weight: 1339.6220

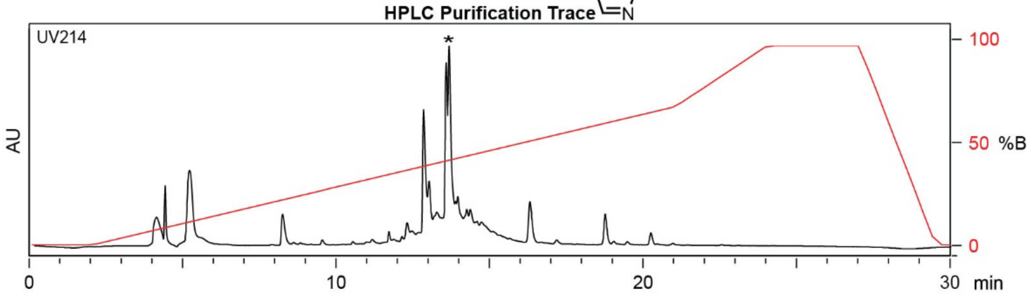

| Time (min) | Solvent B (%) |
|---|---|
| 0 | 2 |
| 2 | 2 |
| 21 | 70 |
| 24 | 100 |
| 27 | 100 |
| 29.5 | 2 |
| 30 | 2 |

Flow Rate: 13 mL / min
Phenomenex Kinetex EVO C18 Prep Column
(100 Å, 5 µm, 21.5 mm X 250 mm)

Solvent A: $H_2O$ + 0.1% (v/v) TFA
Solvent B: MeCN + 0.1% (v/v) TFA

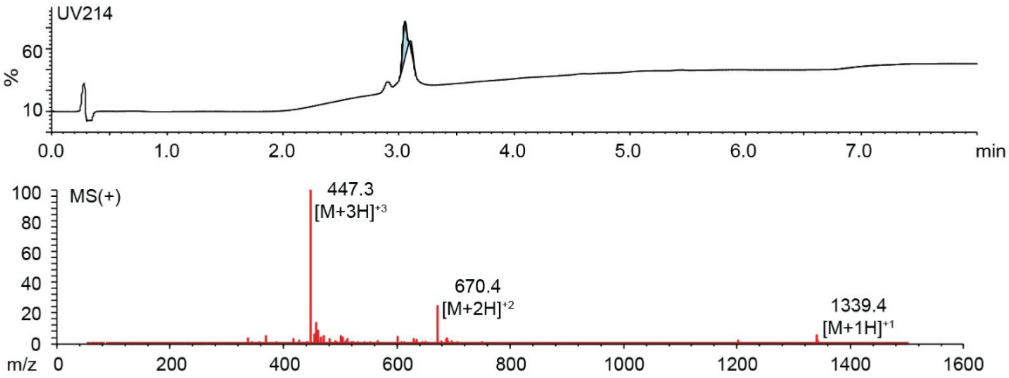

**Figure S2.28.** Synthesis summary of ACWKMNCLH-**PPA** (**17c**).
Expected: [M+1] = 1339.5573, $[M+2]^{+2}$ = 670.2786, $[M+3]^{+3}$ = 447.1858. Observed: [M+1] = 1339.4, $[M+2]^{+2}$ = 670.4, $[M+3]^{+3}$ = 447.3. Note: Two regioisomeric forms of the pyrazole are possible; the structure shown represents one isomer.

#### ACRGFLCNY

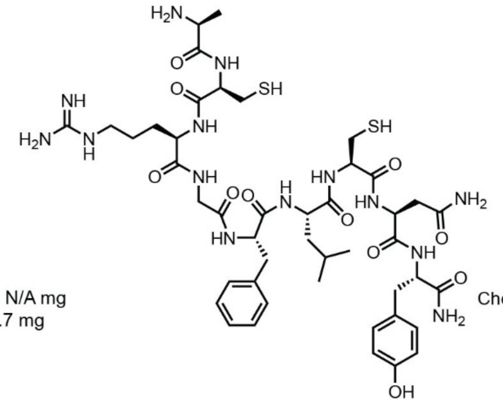

Starting material mass: N/A mg
Final product mass: 89.7 mg

Chemical Formula: $C_{45}H_{68}N_{14}O_{11}S_2$
Exact Mass: 1044.46
Molecular Weight: 1045.25

**Combiflash Purification Trace**

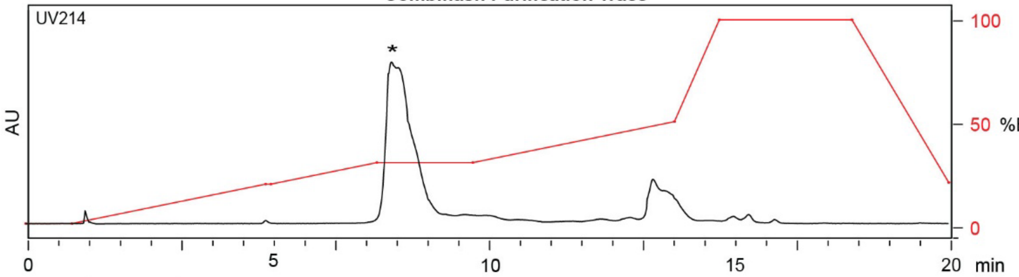

| Time (min) | Solvent B (%) |
|---|---|
| 0 | 0 |
| 1 | 0 |
| 5 | 25 |
| 9.5 | 25 |
| 13 | 50 |
| 15 | 100 |
| 20 | 25 |

Flow Rate: 35 mL / min
Redisep Rf Gold C18 Reverse Phase Column 30g
(100 Å, 20-40 μm, 300 ± 50 m²/g)

Solvent A: $H_2O$ + 0.1% (v/v) TFA
Solvent B: MeCN + 0.1% (v/v) TFA

**LCMS Trace**

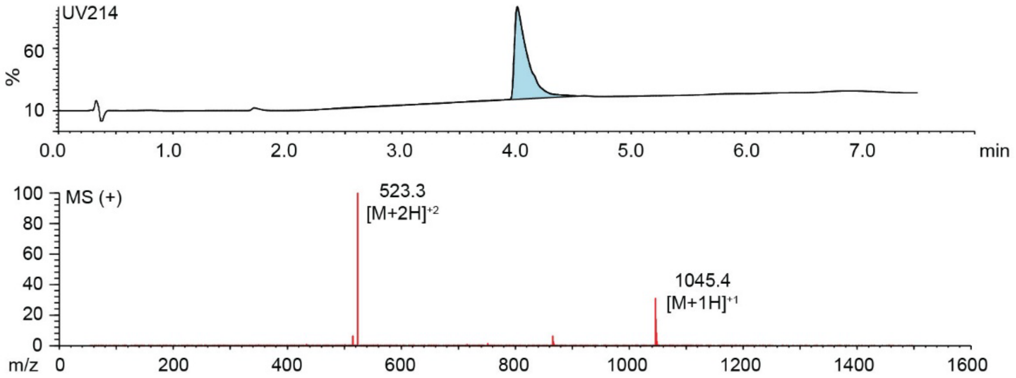

**Figure S2.29.** Synthesis summary of ACRGFLCNY (**18a**).
Expected: [M+1] = 1045.46, $[M+2]^{\div 2}$ = 523.23. Observed: [M+1] = 1045.4, $[M+2]^{\div 2}$ = 523.3.

#### ACRGFLCNY-DKL

Starting material mass:
Final product mass:

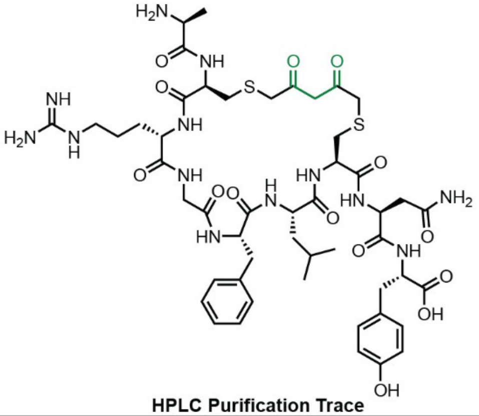

Chemical Formula: $C_{50}H_{71}N_{13}O_{14}S_2$
Exact Mass: 1141.4685
Molecular Weight: 1142.3150

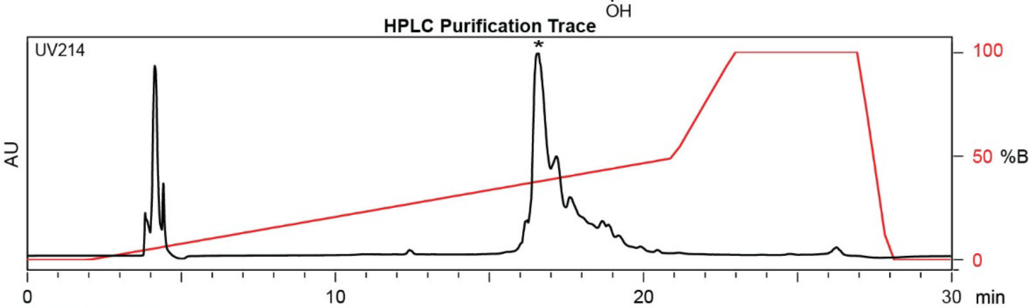

| Time (min) | Solvent B (%) |
|---|---|
| 0 | 2 |
| 2 | 2 |
| 22 | 50 |
| 24 | 100 |
| 27 | 100 |
| 28 | 2 |
| 30 | 2 |

Flow Rate: 13 mL / min
Phenomenex Kinetex EVO C18 Prep Column
(100 Å, 5 µm, 21.5 mm X 250 mm)

Solvent A: $H_2O$ + 0.1% (v/v) TFA
Solvent B: MeCN + 0.1% (v/v) TFA

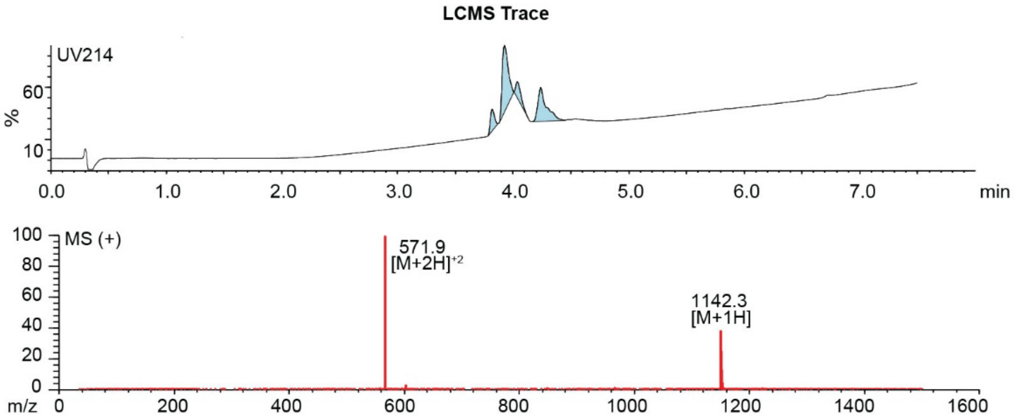

**Figure S2.30.** Synthesis summary of ACRGFLCNY-**DKL** (**18b**).
Expected: [M+1] = 1142.4685, $[M+2]^{+2}$ = 571.7342. Observed: [M+1] = 1142.3, $[M+2]^{+2}$ = 571.9.

**ARCGFLCNY-PPA**

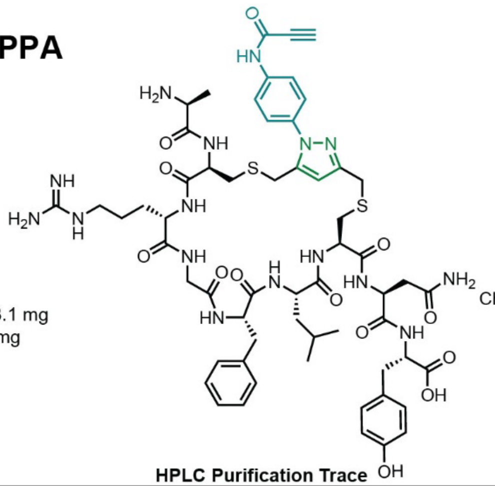

Starting material mass: 3.1 mg
Final product mass: 1.5 mg

Chemical Formula: $C_{59}H_{76}N_{16}O_{13}S_2$
Exact Mass: 1280.5219
Molecular Weight: 1281.4760

**HPLC Purification Trace**

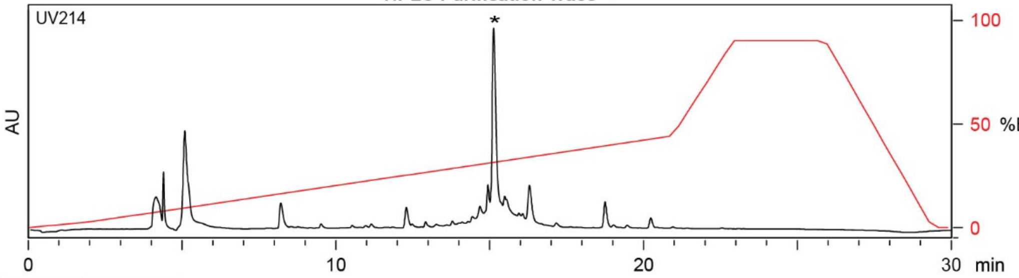

| Time (min) | Solvent B (%) |
|---|---|
| 0 | 2 |
| 2 | 2 |
| 21 | 50 |
| 24 | 100 |
| 27 | 100 |
| 29.5 | 2 |
| 30 | 2 |

Flow Rate: 13 mL / min
Phenomenex Kinetex EVO C18 Prep Column
(100 Å, 5 μm, 21.5 mm X 250 mm)

Solvent A: $H_2O$ + 0.1% (v/v) TFA
Solvent B: MeCN + 0.1% (v/v) TFA

**LCMS Trace**

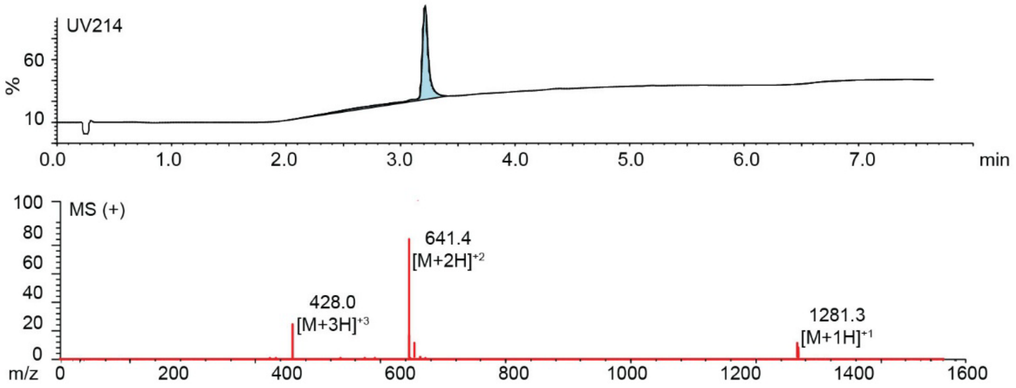

**Figure S2.31.** Synthesis summary of ACRGFLCNY-**PPA** (**18c**).
Expected: [M+1] = 1281.5219, $[M+2]^{+2}$ = 641.2610, $[M+3]^{+3}$ = 427.8406. Observed: [M+1] = 1281.3, $[M+2]^{+2}$ = 641.4, $[M+3]^{+3}$ = 428.0. Note: Two regioisomeric forms of the pyrazole are possible; the structure shown represents one isomer.

#### ARCGFLCNY-PEA

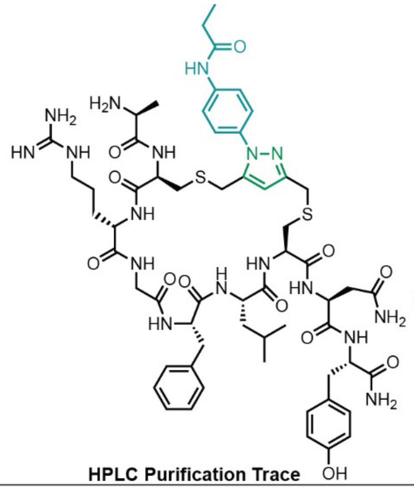

Starting material mass: 3.1 mg
Final product mass: 1.6 mg

Chemical Formula: $C_{59}H_{81}N_{17}O_{12}S_2$
Exact Mass: 1283.57
Molecular Weight: 1284.52

**HPLC Purification Trace**

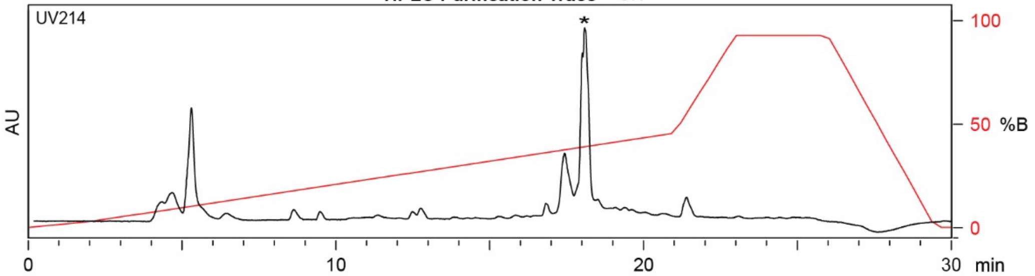

| Time (min) | Solvent B (%) |
|---|---|
| 0 | 2 |
| 2 | 2 |
| 21 | 50 |
| 24 | 100 |
| 27 | 100 |
| 29.5 | 2 |
| 30 | 2 |

Flow Rate: 13 mL / min
Phenomenex Kinetex EVO C18 Prep Column
(100 Å, 5 µm, 21.5 mm X 250 mm)

Solvent A: $H_2O$ + 0.1% (v/v) TFA
Solvent B: MeCN + 0.1% (v/v) TFA

**LCMS Trace**

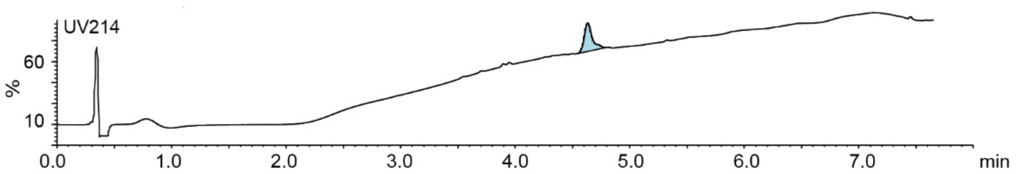

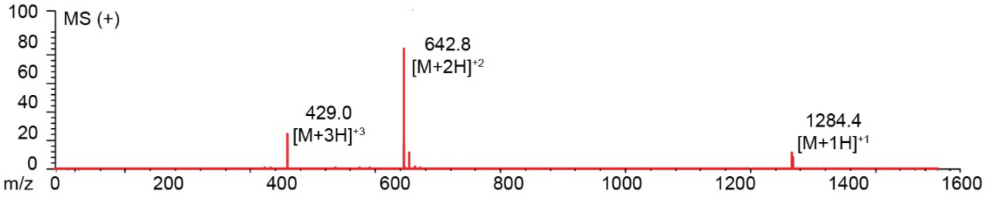

**Figure S2.32.** Synthesis summary of ACRGFLCNY-**PEA** (**18d**).
Expected: [M+1] = 1284.57, $[M+2]^{+2}$ = 642.78, $[M+3]^{+3}$ = 428.86. Observed: [M+1] = 1284.4, $[M+2]^{+2}$ = 642.8, $[M+3]^{+3}$ = 429.0. Note: Two regioisomeric forms of the pyrazole are possible; the structure shown represents one isomer.

#### ACHFDAFCT-DKL

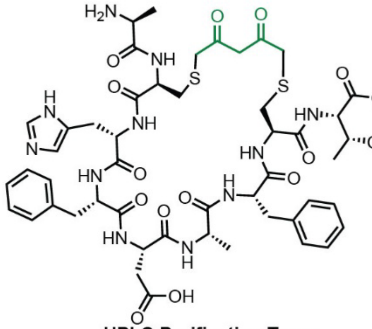

Starting material mass: mg
Final product mass: mg

Chemical Formula: $C_{49}H_{63}N_{11}O_{15}S_2$
Exact Mass: 1109.3947
Molecular Weight: 1110.2250

**HPLC Purification Trace**

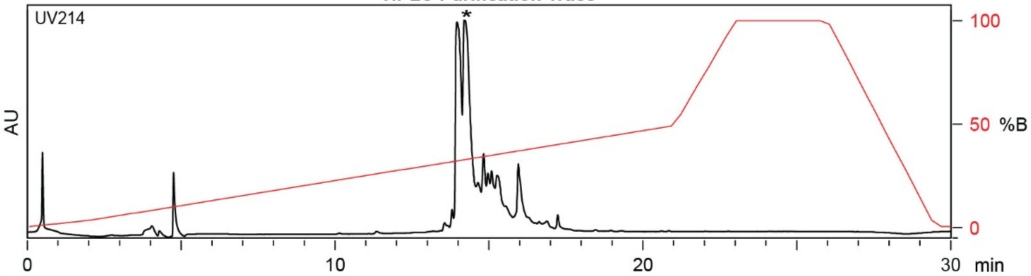

| Time (min) | Solvent B (%) |
|---|---|
| 0 | 2 |
| 2 | 2 |
| 22 | 50 |
| 24 | 100 |
| 27 | 100 |
| 28 | 2 |
| 30 | 2 |

Flow Rate: 13 mL / min
Phenomenex Kinetex EVO C18 Prep Column
(100 Å, 5 μm, 21.5 mm X 250 mm)

Solvent A: $H_2O$ + 0.1% (v/v) TFA
Solvent B: MeCN + 0.1% (v/v) TFA

**LCMS Trace**

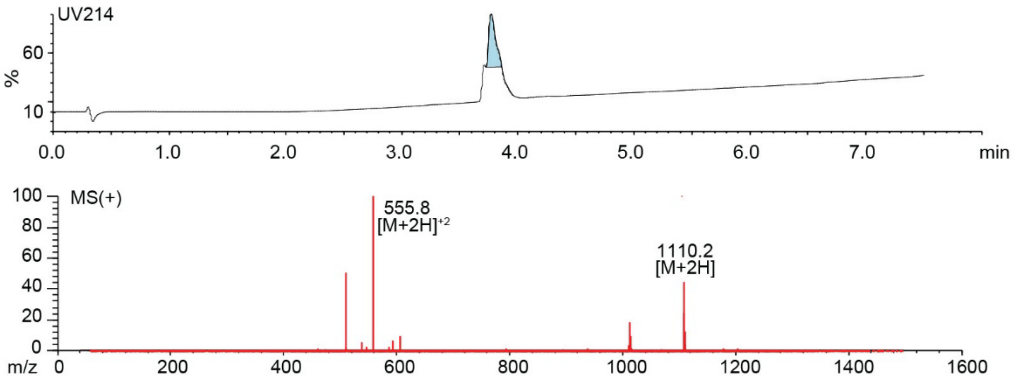

**Figure S2.33.** Synthesis summary of ACHFDAFCT-**DKL** (**19b**).
Expected: [M+1] = 1110.3947, $[M+2]^{+2}$ = 555.6974. Observed: [M+1] = 1110.2, $[M+2]^{+2}$ = 555.8.

#### ACHFDAFCT-PPA

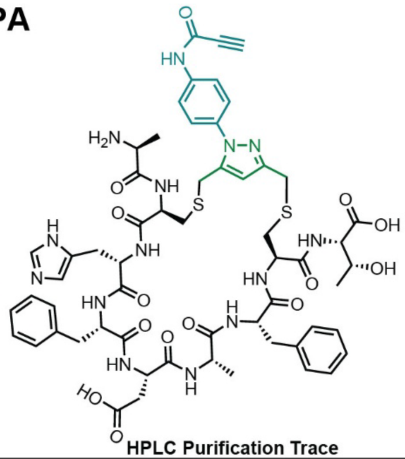

Starting material mass: mg
Final product mass: mg

Chemical Formula: $C_{58}H_{68}N_{14}O_{14}S_2$
Exact Mass: 1248.4481
Molecular Weight: 1249.3860

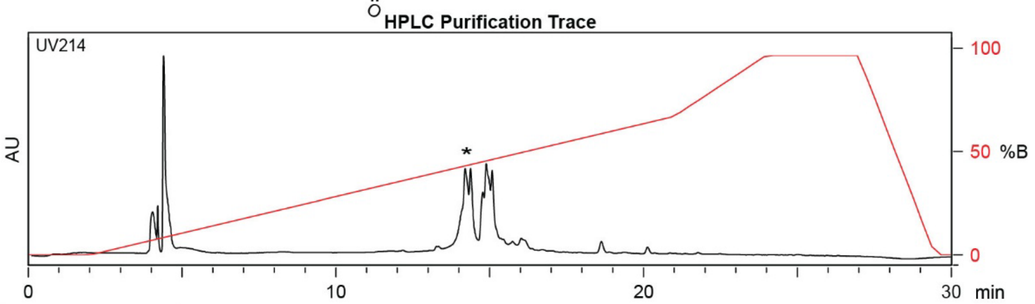

| Time (min) | Solvent B (%) |
| --- | --- |
| 0 | 2 |
| 2 | 2 |
| 21 | 70 |
| 24 | 100 |
| 27 | 100 |
| 29.5 | 2 |
| 30 | 2 |

Flow Rate: 13 mL / min
Phenomenex Kinetex EVO C18 Prep Column
(100 Å, 5 µm, 21.5 mm X 250 mm)

Solvent A: $H_2O$ + 0.1% (v/v) TFA
Solvent B: MeCN + 0.1% (v/v) TFA

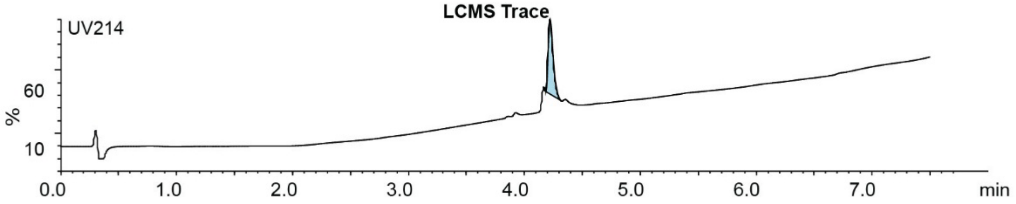

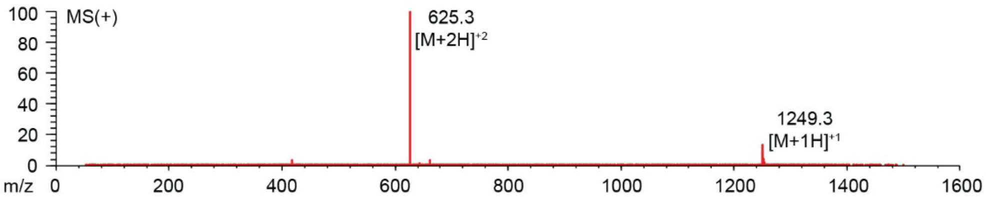

**Figure S2.34.** Synthesis summary of ACHFDAFCT-**PPA** (**19c**).
Expected: [M+1] = 1249.4481, $[M+2]^{+2}$ = 625.2240. Observed: [M+1] = 1249.3, $[M+2]^{+2}$ = 625.3.
Note: Two regioisomeric forms of the pyrazole are possible; the structure shown represents one isomer.

#### ACNFSTFCR

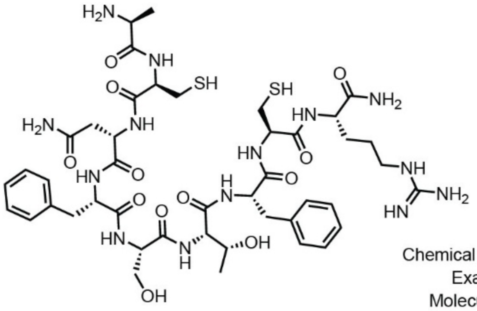

Starting material mass: N/A mg
Final product mass: 66.1 mg

Chemical Formula: $C_{44}H_{66}N_{14}O_{12}S_2$
Exact Mass: 1046.4426
Molecular Weight: 1047.2180

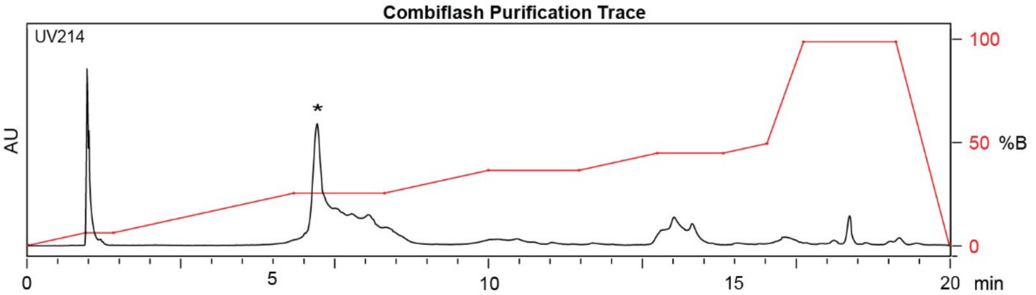

| Time (min) | Solvent B (%) |
|---|---|
| 0 | 0 |
| 1 | 5 |
| 5 | 25 |
| 10 | 40 |
| 13 | 50 |
| 16 | 100 |
| 20 | 0 |

Flow Rate: 35 mL / min
Redisep Rf Gold C18 Reverse Phase Column 30g
(100 Å, 20-40 µm, 300 ± 50 m²/g)

Solvent A: $H_2O$ + 0.1% (v/v) TFA
Solvent B: MeCN + 0.1% (v/v) TFA

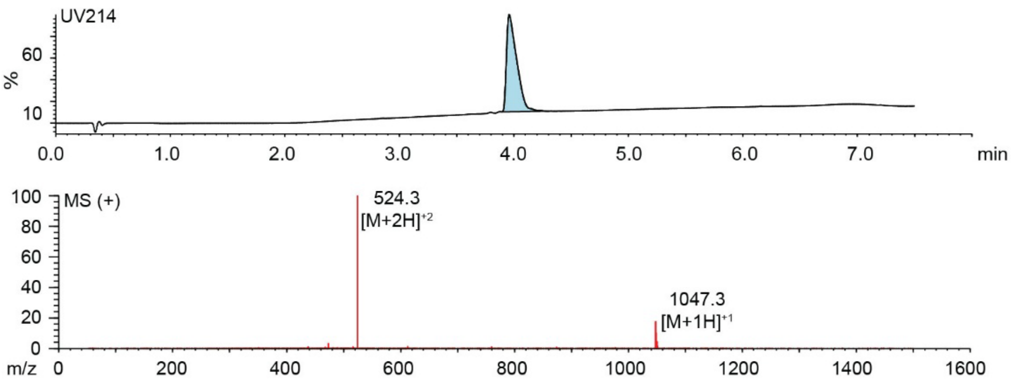

**Figure S2.35.** Synthesis summary of ACNFSTFCR (**20a**).
Expected: [M+1] = 1046.4428, $[M+2]^{+2}$ = 523.7214. Observed: [M+1] = 1047.3, $[M+2]^{+2}$ = 524.3.

#### ACNFSTFCR-DKL

Starting material mass:
Final product mass:

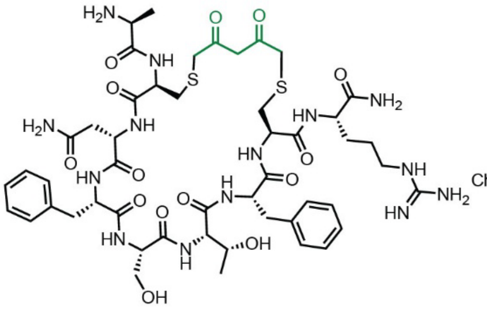

Chemical Formula: $C_{49}H_{70}N_{14}O_{14}S_2$
Exact Mass: 1142.46
Molecular Weight: 1143.30

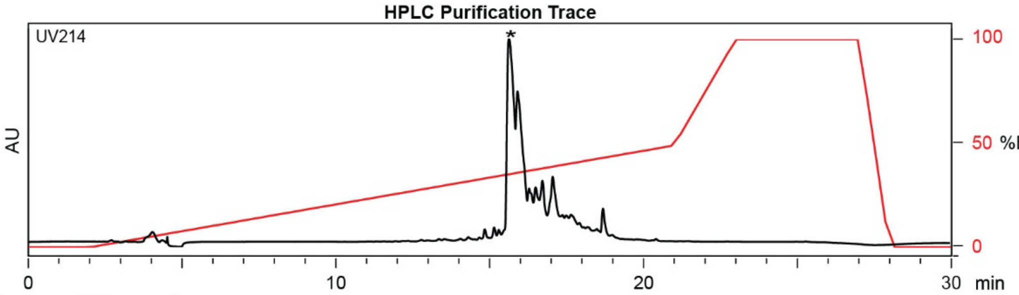

| Time (min) | Solvent B (%) |
|---|---|
| 0 | 2 |
| 2 | 2 |
| 22 | 50 |
| 24 | 100 |
| 27 | 100 |
| 28 | 2 |
| 30 | 2 |

Flow Rate: 13 mL / min
Phenomenex Kinetex EVO C18 Prep Column
(100 Å, 5 µm, 21.5 mm X 250 mm)

Solvent A: $H_2O$ + 0.1% (v/v) TFA
Solvent B: MeCN + 0.1% (v/v) TFA

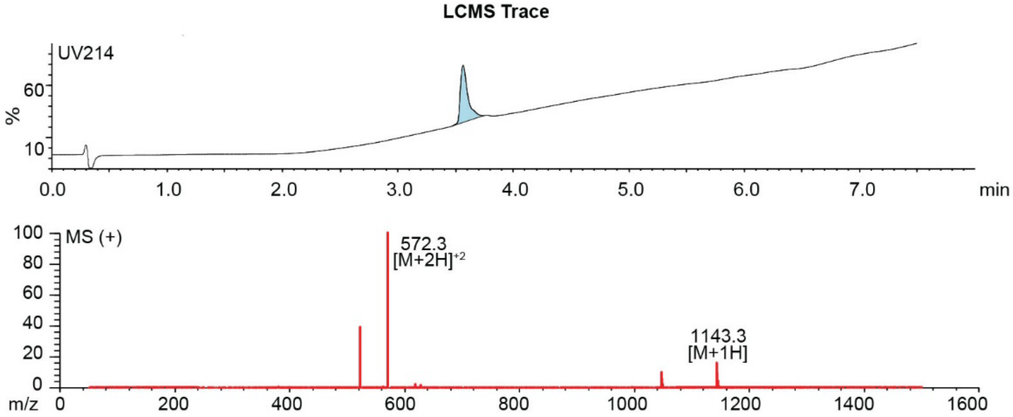

**Figure S2.36.** Synthesis summary of ACNFSTFCR-**DKL** (**20b**).
Expected: [M+1] = 1143.46, $[M+2]^{+2}$ = 572.23. Observed: [M+1] = 1143.3, $[M+2]^{+2}$ = 572.3.

#### ACNFSTFCR-PPA

Starting material mass: 3.0 mg
Final product mass: 1.6 mg

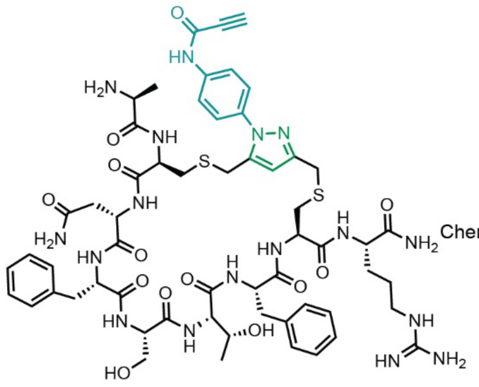

Chemical Formula: $C_{58}H_{75}N_{17}O_{13}S_2$
Exact Mass: 1281.52
Molecular Weight: 1282.46

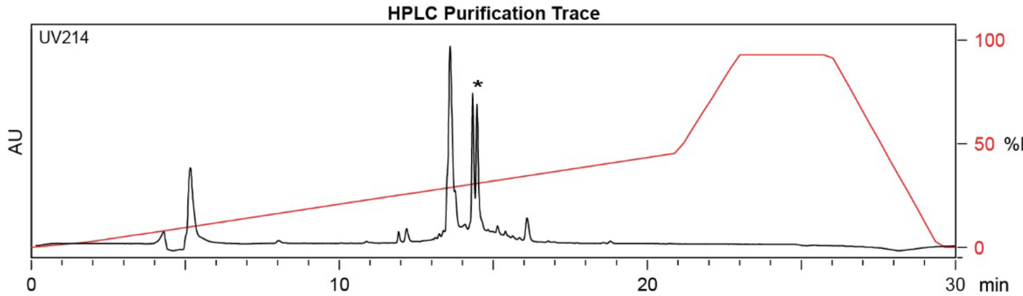

| Time (min) | Solvent B (%) |
|---|---|
| 0 | 2 |
| 2 | 2 |
| 21 | 50 |
| 24 | 100 |
| 27 | 100 |
| 29.5 | 2 |
| 30 | 2 |

Flow Rate: 13 mL / min
Phenomenex Kinetex EVO C18 Prep Column
(100 Å, 5 μm, 21.5 mm X 250 mm)

Solvent A: $H_2O$ + 0.1% (v/v) TFA
Solvent B: MeCN + 0.1% (v/v) TFA

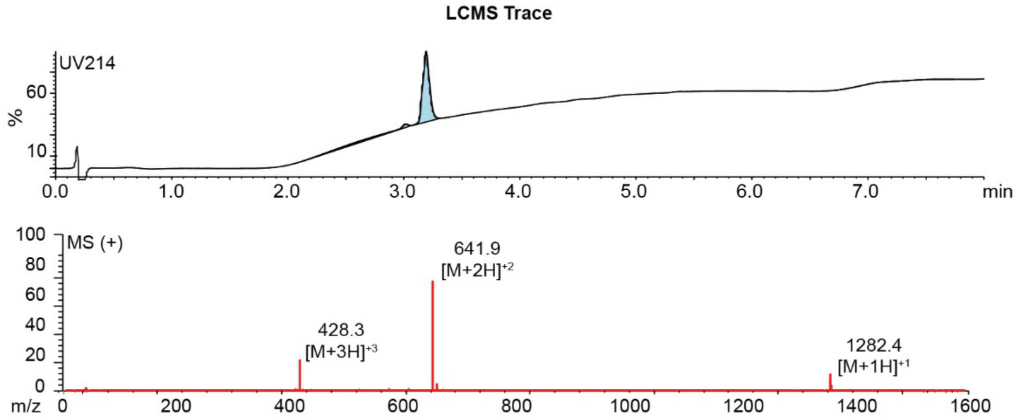

**Figure S2.37.** Synthesis summary of ACNFSTFCR-**PPA** (**20c**).
Expected: [M+1] = 1282.52, $[M+2]^{+2}$ = 641.76, $[M+3]^{+3}$ = 428.17. Observed: [M+1] = 1282.4, $[M+2]^{+2}$ = 641.9, $[M+3]^{+3}$ = 428.3. Note: Two regioisomeric forms of the pyrazole are possible; the structure shown represents one isomer.

#### ACNFSTFCR-PEA

Starting material mass: 3.0 mg
Final product mass: 1.6 mg

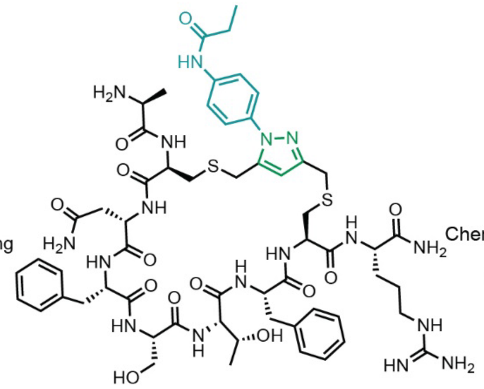

Chemical Formula: $C_{58}H_{79}N_{17}O_{13}S_2$
Exact Mass: 1285.55
Molecular Weight: 1286.50

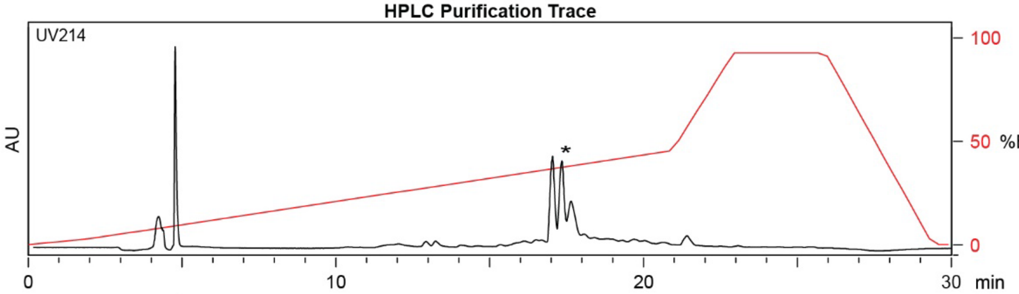

| Time (min) | Solvent B (%) |
|---|---|
| 0 | 2 |
| 2 | 2 |
| 21 | 50 |
| 24 | 100 |
| 27 | 100 |
| 29.5 | 2 |
| 30 | 2 |

Flow Rate: 13 mL / min
Phenomenex Kinetex EVO C18 Prep Column
(100 Å, 5 µm, 21.5 mm X 250 mm)

Solvent A: $H_2O$ + 0.1% (v/v) TFA
Solvent B: MeCN + 0.1% (v/v) TFA

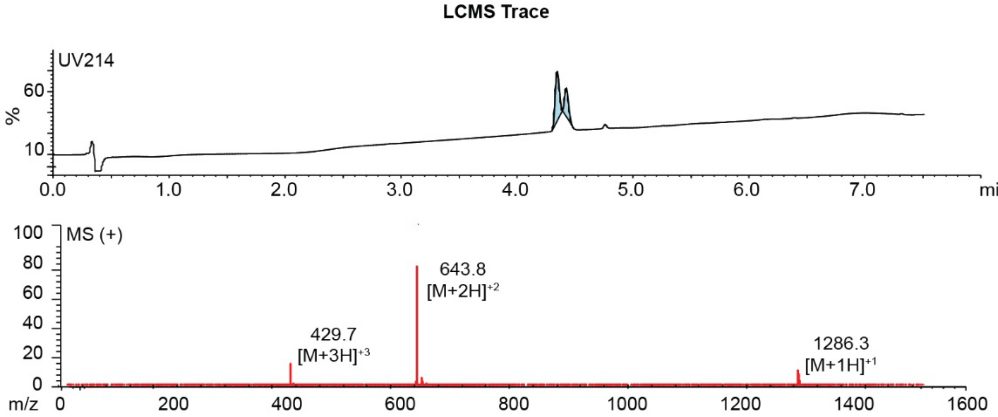

**Figure S2.38.** Synthesis summary of ACNFSTFCR-**PEA** (**20d**).
Expected: [M+1] = 1286.55, $[M+2]^{+2}$ = 643.78, $[M+3]^{+3}$ = 429.52. Observed: [M+1] = 1286.3, $[M+2]^{+2}$ = 643.8, $[M+3]^{+3}$ = 429.7. Note: Two regioisomeric forms of the pyrazole are possible; the structure shown represents one isomer.

### ACVPYFRCI

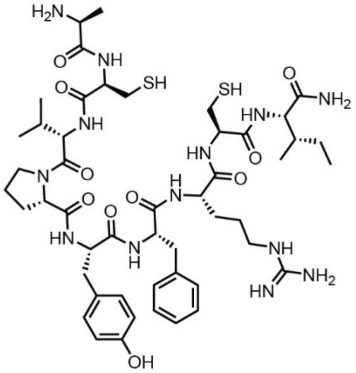

Starting material mass: N/A mg
Final product mass: 70.7 mg

Chemical Formula: $C_{49}H_{75}N_{13}O_{10}S_2$
Exact Mass: 1069.52
Molecular Weight: 1070.34

Combiflash Purification Trace

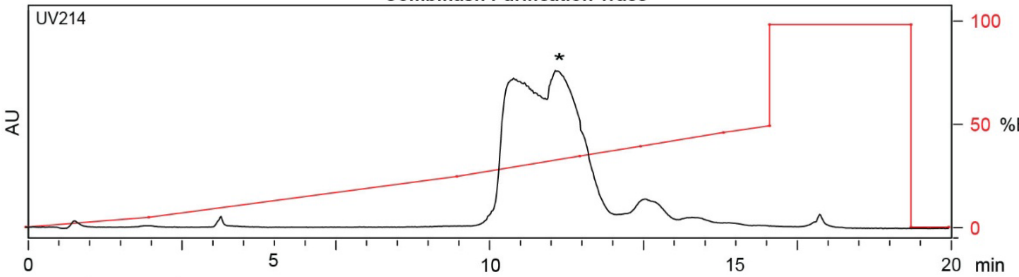

| Time (min) | Solvent B (%) |
|---|---|
| 0 | 0 |
| 1 | 2 |
| 5 | 20 |
| 10 | 40 |
| 15.5 | 100 |
| 17.8 | 0 |
| 20 | 0 |

Flow Rate: 35 mL / min
Redisep Rf Gold C18 Reverse Phase Column 30g
(100 Å, 20-40 μm, 300 ± 50 $m^2/g$)

Solvent A: $H_2O$ + 0.1% (v/v) TFA
Solvent B: MeCN + 0.1% (v/v) TFA

LCMS Trace

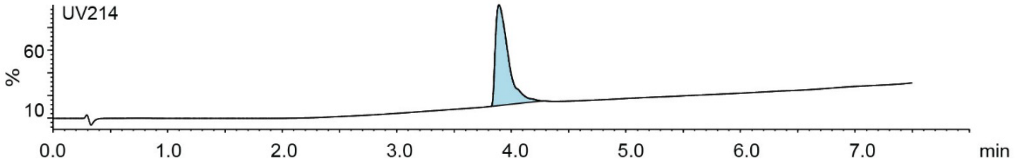

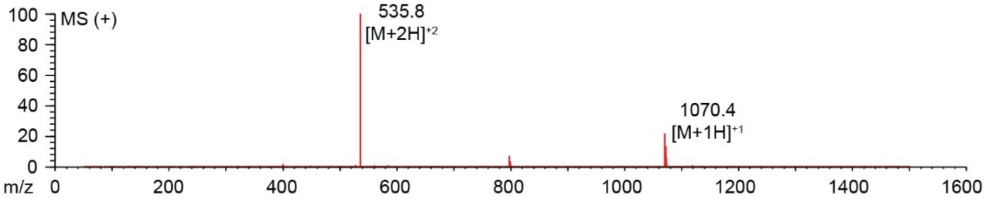

**Figure S2.39.** Synthesis summary of ACVPYFRCI (**21a**).
Expected: [M+1] = 1070.52, $[M+2]^{\div 2}$ = 535.76. Observed: [M+1] = 1070.4, $[M+2]^{\div 2}$ = 535.8.

#### ACVPYFRCI-DKL

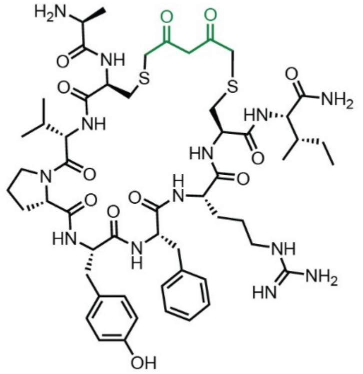

Starting material mass: 5.4 mg
Final product mass: 2.3 mg

Chemical Formula: $C_{54}H_{79}N_{13}O_{12}S_2$
Exact Mass: 1165.5413
Molecular Weight: 1166.4250

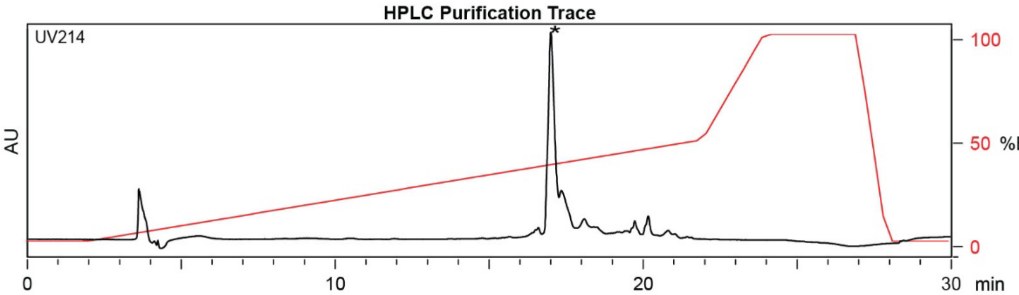

| Time (min) | Solvent B (%) |
|---|---|
| 0 | 2 |
| 2 | 2 |
| 22 | 50 |
| 24 | 100 |
| 27 | 100 |
| 28 | 2 |
| 30 | 2 |

Flow Rate: 13 mL / min
Phenomenex Kinetex EVO C18 Prep Column
(100 Å, 5 μm, 21.5 mm X 250 mm)

Solvent A: $H_2O$ + 0.1% (v/v) TFA
Solvent B: MeCN + 0.1% (v/v) TFA

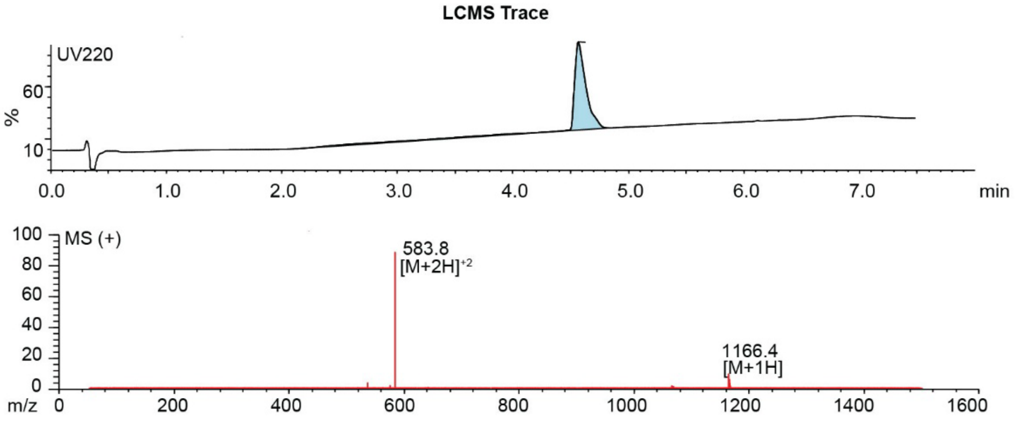

**Figure S2.40.** Synthesis summary of ACVPYFRCI-**DKL** (**21b**).
Expected: [M+1] = 1166.5413, $[M+2]^{+2}$ = 583.7706. Observed: [M+1] = 1166.4, $[M+2]^{+2}$ = 583.8.

#### ACVPYFRCI-PPA

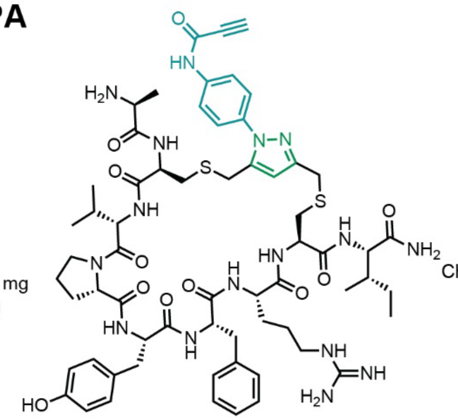

Starting material mass: 2.7 mg
Final product mass: 0.8 mg

Chemical Formula: $C_{63}H_{84}N_{16}O_{11}S_2$
Exact Mass: 1304.59
Molecular Weight: 1305.59

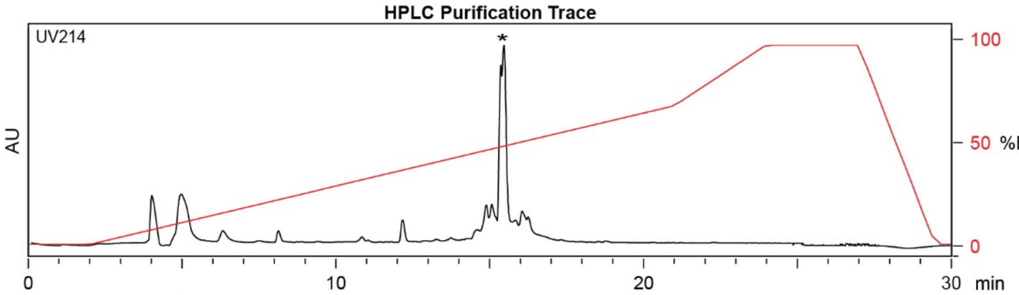

| Time (min) | Solvent B (%) |
|---|---|
| 0 | 2 |
| 2 | 2 |
| 21 | 70 |
| 24 | 100 |
| 27 | 100 |
| 29.5 | 2 |
| 30 | 2 |

Flow Rate: 13 mL / min
Phenomenex Kinetex EVO C18 Prep Column
(100 Å, 5 µm, 21.5 mm X 250 mm)

Solvent A: $H_2O$ + 0.1% (v/v) TFA
Solvent B: MeCN + 0.1% (v/v) TFA

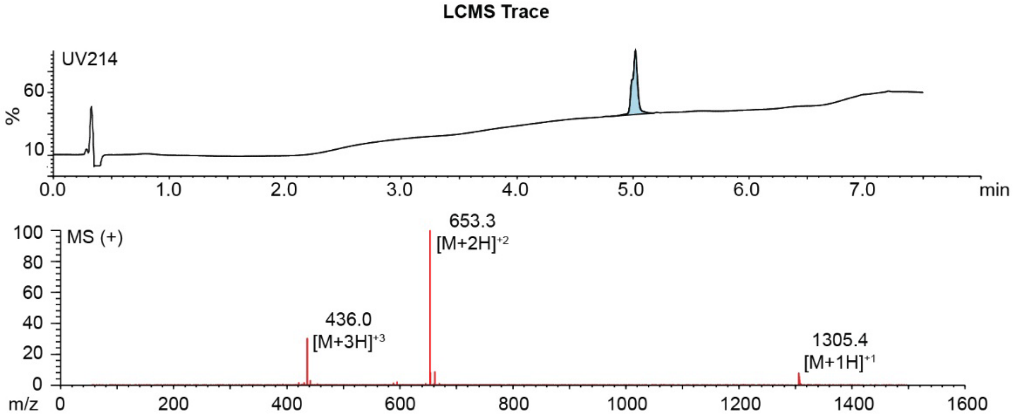

**Figure S2.41.** Synthesis summary of ACVPYFRCI-**PPA** (**21c**).
Expected: [M+1] = 1305.59, $[M+2]^{+2}$ = 653.30, $[M+3]^{+3}$ = 435.86. Observed: [M+1] = 1305.4, $[M+2]^{+2}$ = 653.3, $[M+3]^{+3}$ = 436.0. Note: Two regioisomeric forms of the pyrazole are possible; the structure shown represents one isomer.

#### ACVPYFRCI-PEA

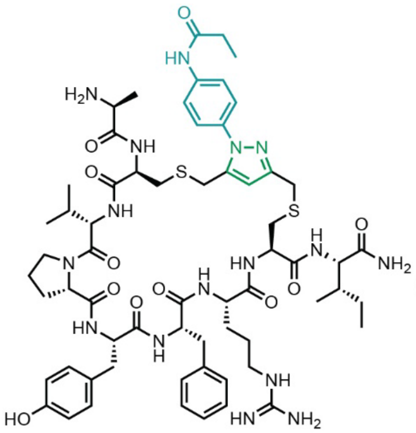

Starting material mass: 3.0 mg
Final product mass: 1.3 mg

Chemical Formula: $C_{63}H_{88}N_{16}O_{11}S_2$
Exact Mass: 1308.63
Molecular Weight: 1309.62

HPLC Purification Trace

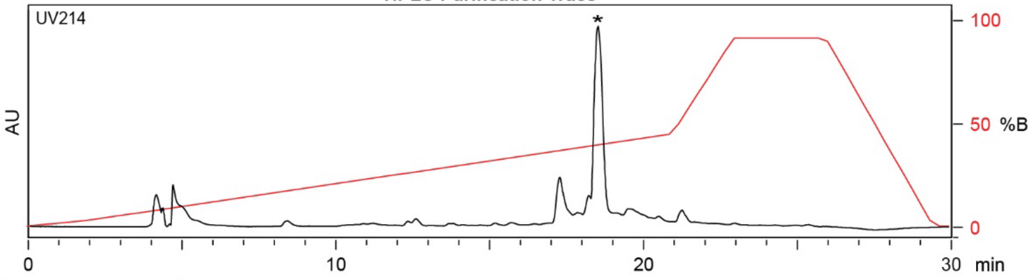

| Time (min) | Solvent B (%) |
|---|---|
| 0 | 2 |
| 2 | 2 |
| 21 | 50 |
| 24 | 100 |
| 27 | 100 |
| 29.5 | 2 |
| 30 | 2 |

Flow Rate: 13 mL / min
Phenomenex Kinetex EVO C18 Prep Column
(100 Å, 5 µm, 21.5 mm X 250 mm)

Solvent A: $H_2O$ + 0.1% (v/v) TFA
Solvent B: MeCN + 0.1% (v/v) TFA

LCMS Trace

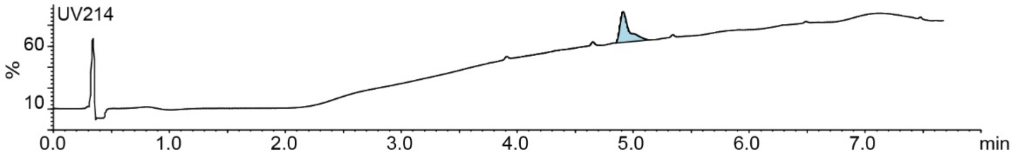

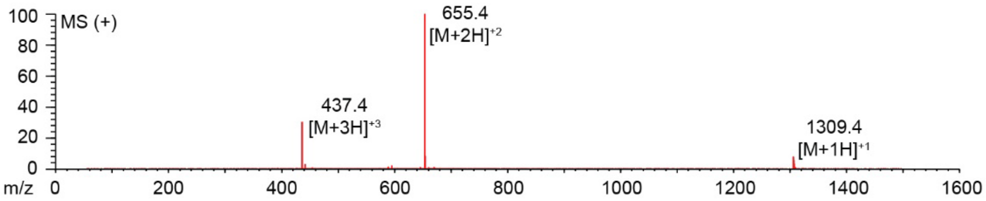

**Figure S2.42.** Synthesis summary of ACVPYFRCI-**PEA** (**21d**).
Expected: [M+1] = 1309.63, $[M+2]^{+2}$ = 655.32, $[M+3]^{+3}$ = 437.21. Observed: [M+1] = 1309.4, $[M+2]^{+2}$ = 655.4, $[M+3]^{+3}$ = 437.4. Note: Two regioisomeric forms of the pyrazole are possible; the structure shown represents one isomer.

### ACWSWTCY-DKL

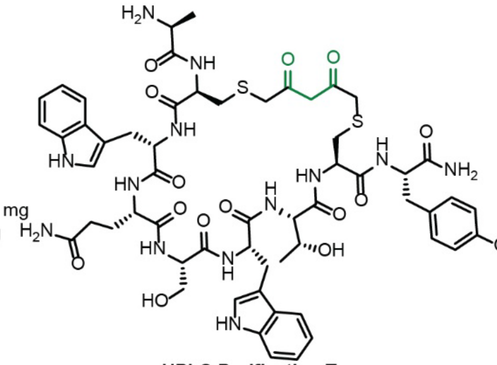

Starting material mass: mg
Final product mass: mg

Chemical Formula: $C_{57}H_{71}N_{13}O_{15}S_2$
Exact Mass: 1241.4634
Molecular Weight: 1242.3910

**HPLC Purification Trace**

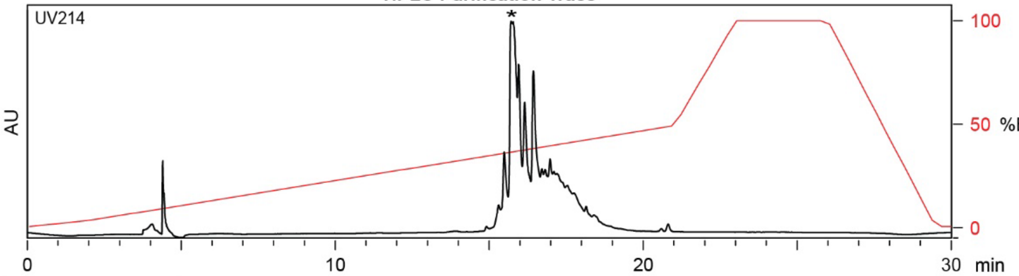

| Time (min) | Solvent B (%) |
|---|---|
| 0 | 2 |
| 2 | 2 |
| 22 | 50 |
| 24 | 100 |
| 27 | 100 |
| 28 | 2 |
| 30 | 2 |

Flow Rate: 13 mL / min
Phenomenex Kinetex EVO C18 Prep Column
(100 Å, 5 µm, 21.5 mm X 250 mm)

Solvent A: $H_2O$ + 0.1% (v/v) TFA
Solvent B: MeCN + 0.1% (v/v) TFA

**LCMS Trace**

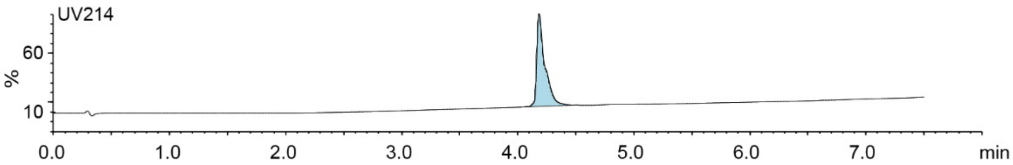

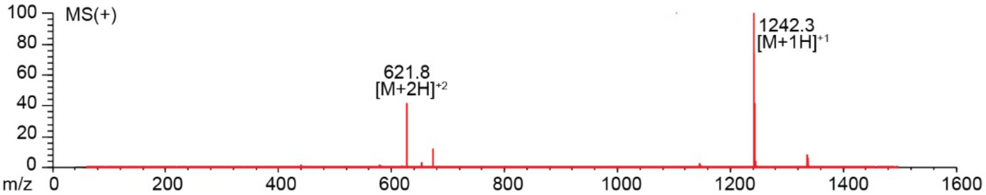

**Figure S2.43.** Synthesis summary of ACWSWTCY-**DKL** (**22b**).
Expected: [M+1] = 1242.4634, $[M+2]^{+2}$ = 621.7317. Observed: [M+1] = 1242.3, $[M+2]^{+2}$ = 621.8.

#### ACWQSWTCY-PPA

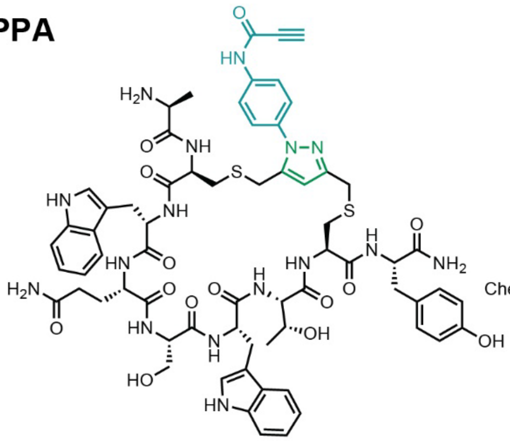

Starting material mass: mg
Final product mass: mg

Chemical Formula: $C_{66}H_{76}N_{16}O_{14}S_2$
Exact Mass: 1380.52
Molecular Weight: 1381.55

**HPLC Purification Trace**

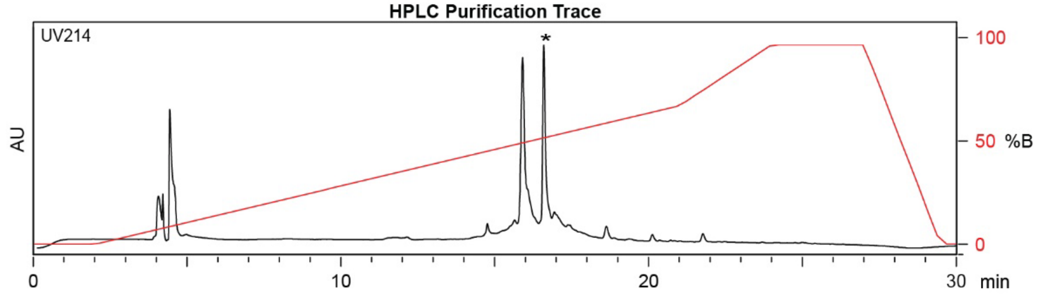

| Time (min) | Solvent B (%) |
|---|---|
| 0 | 2 |
| 2 | 2 |
| 21 | 70 |
| 24 | 100 |
| 27 | 100 |
| 29.5 | 2 |
| 30 | 2 |

Flow Rate: 13 mL / min
Phenomenex Kinetex EVO C18 Prep Column
(100 Å, 5 µm, 21.5 mm X 250 mm)

Solvent A: $H_2O$ + 0.1% (v/v) TFA
Solvent B: MeCN + 0.1% (v/v) TFA

**LCMS Trace**

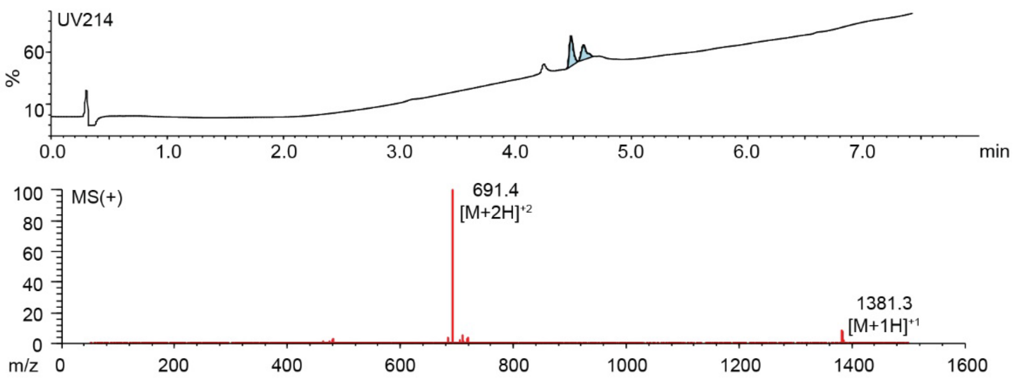

**Figure S2.44.** Synthesis summary of ACWSWTCY-**PPA** (**22c**).
Expected: [M+1] = 1381.52, $[M+2]^{\div 2}$ = 691.26. Observed: [M+1] =1381.3 , $[M+2]^{\div 2}$ = 691.4. Note: Two regioisomeric forms of the pyrazole are possible; the structure shown represents one isomer.

#### ACASRFHEC-DKL

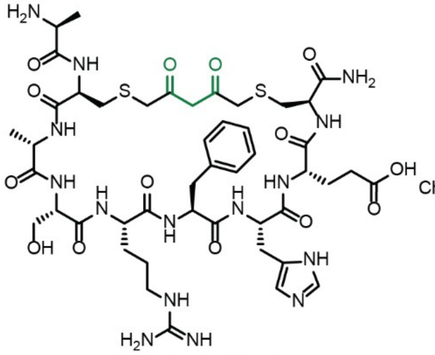

Starting material mass: mg
Final product mass: mg

Chemical Formula: $C_{46}H_{67}N_{15}O_{14}S_2$
Exact Mass: 1117.4433
Molecular Weight: 1118.2530

**HPLC Purification Trace**

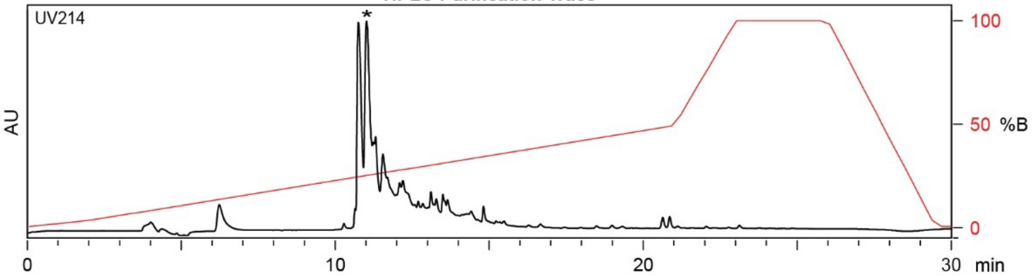

| Time (min) | Solvent B (%) |
|---|---|
| 0 | 2 |
| 2 | 2 |
| 22 | 50 |
| 24 | 100 |
| 27 | 100 |
| 28 | 2 |
| 30 | 2 |

Flow Rate: 13 mL / min
Phenomenex Kinetex EVO C18 Prep Column
(100 Å, 5 µm, 21.5 mm X 250 mm)

Solvent A: $H_2O$ + 0.1% (v/v) TFA
Solvent B: MeCN + 0.1% (v/v) TFA

**LCMS Trace**

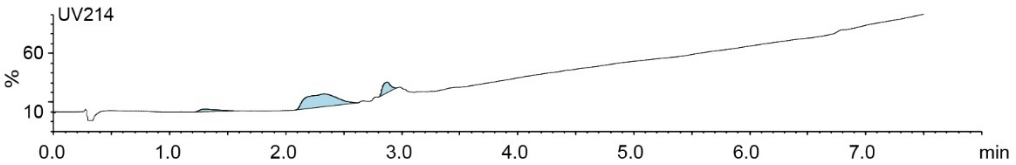

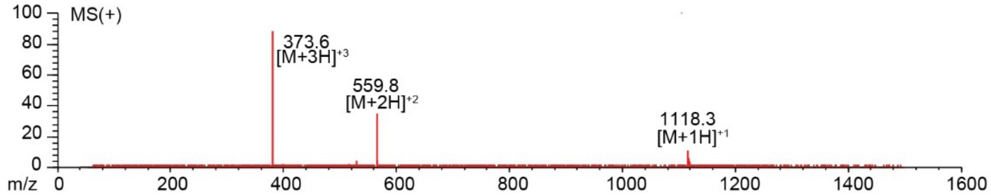

**Figure S2.45.** Synthesis summary of ACASRFHEC-**DKL** (**23b**).
Expected: [M+1] = 1118.4433, $[M+2]^{\div 2}$ = 559.7216, $[M+3]^{\div 3}$ = 373.4811. Observed: [M+1] = 1118.3, $[M+2]^{\div 2}$ = 559.8, $[M+3]^{\div 3}$ = 373.6.

#### ACASRFHEC-PPA

Starting material mass: mg
Final product mass: mg

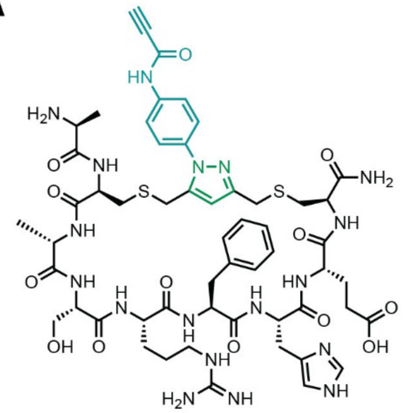

Chemical Formula: $C_{55}H_{72}N_{18}O_{13}S_2$
Exact Mass: 1256.4968
Molecular Weight: 1257.4140

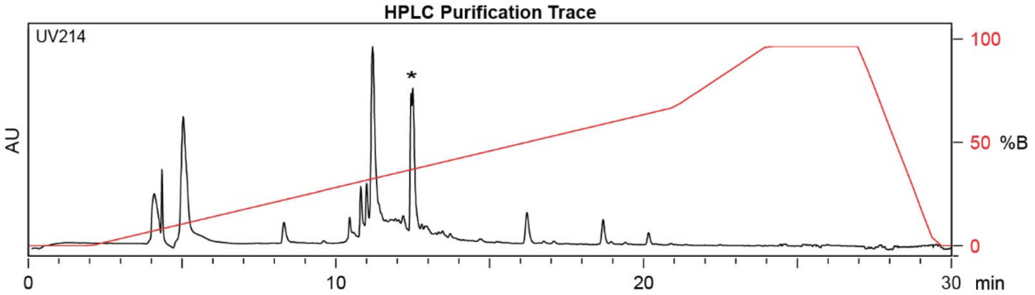

| Time (min) | Solvent B (%) |
|---|---|
| 0 | 2 |
| 2 | 2 |
| 21 | 70 |
| 24 | 100 |
| 27 | 100 |
| 29.5 | 2 |
| 30 | 2 |

Flow Rate: 13 mL / min
Phenomenex Kinetex EVO C18 Prep Column
(100 Å, 5 μm, 21.5 mm X 250 mm)

Solvent A: $H_2O$ + 0.1% (v/v) TFA
Solvent B: MeCN + 0.1% (v/v) TFA

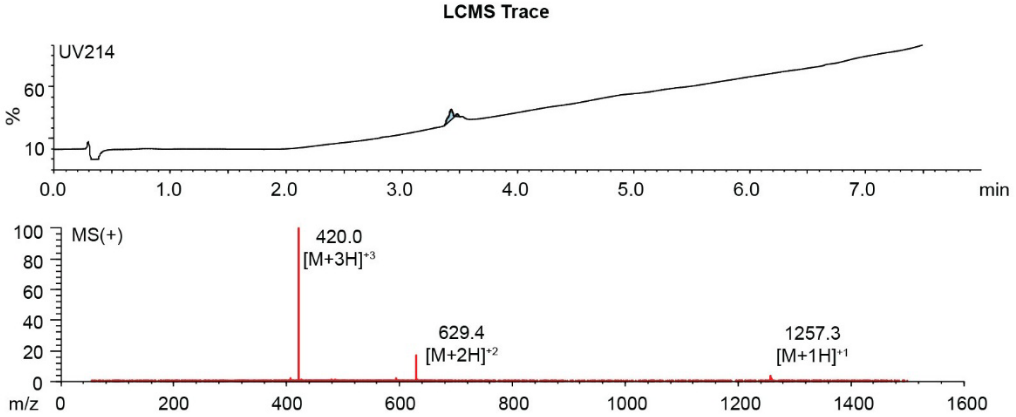

**Figure S2.46.** Synthesis summary of ACASRFHEC-**PPA** (**23c**).
Expected: [M+1] = 1257.4968, $[M+2]^{+2}$ = 629.2484, $[M+3]^{+3}$ = 419.8323. Observed: [M+1] = 1257.3, $[M+2]^{+2}$ = 629.4, $[M+3]^{+3}$ = 420.0. Note: Two regioisomeric forms of the pyrazole are possible; the structure shown represents one isomer.

#### ACVGSFFSC-DKL

Starting material mass: mg
Final product mass: mg

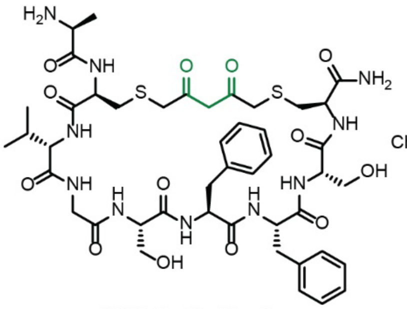

Chemical Formula: $C_{45}H_{62}N_{10}O_{13}S_2$
Exact Mass: 1014.3939
Molecular Weight: 1015.1680

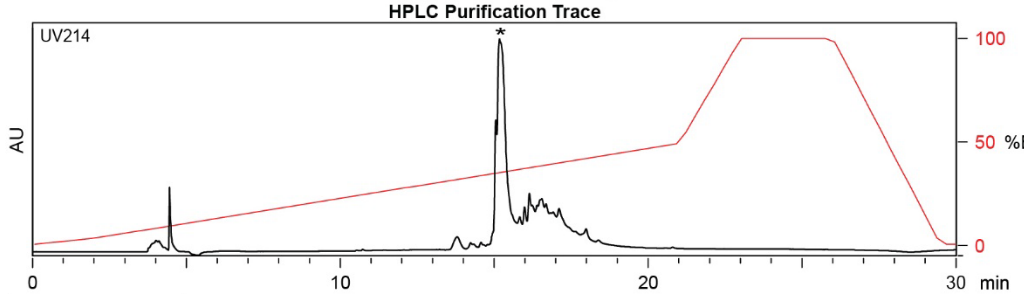

| Time (min) | Solvent B (%) |
|---|---|
| 0 | 2 |
| 2 | 2 |
| 22 | 50 |
| 24 | 100 |
| 27 | 100 |
| 28 | 2 |
| 30 | 2 |

Flow Rate: 13 mL / min
Phenomenex Kinetex EVO C18 Prep Column
(100 Å, 5 μm, 21.5 mm X 250 mm)

Solvent A: $H_2O$ + 0.1% (v/v) TFA
Solvent B: MeCN + 0.1% (v/v) TFA

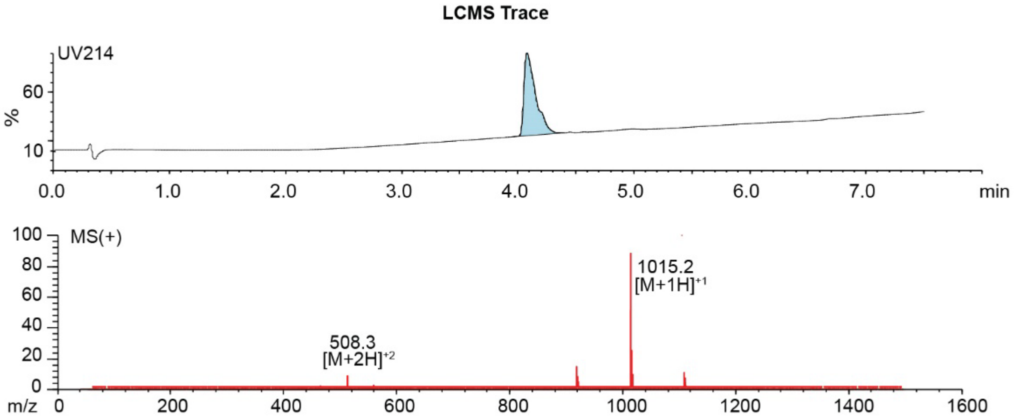

**Figure S2.47.** Synthesis summary of ACVGSFFSC-**DKL** (**24b**).
Expected: [M+1] = 1015.3939, $[M+2]^{+2}$ = 508.1970. Observed: [M+1] =1015.2 , $[M+2]^{+2}$ = 508.3.

#### ACVGSFFSC-PPA

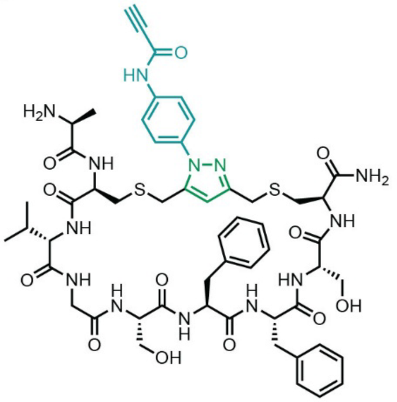

Starting material mass: mg
Final product mass: mg

Chemical Formula: $C_{54}H_{67}N_{13}O_{12}S_2$
Exact Mass: 1153.45
Molecular Weight: 1154.33

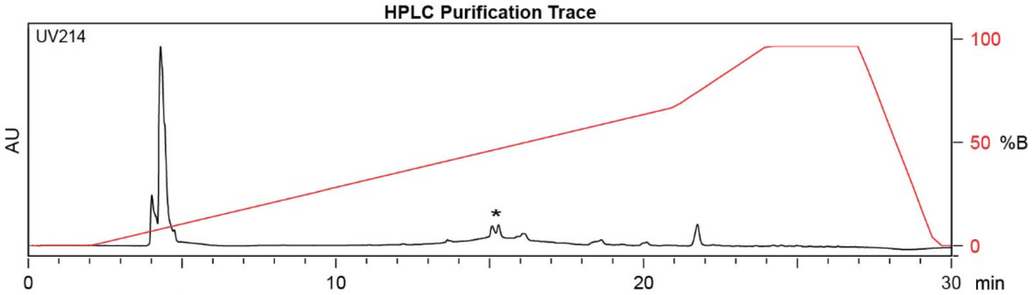

| Time (min) | Solvent B (%) |
|---|---|
| 0 | 2 |
| 2 | 2 |
| 21 | 70 |
| 24 | 100 |
| 27 | 100 |
| 29.5 | 2 |
| 30 | 2 |

Flow Rate: 13 mL / min
Phenomenex Kinetex EVO C18 Prep Column
(100 Å, 5 µm, 21.5 mm X 250 mm)

Solvent A: $H_2O$ + 0.1% (v/v) TFA
Solvent B: MeCN + 0.1% (v/v) TFA

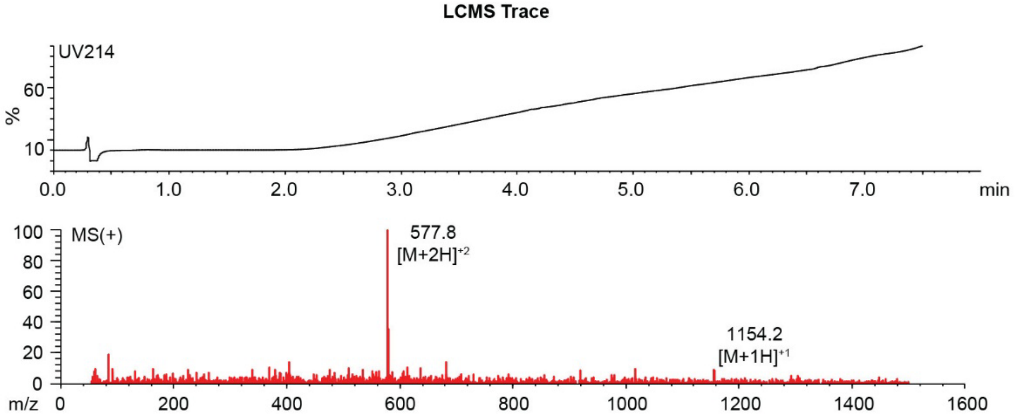

**Figure S2.48.** Synthesis summary of ACVGSFFSC-**PPA** (**24c**).
Expected: [M+1] = 1154.45, $[M+2]^{+2}$ = 577.72. Observed: [M+1] = 1154.2, $[M+2]^{+2}$ = 577.8. Note: Two regioisomeric forms of the pyrazole are possible; the structure shown represents one isomer.

#### ACNFNKSSC-DKL

Starting material mass: mg
Final product mass: mg

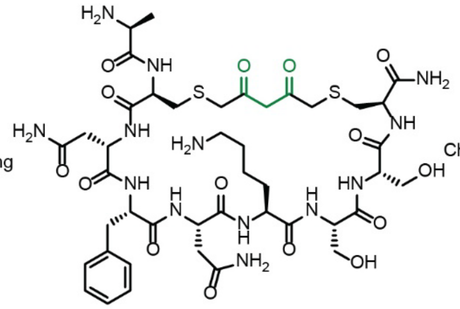

Chemical Formula: $C_{43}H_{65}N_{13}O_{15}S_2$
Exact Mass: 1067.4164
Molecular Weight: 1068.1890

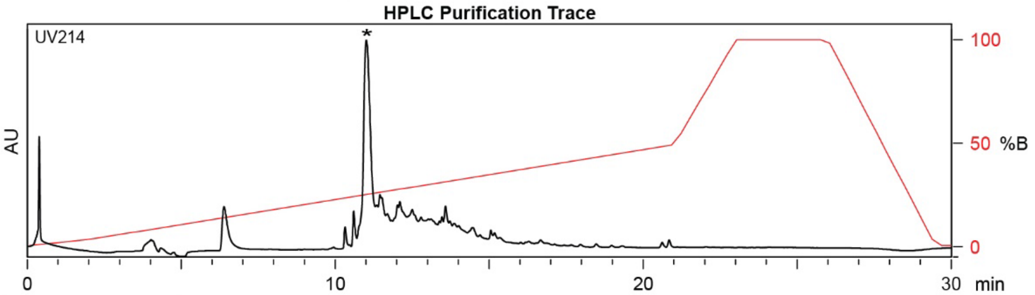

| Time (min) | Solvent B (%) |
|---|---|
| 0 | 2 |
| 2 | 2 |
| 22 | 50 |
| 24 | 100 |
| 27 | 100 |
| 28 | 2 |
| 30 | 2 |

Flow Rate: 13 mL / min
Phenomenex Kinetex EVO C18 Prep Column
(100 Å, 5 µm, 21.5 mm X 250 mm)

Solvent A: $H_2O$ + 0.1% (v/v) TFA
Solvent B: MeCN + 0.1% (v/v) TFA

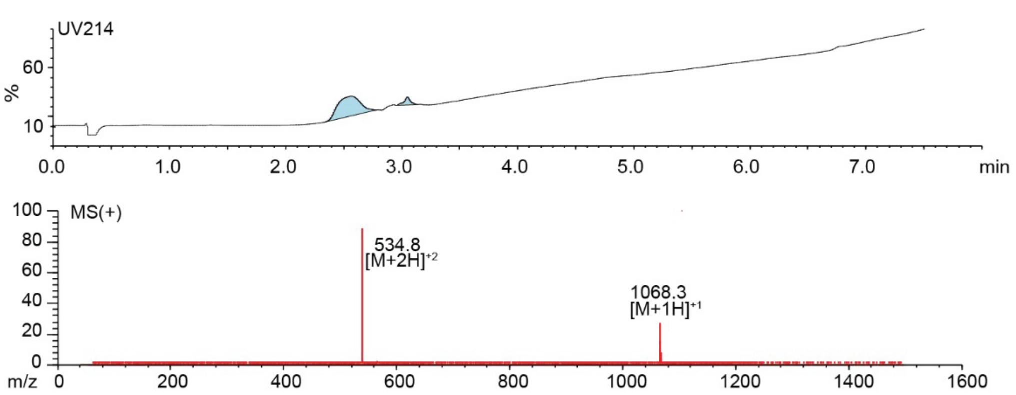

**Figure S2.49.** Synthesis summary of ACNFNKSSC-**DKL** (**25b**).
Expected: [M+1] = 1068.4164, $[M+2]^{+2}$ = 534.2082. Observed: [M+1] = 1068.3, $[M+2]^{+2}$ = 534.8.

#### ACNFNKSSC-PPA

Starting material mass: mg
Final product mass: mg

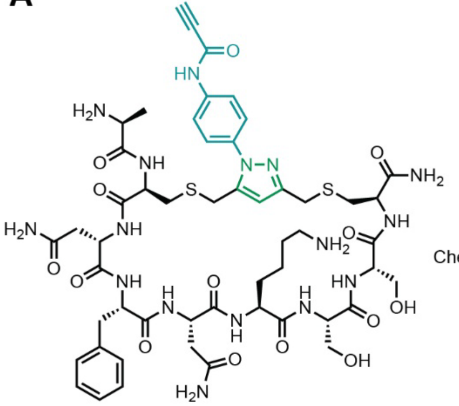

Chemical Formula: $C_{52}H_{70}N_{16}O_{14}S_2$
Exact Mass: 1206.47
Molecular Weight: 1207.35

**HPLC Purification Trace**

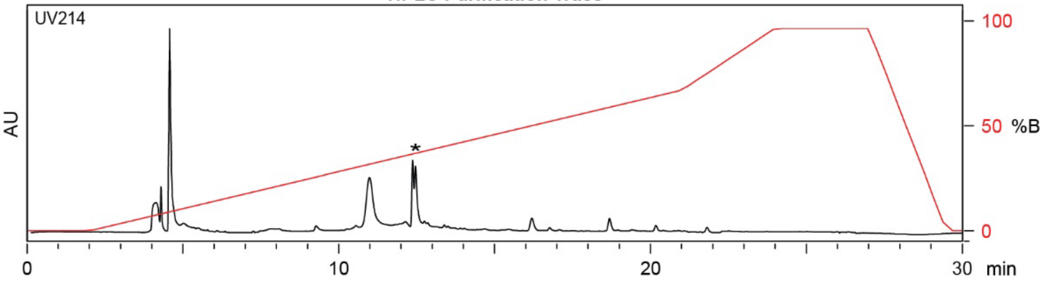

| Time (min) | Solvent B (%) |
|---|---|
| 0 | 2 |
| 2 | 2 |
| 21 | 70 |
| 24 | 100 |
| 27 | 100 |
| 29.5 | 2 |
| 30 | 2 |

Flow Rate: 13 mL / min
Phenomenex Kinetex EVO C18 Prep Column
(100 Å, 5 μm, 21.5 mm X 250 mm)

Solvent A: $H_2O$ + 0.1% (v/v) TFA
Solvent B: MeCN + 0.1% (v/v) TFA

**LCMS Trace**

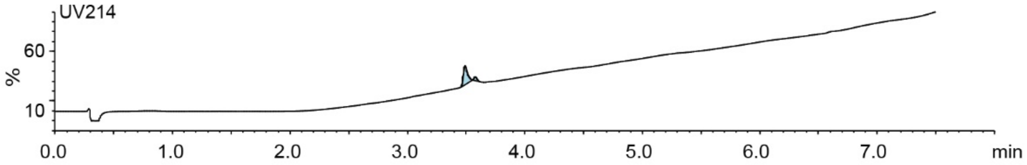

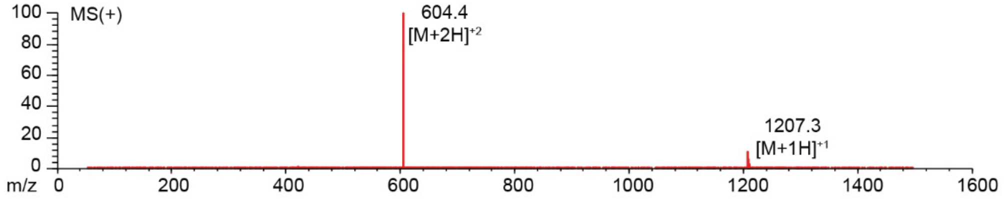

**Figure S2.50.** Synthesis summary of ACNFNKSSC-**PPA** (**25c**).
Expected: [M+1] = 1207.47, $[M+2]^{+2}$ = 603.74. Observed: [M+1] =1207.3 , $[M+2]^{+2}$ = 604.4. Note: Two regioisomeric forms of the pyrazole are possible; the structure shown represents one isomer.

#### ACLPFYSRC

Starting material mass: N/A mg
Final product mass: 94.4 mg

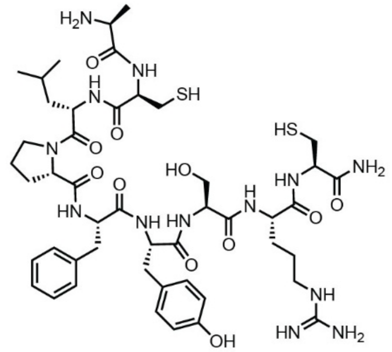

Chemical Formula: $C_{47}H_{71}N_{13}O_{11}S_2$
Exact Mass: 1057.48
Molecular Weight: 1058.29

**Combiflash Purification Trace**

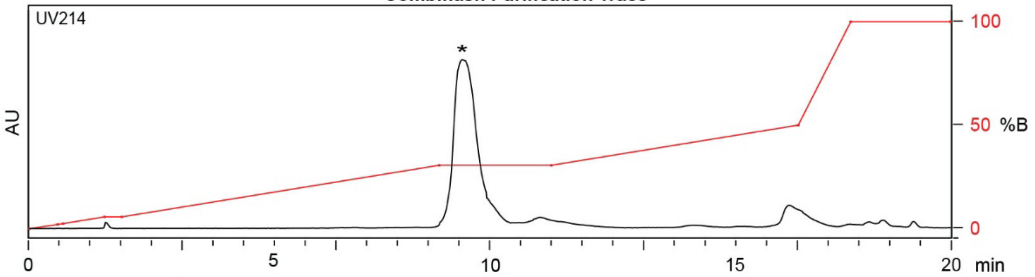

| Time (min) | Solvent B (%) |
|---|---|
| 0 | 0 |
| 1 | 2 |
| 5 | 25 |
| 8 | 35 |
| 11 | 35 |
| 16 | 100 |
| 20 | 100 |

Flow Rate: 35 mL / min
Redisep Rf Gold C18 Reverse Phase Column 30g
(100 Å, 20-40 µm, 300 ± 50 m²/g)

Solvent A: $H_2O$ + 0.1% (v/v) TFA
Solvent B: MeCN + 0.1% (v/v) TFA

**LCMS Trace**

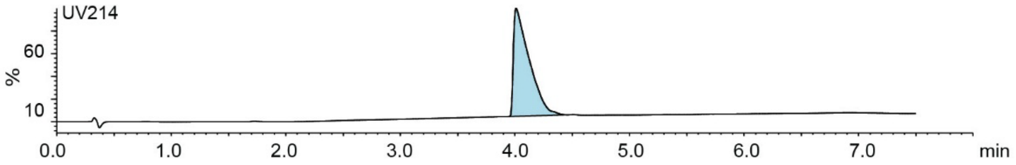

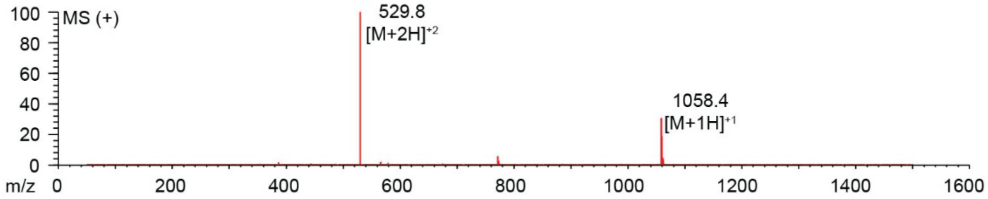

**Figure S2.51.** Synthesis summary of ACLPFYSRC (**26a**).
Expected: [M+1] = 1058.48, $[M+2]^{\div 2}$ = 529.74. Observed: [M+1] = 1058.4, $[M+2]^{\div 2}$ = 529.8.

#### ACLPFYSRC-DKL

Starting material mass:
Final product mass:

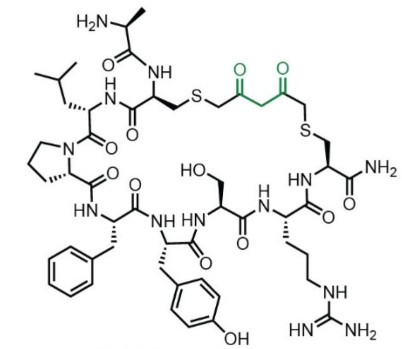

Chemical Formula: $C_{52}H_{75}N_{13}O_{13}S_2$
Exact Mass: 1153.50
Molecular Weight: 1154.37

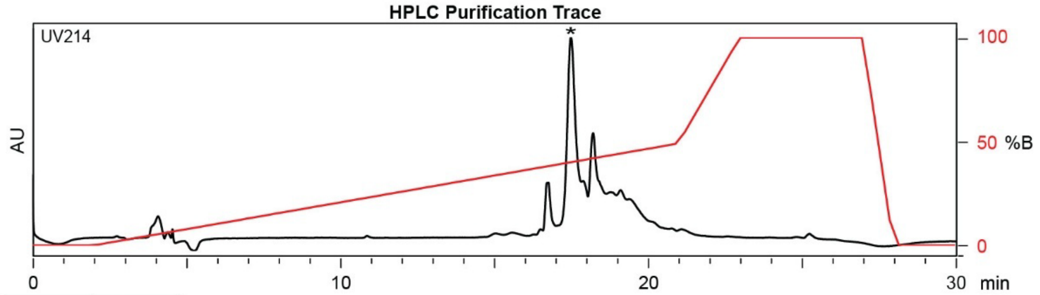

| Time (min) | Solvent B (%) |
|---|---|
| 0 | 2 |
| 2 | 2 |
| 22 | 50 |
| 24 | 100 |
| 27 | 100 |
| 28 | 2 |
| 30 | 2 |

Flow Rate: 13 mL / min
Phenomenex Kinetex EVO C18 Prep Column
(100 Å, 5 μm, 21.5 mm X 250 mm)

Solvent A: $H_2O$ + 0.1% (v/v) TFA
Solvent B: MeCN + 0.1% (v/v) TFA

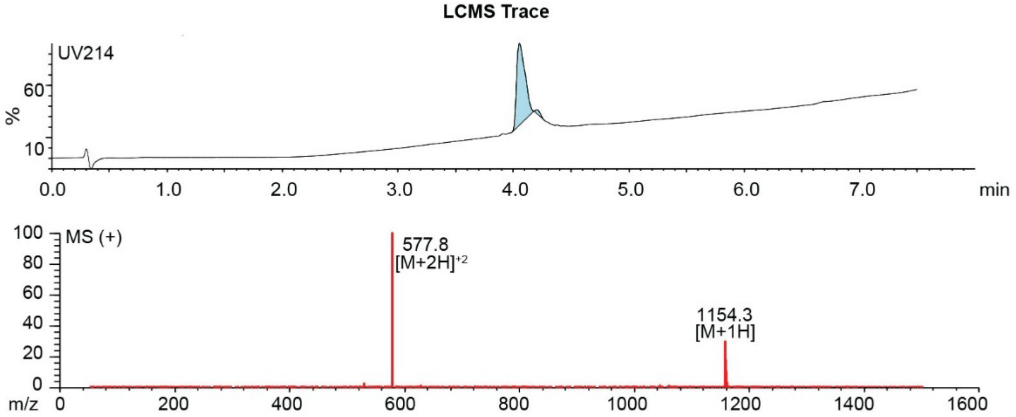

**Figure S2.52.** Synthesis summary of ACLPFYSRC-**DKL** (**26b**).
Expected: [M+1] = 1154.50, $[M+2]^{\div 2}$ = 577.75. Observed: [M+1] = 1154.3, $[M+2]^{\div 2}$ = 577.8.

#### ACLPFYSRC-PPA

Starting material mass: 3.0 mg
Final product mass: 1.7 mg

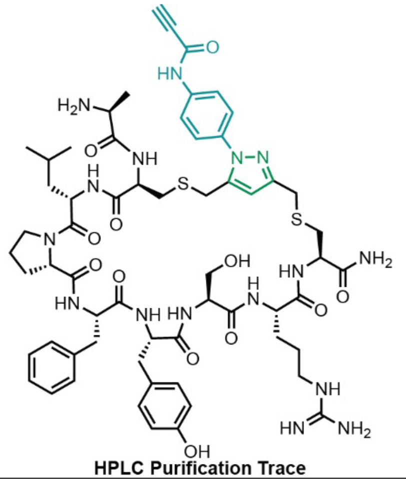

Chemical Formula: $C_{61}H_{80}N_{16}O_{12}S_2$
Exact Mass: 1292.56
Molecular Weight: 1293.53

**HPLC Purification Trace**

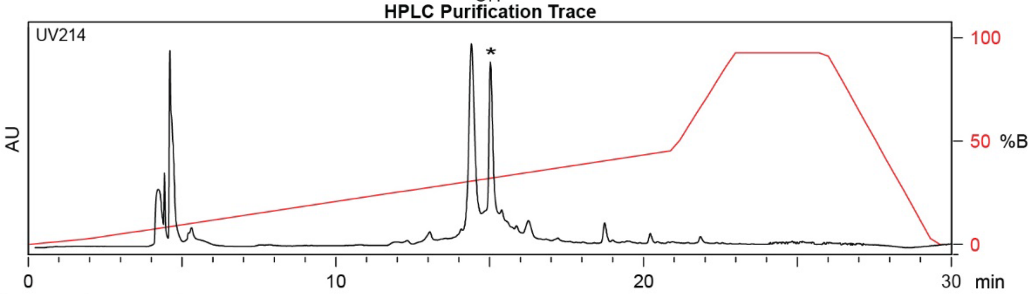

| Time (min) | Solvent B (%) |
|---|---|
| 0 | 2 |
| 2 | 2 |
| 21 | 50 |
| 24 | 100 |
| 27 | 100 |
| 29.5 | 2 |
| 30 | 2 |

Flow Rate: 13 mL / min
Phenomenex Kinetex EVO C18 Prep Column
(100 Å, 5 µm, 21.5 mm X 250 mm)

Solvent A: $H_2O$ + 0.1% (v/v) TFA
Solvent B: MeCN + 0.1% (v/v) TFA

**LCMS Trace**

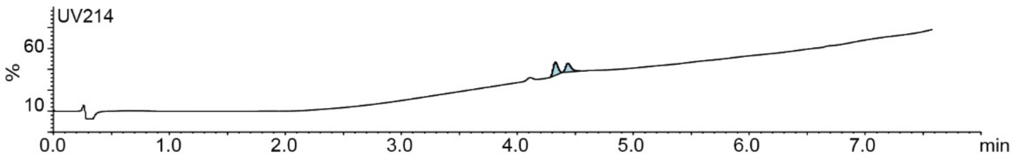

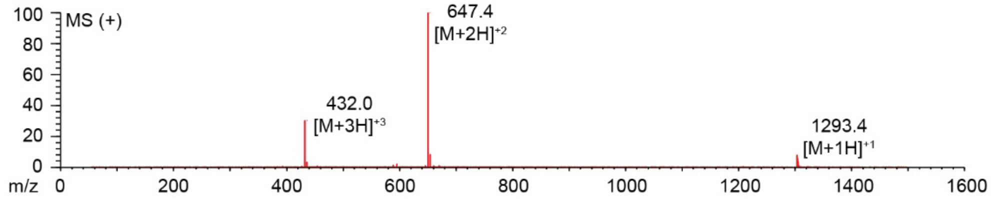

**Figure S2.53.** Synthesis summary of ACLPFYSRC-**PPA** (**26c**).
Expected: [M+1] = 1293.56, $[M+2]^{+2}$ = 647.28, $[M+3]^{+3}$ = 431.85. Observed: [M+1] = 1293.4, $[M+2]^{+2}$ = 647.4, $[M+3]^{+3}$ = 432.0. Note: Two regioisomeric forms of the pyrazole are possible; the structure shown represents one isomer.

#### ACLPFYSRC-PEA

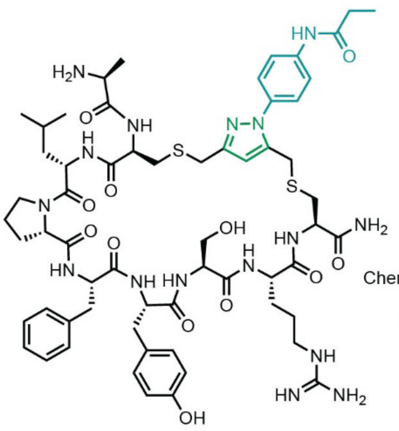

Starting material mass: 3.0 mg
Final product mass: 1.5 mg

Chemical Formula: $C_{61}H_{84}N_{16}O_{12}S_2$
Exact Mass: 1296.59
Molecular Weight: 1297.56

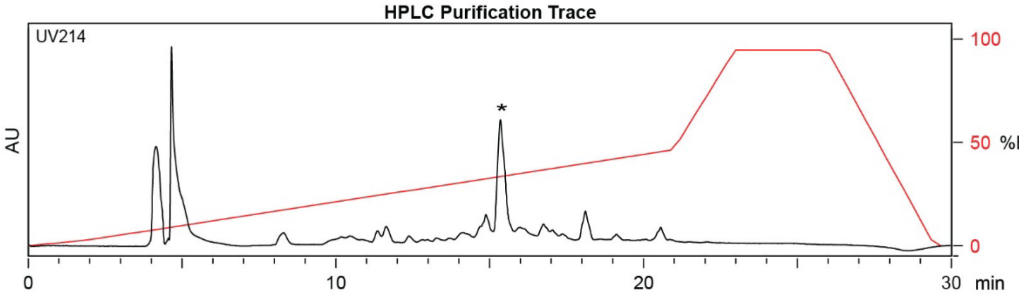

| Time (min) | Solvent B (%) |
|---|---|
| 0 | 2 |
| 2 | 2 |
| 21 | 50 |
| 24 | 100 |
| 27 | 100 |
| 29.5 | 2 |
| 30 | 2 |

Flow Rate: 13 mL / min
Phenomenex Kinetex EVO C18 Prep Column
(100 Å, 5 μm, 21.5 mm X 250 mm)

Solvent A: $H_2O$ + 0.1% (v/v) TFA
Solvent B: MeCN + 0.1% (v/v) TFA

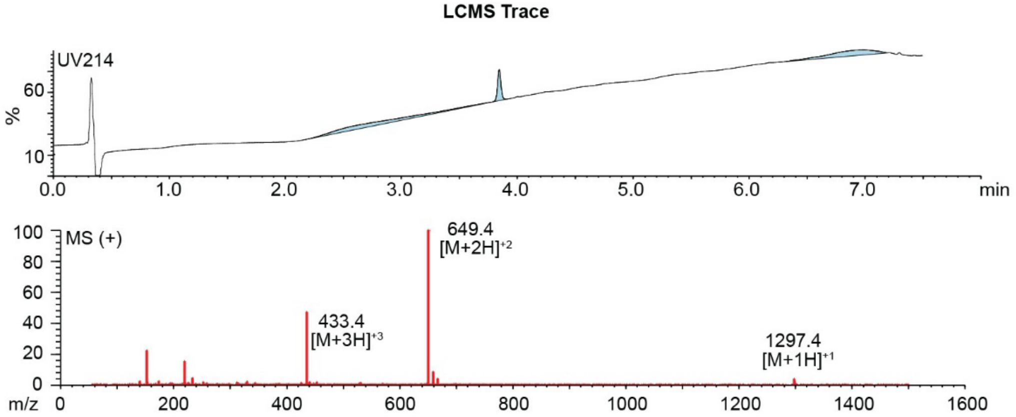

**Figure S2.54.** Synthesis summary of ACLPFYSRC-**PEA** (**26d**).
Expected: [M+1] = 1297.59, $[M+2]^{+2}$ = 649.30, $[M+3]^{+3}$ = 433.20. Observed: [M+1] = 1297.4, $[M+2]^{+2}$ = 649.4, $[M+3]^{+3}$ = 433.4. Note: Two regioisomeric forms of the pyrazole are possible; the structure shown represents one isomer.

#### ACFFYSKNC

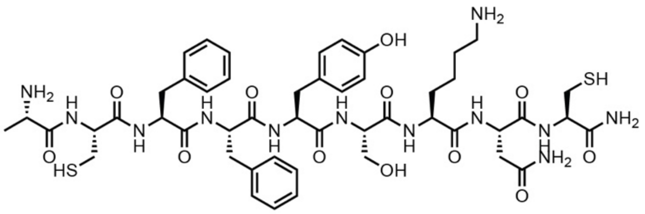

Chemical Formula: $C_{49}H_{68}N_{12}O_{12}S_2$
Exact Mass: 1080.45
Molecular Weight: 1081.28

Starting material mass: N/A mg
Final product mass: 87.3 mg

**Combiflash Purification Trace**

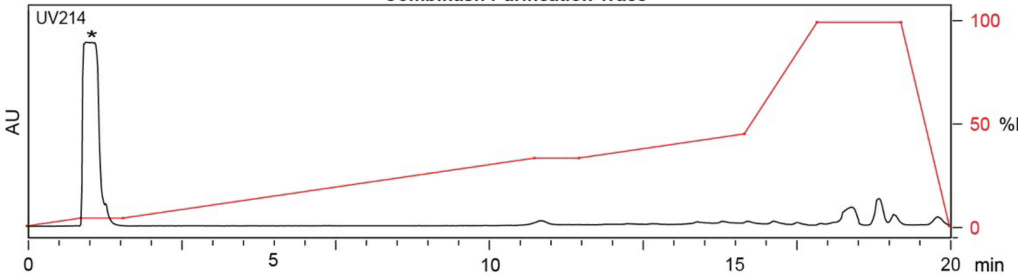

| Time (min) | Solvent B (%) |
|---|---|
| 0 | 0 |
| 1 | 2 |
| 5 | 20 |
| 10 | 40 |
| 15 | 50 |
| 16 | 100 |
| 20 | 0 |

Flow Rate: 35 mL / min
Redisep Rf Gold C18 Reverse Phase Column 30g
(100 Å, 20-40 µm, 300 ± 50 $m^2$/g)

Solvent A: $H_2O$ + 0.1% (v/v) TFA
Solvent B: MeCN + 0.1% (v/v) TFA

**LCMS Trace**

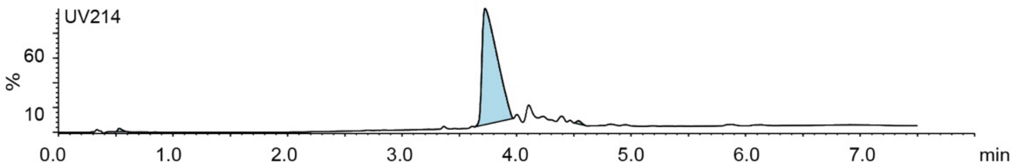

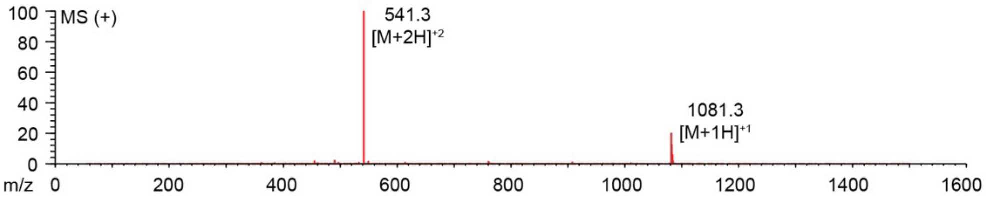

**Figure S2.55.** Synthesis summary of ACFFYSKNC (**27a**).
Expected: [M+1] = 1081.45, $[M+2]^{+2}$ = 541.22. Observed: [M+1] = 1061.3, $[M+2]^{+2}$ = 541.3.

#### ACFFYSKNC-DKL

Starting material mass: 5.4 mg
Final product mass: 0.5 mg

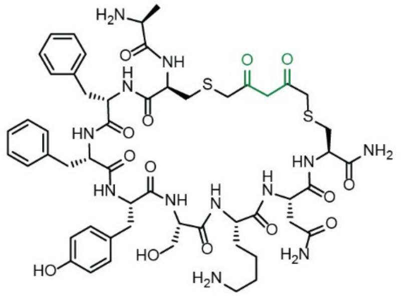

Chemical Formula: $C_{54}H_{72}N_{12}O_{14}S_2$
Exact Mass: 1176.4732
Molecular Weight: 1177.3600

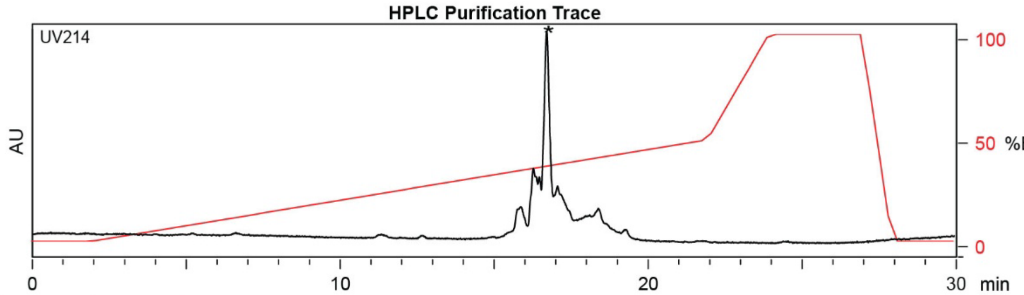

| Time (min) | Solvent B (%) |
|---|---|
| 0 | 2 |
| 2 | 2 |
| 22 | 50 |
| 24 | 100 |
| 27 | 100 |
| 28 | 2 |
| 30 | 2 |

Flow Rate: 13 mL / min
Phenomenex Kinetex EVO C18 Prep Column
(100 Å, 5 µm, 21.5 mm X 250 mm)

Solvent A: $H_2O$ + 0.1% (v/v) TFA
Solvent B: MeCN + 0.1% (v/v) TFA

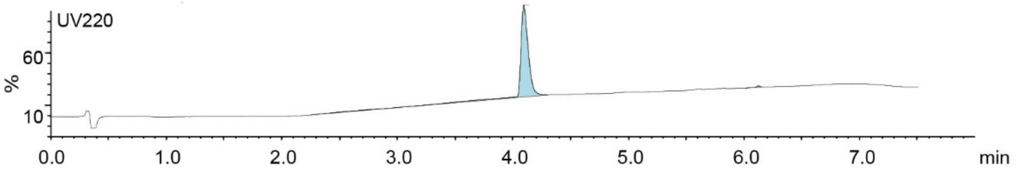

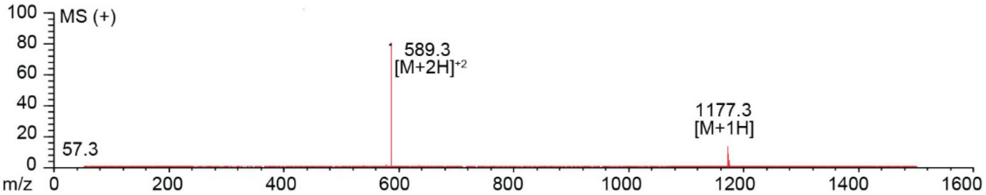

**Figure S2.56.** Synthesis summary of ACFFYSKNC-**DKL** (**27b**).
Expected: [M+1] = 1177.4732, $[M+2]^{\div 2}$ = 589.2366. Observed: [M+1] = 1177.3, $[M+2]^{\div 2}$ = 589.3.

#### ACFFYSKNC-PPA

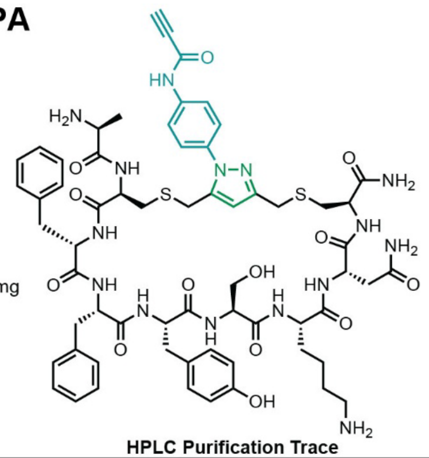

Starting material mass: 3.6 mg
Final product mass: 1.3 mg

Chemical Formula: $C_{63}H_{77}N_{15}O_{13}S_2$
Exact Mass: 1315.5267
Molecular Weight: 1316.5210

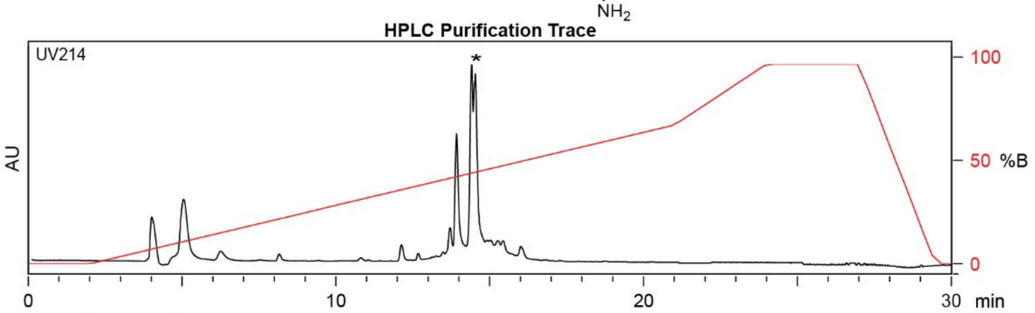

| Time (min) | Solvent B (%) |
|---|---|
| 0 | 2 |
| 2 | 2 |
| 21 | 50 |
| 24 | 100 |
| 27 | 100 |
| 29.5 | 2 |
| 30 | 2 |

Flow Rate: 13 mL / min
Phenomenex Kinetex EVO C18 Prep Column
(100 Å, 5 μm, 21.5 mm X 250 mm)

Solvent A: $H_2O$ + 0.1% (v/v) TFA
Solvent B: MeCN + 0.1% (v/v) TFA

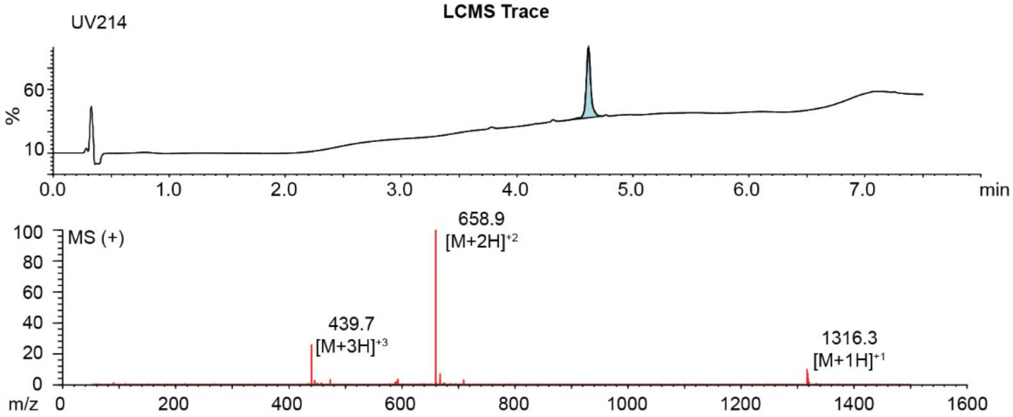

**Figure S2.57.** Synthesis summary of ACFFYSKNC-**PPA** (**27c**).
Expected: [M+1] = 1316.5267, $[M+2]^{+2}$ = 658.7634, $[M+3]^{+3}$ = 439.5089. Observed: [M+1] = 1316.3, $[M+2]^{+2}$ = 658.9, $[M+3]^{+3}$ = 439.7. Note: Two regioisomeric forms of the pyrazole are possible; the structure shown represents one isomer.

#### ACFFYSKNC-PEA

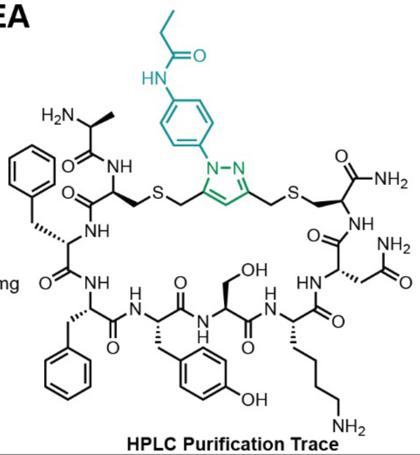

Starting material mass: 3.1 mg
Final product mass: 1.2 mg

Chemical Formula: $C_{63}H_{81}N_{15}O_{13}S_2$
Exact Mass: 1319.56
Molecular Weight: 1320.55

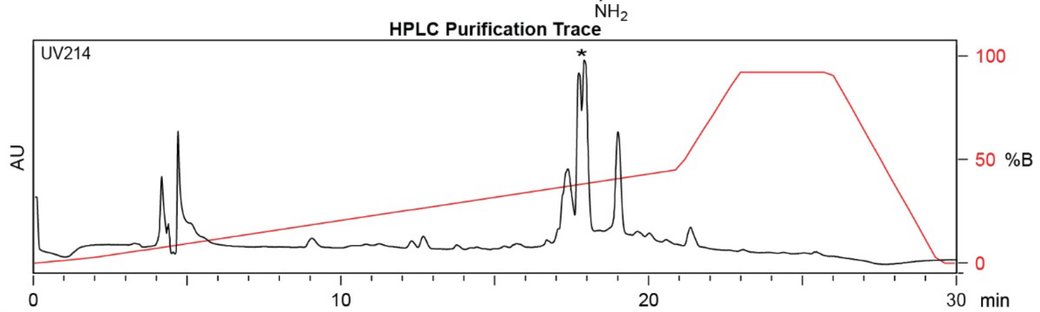

| Time (min) | Solvent B (%) |
|---|---|
| 0 | 2 |
| 2 | 2 |
| 21 | 50 |
| 24 | 100 |
| 27 | 100 |
| 29.5 | 2 |
| 30 | 2 |

Flow Rate: 13 mL / min
Phenomenex Kinetex EVO C18 Prep Column
(100 Å, 5 µm, 21.5 mm X 250 mm)

Solvent A: $H_2O$ + 0.1% (v/v) TFA
Solvent B: MeCN + 0.1% (v/v) TFA

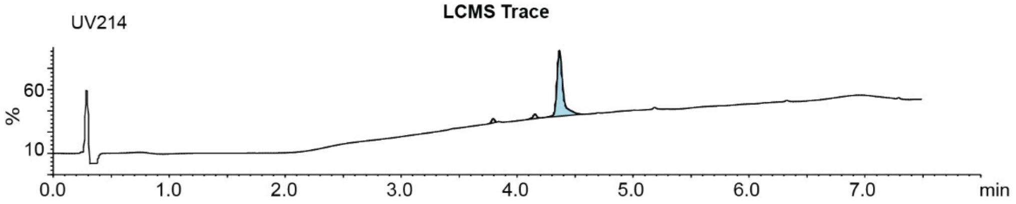

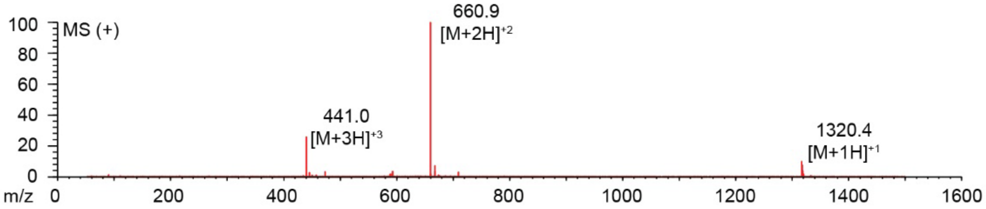

**Figure S2.58.** Synthesis summary of ACFFYSKNC-**PEA** (**27d**).
Expected: [M+1] = 1320.56, $[M+2]^{+2}$ = 660.78, $[M+3]^{+3}$ = 440.85. Observed: [M+1] = 1320.4, $[M+2]^{+2}$ = 660.9, $[M+3]^{+3}$ = 441.0. Note: Two regioisomeric forms of the pyrazole are possible; the structure shown represents one isomer.

#### ACTDRFFKC-DKL

Starting material mass:
Final product mass:

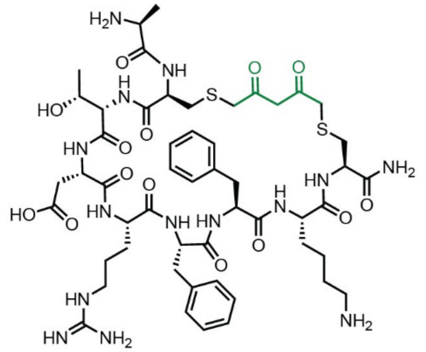

Chemical Formula: $C_{52}H_{76}N_{14}O_{14}S_2$
Exact Mass: 1184.51
Molecular Weight: 1185.38

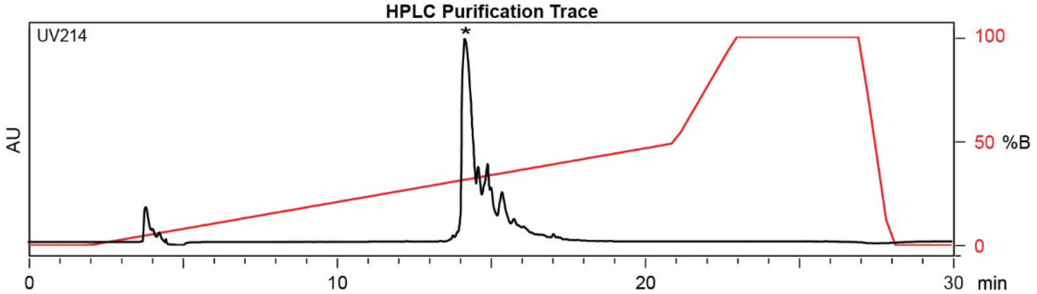

| Time (min) | Solvent B (%) |
|---|---|
| 0 | 2 |
| 2 | 2 |
| 22 | 50 |
| 24 | 100 |
| 27 | 100 |
| 28 | 2 |
| 30 | 2 |

Flow Rate: 13 mL / min
Phenomenex Kinetex EVO C18 Prep Column
(100 Å, 5 µm, 21.5 mm X 250 mm)

Solvent A: $H_2O$ + 0.1% (v/v) TFA
Solvent B: MeCN + 0.1% (v/v) TFA

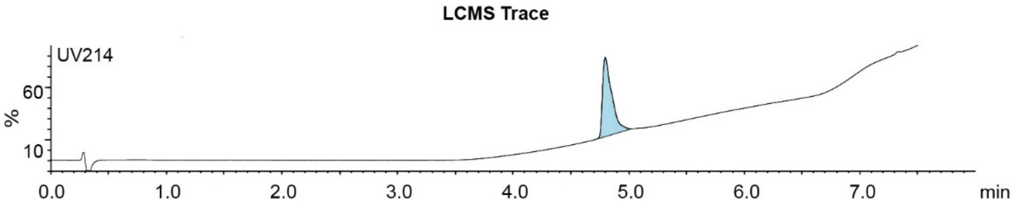

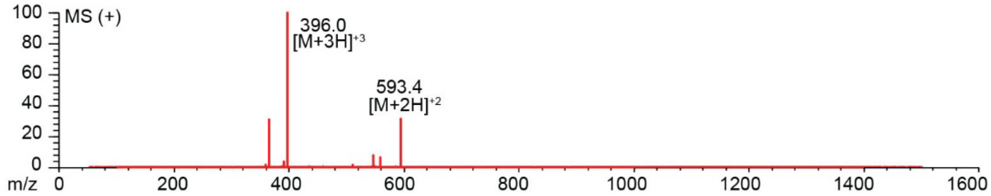

**Figure S2.59.** Synthesis summary of ACTDRFFKC-**DKL** (**28b**).
Expected: [M+1] = 1185.51, $[M+2]^{+2}$ = 593.26, $[M+3]^{+3}$ = 395.84. Observed: $[M+2]^{+2}$ = 593.4, $[M+3]^{+3}$ = 396.0.

#### ACTDRFFKC-PPA

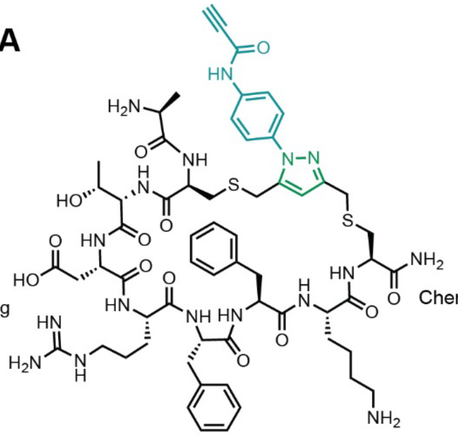

Starting material mass: 5.3 mg
Final product mass: 2.2 mg

Chemical Formula: $C_{61}H_{81}N_{17}O_{13}S_2$
Exact Mass: 1323.56
Molecular Weight: 1324.55

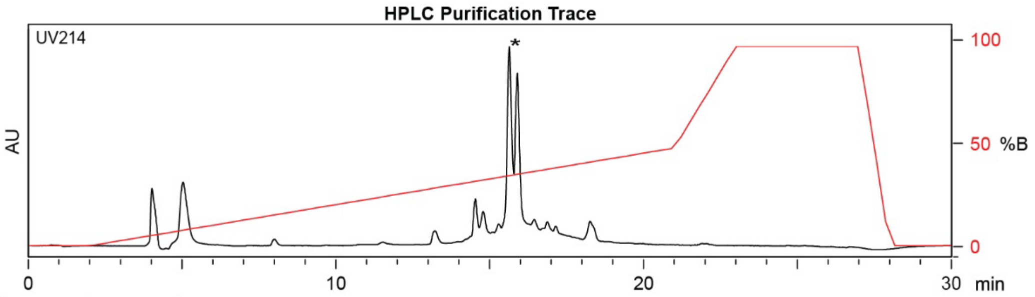

| Time (min) | Solvent B (%) |
|---|---|
| 0 | 2 |
| 2 | 2 |
| 21 | 50 |
| 24 | 100 |
| 27 | 100 |
| 29.5 | 2 |
| 30 | 2 |

Flow Rate: 13 mL / min
Phenomenex Kinetex EVO C18 Prep Column
(100 Å, 5 µm, 21.5 mm X 250 mm)

Solvent A: $H_2O$ + 0.1% (v/v) TFA
Solvent B: MeCN + 0.1% (v/v) TFA

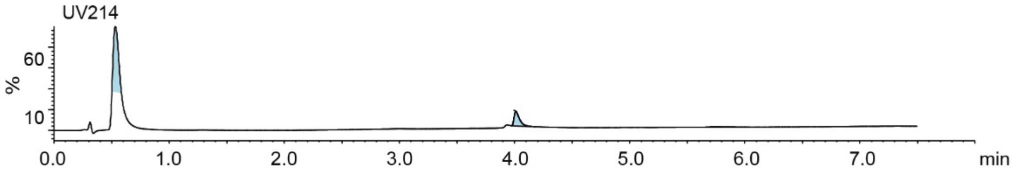

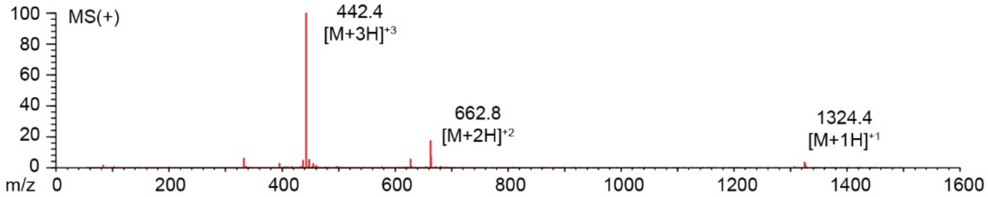

**Figure S2.60.** Synthesis summary of ACTDRFFKC-**PPA** (**28c**).
Expected: [M+1] = 1324.56, $[M+2]^{\div 2}$ = 662.78, $[M+3]^{\div 3}$ = 442.19. Observed: [M+1] = 1324.4, $[M+2]^{\div 2}$ = 662.8, $[M+3]^{\div 3}$ = 442.4. Note: Two regioisomeric forms of the pyrazole are possible; the structure shown represents one isomer.

### ACFHSSFFC-DKL

Starting material mass:
Final product mass:

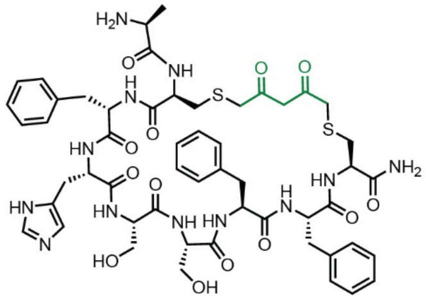

Chemical Formula: $C_{53}H_{66}N_{12}O_{13}S_2$
Exact Mass: 1142.4314
Molecular Weight: 1143.3020

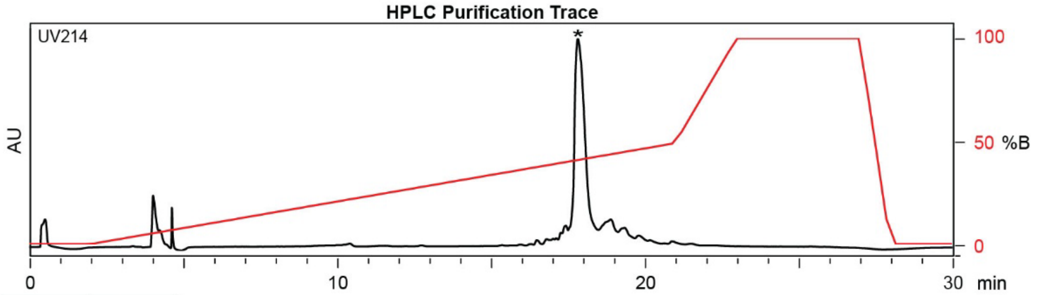

| Time (min) | Solvent B (%) |
|---|---|
| 0 | 2 |
| 2 | 2 |
| 22 | 50 |
| 24 | 100 |
| 27 | 100 |
| 28 | 2 |
| 30 | 2 |

Flow Rate: 13 mL / min
Phenomenex Kinetex EVO C18 Prep Column
(100 Å, 5 µm, 21.5 mm X 250 mm)

Solvent A: $H_2O$ + 0.1% (v/v) TFA
Solvent B: MeCN + 0.1% (v/v) TFA

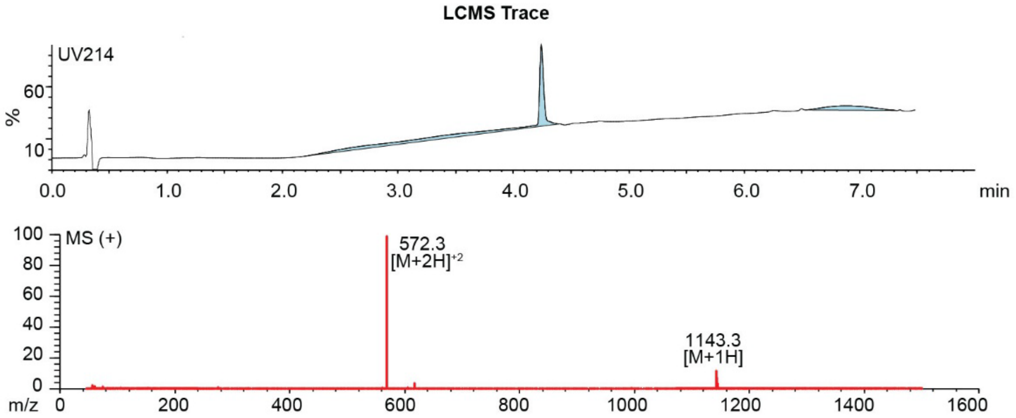

**Figure S2.61.** Synthesis summary of ACFHSSFFC-**DKL** (**29b**).
Expected: [M+1] = 1143.4314, $[M+2]^{+2}$ = 572.2157. Observed: [M+1] = 1143.3, $[M+2]^{+2}$ = 572.3.

#### ACTDRFFKC-PPA

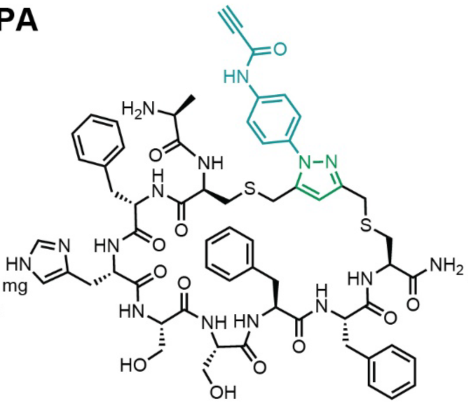

Starting material mass: 2.9 mg
Final product mass: 1.0 mg

Chemical Formula: $C_{62}H_{71}N_{15}O_{12}S_2$
Exact Mass: 1281.48
Molecular Weight: 1282.46

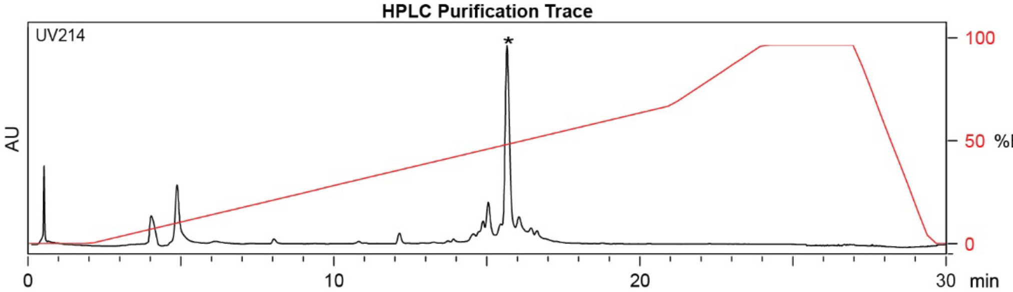

| Time (min) | Solvent B (%) |
|---|---|
| 0 | 2 |
| 2 | 2 |
| 21 | 70 |
| 24 | 100 |
| 27 | 100 |
| 29.5 | 2 |
| 30 | 2 |

Flow Rate: 13 mL / min
Phenomenex Kinetex EVO C18 Prep Column
(100 Å, 5 µm, 21.5 mm X 250 mm)

Solvent A: $H_2O$ + 0.1% (v/v) TFA
Solvent B: MeCN + 0.1% (v/v) TFA

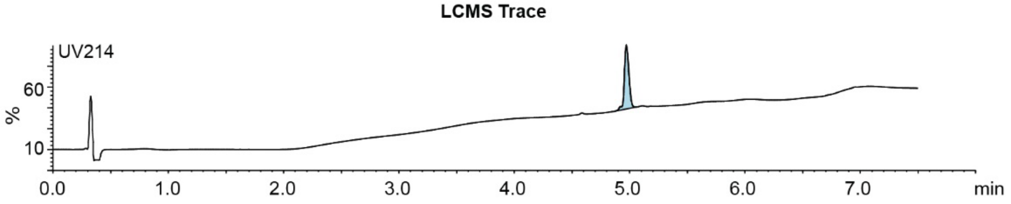

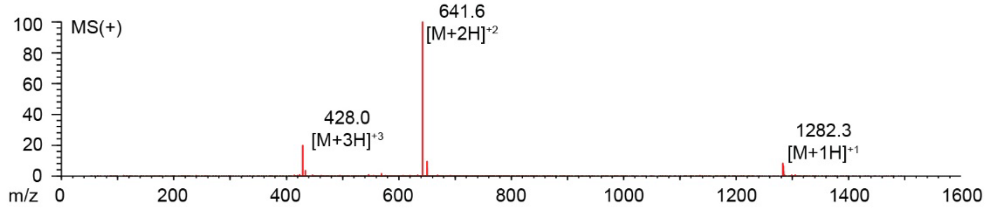

**Figure S2.62.** Synthesis summary of ACFHSSFFC-**PPA** (**29c**).
Expected: [M+1] = 1282.48, $[M+2]^{+2}$ = 641.74, $[M+3]^{+3}$ = 428.16. Observed: [M+1] = 1282.3, $[M+2]^{+2}$ = 641.6, $[M+3]^{+3}$ = 428.0. Note: Two regioisomeric forms of the pyrazole are possible; the structure shown represents one isomer.

#### RCGIGCV

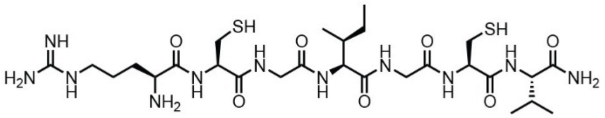

Chemical Formula: $C_{27}H_{51}N_{11}O_7S_2$
Exact Mass: 705.3414
Molecular Weight: 705.8950

Starting material mass: N/A mg
Final product mass: 193 mg

**Combiflash Purification Trace**

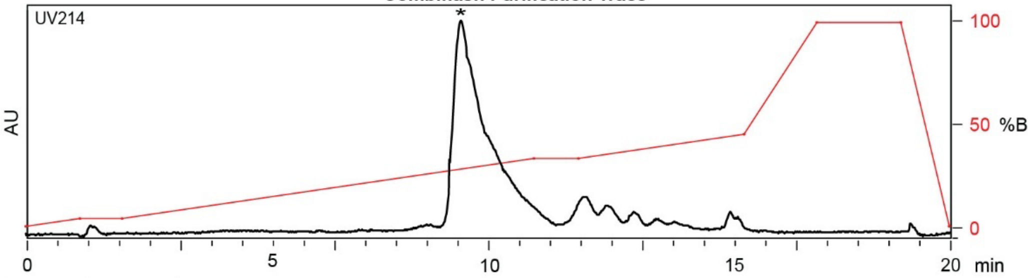

| Time (min) | Solvent B (%) |
|---|---|
| 0 | 0 |
| 1 | 2 |
| 5 | 20 |
| 10 | 40 |
| 15 | 50 |
| 16 | 100 |
| 20 | 0 |

Flow Rate: 35 mL / min
Redisep Rf Gold C18 Reverse Phase Column 30g
(100 Å, 20-40 μm, 300 ± 50 m²/g)

Solvent A: $H_2O$ + 0.1% (v/v) TFA
Solvent B: MeCN + 0.1% (v/v) TFA

**LCMS Trace**

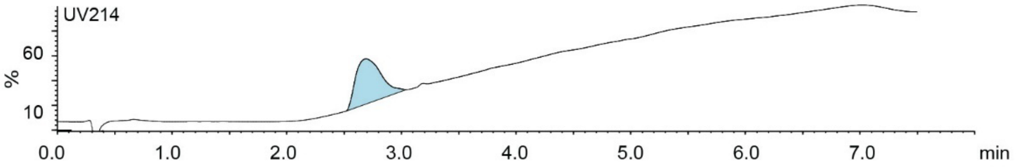

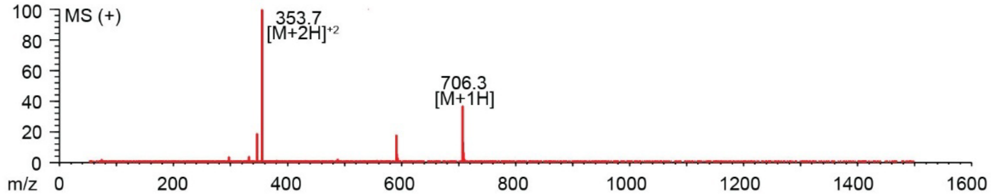

**Figure S2.63.** Synthesis summary of RCGIGCV (**30a**).
Expected: [M+1] = 706.3414, $[M+2]^{÷2}$ = 353.6707. Observed: [M+1] = 706.3, $[M+2]^{÷2}$ = 353.7.

#### ACFFYSKNCGGG(Z)-DKL

Starting material mass: 7.3 mg
Final product mass: 3.0 mg

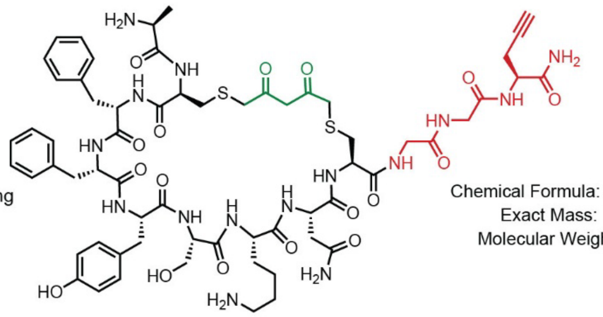

Chemical Formula: $C_{63}H_{83}N_{15}O_{17}S_2$
Exact Mass: 1385.5533
Molecular Weight: 1386.5650

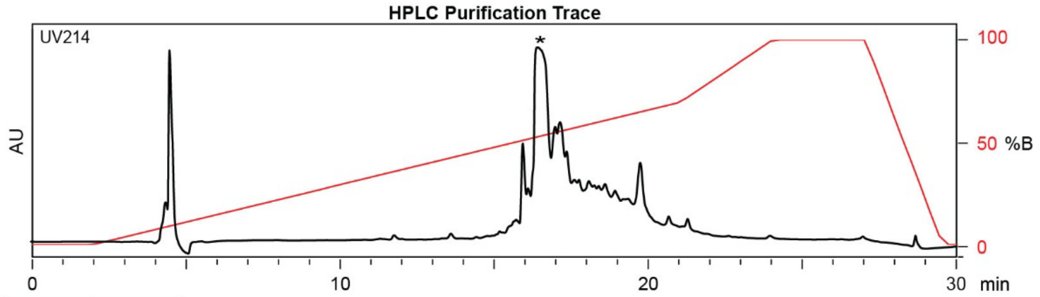

| Time (min) | Solvent B (%) |
|---|---|
| 0 | 2 |
| 2 | 2 |
| 22 | 50 |
| 24 | 100 |
| 27 | 100 |
| 28 | 2 |
| 30 | 2 |

Flow Rate: 13 mL / min
Phenomenex Kinetex EVO C18 Prep Column
(100 Å, 5 µm, 21.5 mm X 250 mm)

Solvent A: $H_2O$ + 0.1% (v/v) TFA
Solvent B: MeCN + 0.1% (v/v) TFA

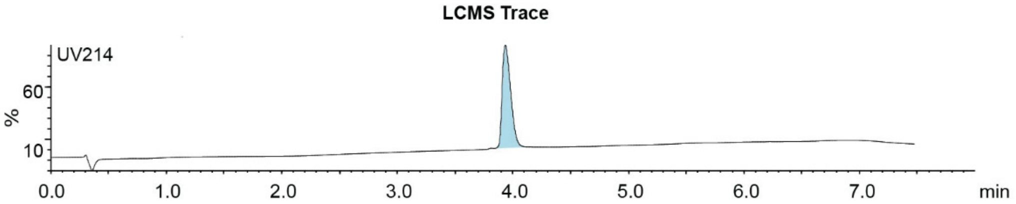

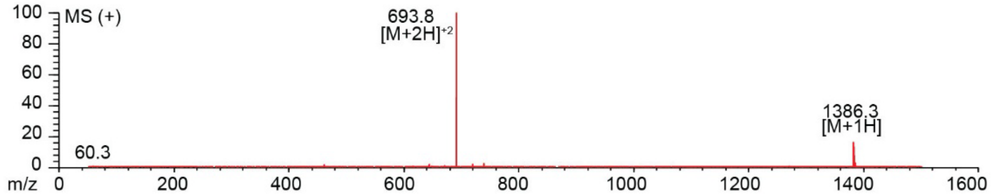

**Figure S2.64.** Synthesis summary of ACFFYSKNCGGG(Z)-**DKL** (**31b**).
Expected: [M+1] = 1386.5533, $[M+2]^{\div 2}$ = 693.7766. Observed: [M+1] = 1386.3, $[M+2]^{\div 2}$ = 693.8.

#### ACFFYSKNCGGG(Z)-PPA

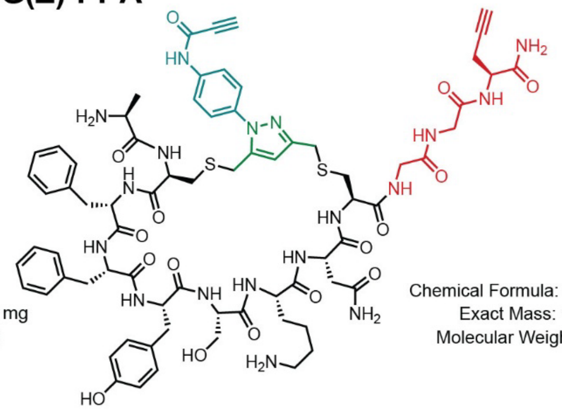

Starting material mass: 3.0 mg
Final product mass: 1.8 mg

Chemical Formula: $C_{72}H_{88}N_{18}O_{16}S_2$
Exact Mass: 1524.6067
Molecular Weight: 1525.7260

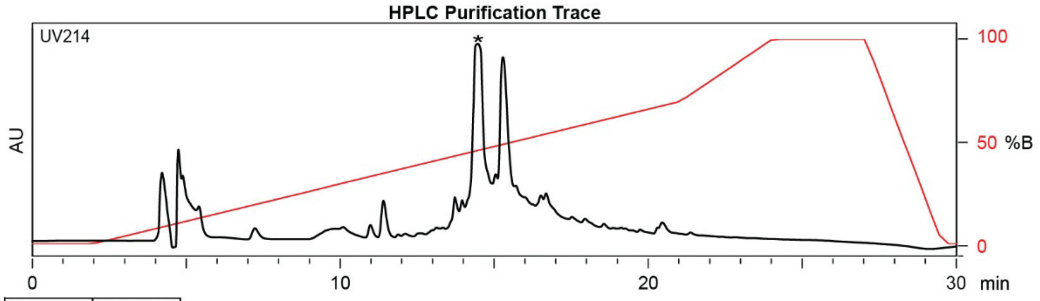

| Time (min) | Solvent B (%) |
|---|---|
| 0 | 2 |
| 2 | 2 |
| 21 | 70 |
| 24 | 100 |
| 27 | 100 |
| 29.5 | 2 |
| 30 | 2 |

Flow Rate: 13 mL / min
Phenomenex Kinetex EVO C18 Prep Column
(100 Å, 5 µm, 21.5 mm X 250 mm)

Solvent A: $H_2O$ + 0.1% (v/v) TFA
Solvent B: MeCN + 0.1% (v/v) TFA

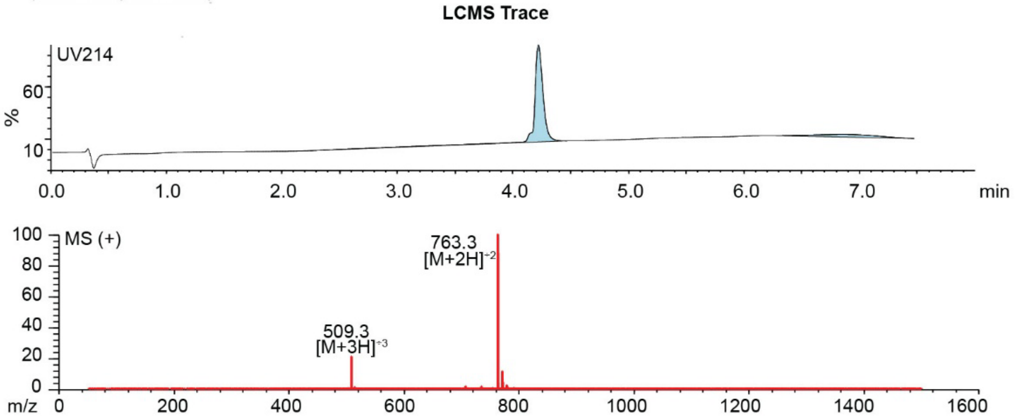

**Figure S2.65.** Synthesis summary of ACFFYSKNCGGG(Z)-**PPA** (**31c**).
Expected: [M+1] = 1525.6067, $[M+2]^{\div 2}$ = 763.3034, $[M+3]^{\div 3}$ = 509.2022. Observed: $[M+2]^{\div 2}$ = 763.3, $[M+3]^{\div 3}$ = 509.3. Note: Two regioisomeric forms of the pyrazole are possible; the structure shown represents one isomer.

#### ACFFASKNC-DKL

Starting material mass: 7.0 mg
Final product mass: 3.8 mg

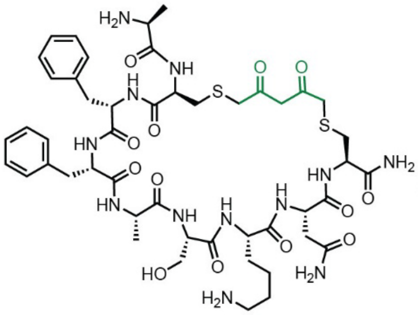

Chemical Formula: $C_{48}H_{68}N_{12}O_{13}S_2$
Exact Mass: 1084.4470
Molecular Weight: 1085.2630

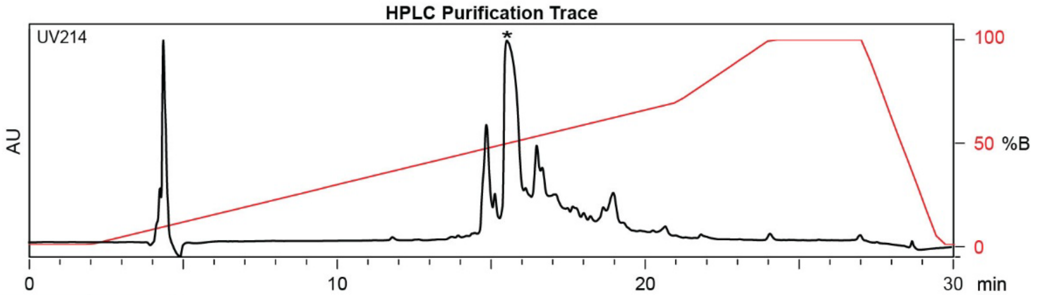

| Time (min) | Solvent B (%) |
|---|---|
| 0 | 2 |
| 2 | 2 |
| 22 | 50 |
| 24 | 100 |
| 27 | 100 |
| 28 | 2 |
| 30 | 2 |

Flow Rate: 13 mL / min
Phenomenex Kinetex EVO C18 Prep Column
(100 Å, 5 µm, 21.5 mm X 250 mm)

Solvent A: $H_2O$ + 0.1% (v/v) TFA
Solvent B: MeCN + 0.1% (v/v) TFA

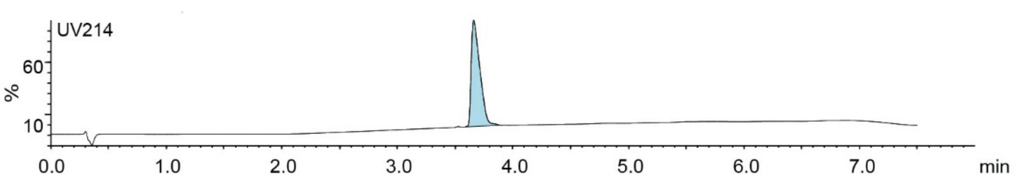

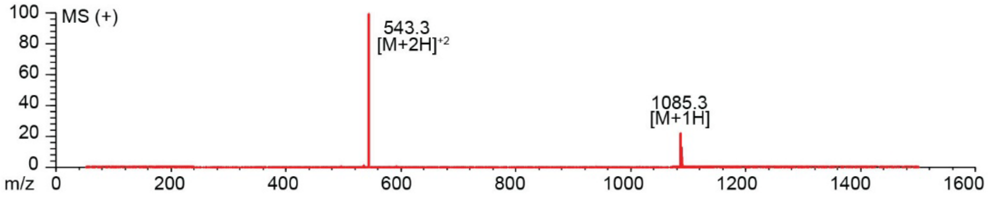

**Figure S2.66.** Synthesis summary of ACFFASKNC-**DKL** (**32b**).
Expected: [M+1] = 1085.4470, $[M+2]^{+2}$ = 543.2235. Observed: [M+1] = 1085.3, $[M+2]^{+2}$ = 543.3.

### ACFFASKNC-PPA

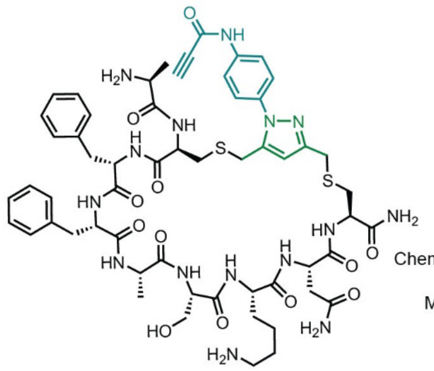

Starting material mass: 3.8 mg
Final product mass: 2.0 mg

Chemical Formula: $C_{57}H_{73}N_{15}O_{12}S_2$
Exact Mass: 1223.5005
Molecular Weight: 1224.4240

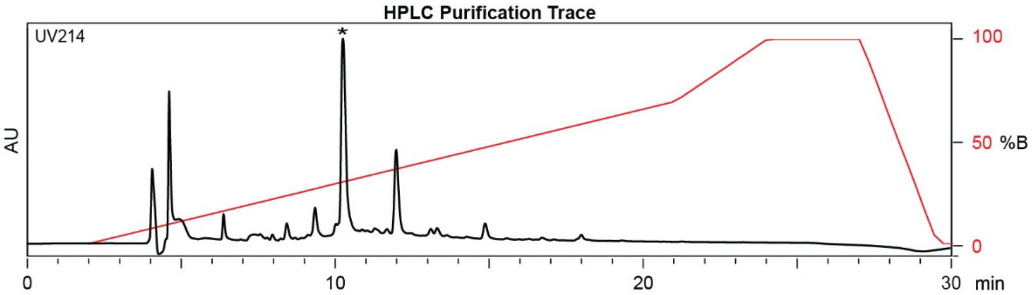

| Time (min) | Solvent B (%) |
|---|---|
| 0 | 2 |
| 2 | 2 |
| 21 | 70 |
| 24 | 100 |
| 27 | 100 |
| 29.5 | 2 |
| 30 | 2 |

Flow Rate: 13 mL / min
Phenomenex Kinetex EVO C18 Prep Column
(100 Å, 5 µm, 21.5 mm X 250 mm)

Solvent A: $H_2O$ + 0.1% (v/v) TFA
Solvent B: MeCN + 0.1% (v/v) TFA

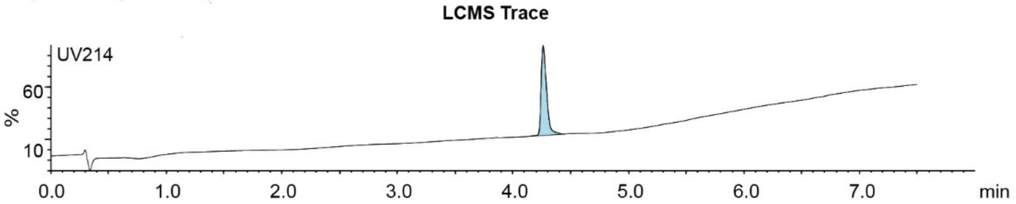

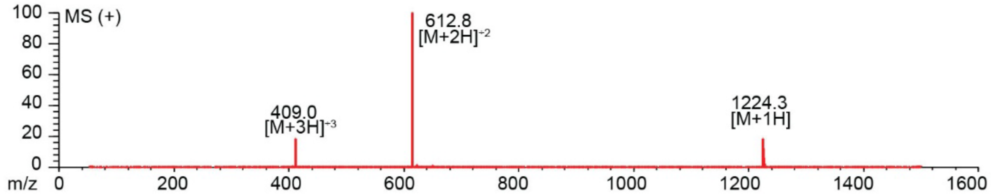

**Figure S2.67.** Synthesis summary of ACFFASKNC-**PPA** (**32c**).
Expected: [M+1] = 1224.5005, $[M+2]^{+2}$ = 612.7502, $[M+3]^{+3}$ = 408.8335. Observed: [M+1] = 1224.3, $[M+2]^{+2}$ = 612.8, $[M+3]^{+3}$ = 409.0. Note: Two regioisomeric forms of the pyrazole are possible; the structure shown represents one isomer.

#### ACFFASKNCGGG(Z)-DKL

Starting material mass: 8.0 mg
Final product mass: 2.9 mg

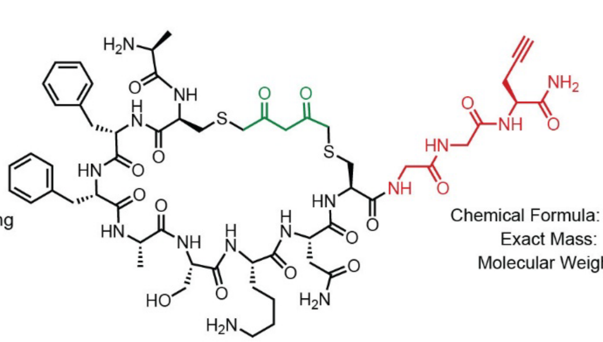

Chemical Formula: $C_{57}H_{79}N_{15}O_{16}S_2$
Exact Mass: 1293.5271
Molecular Weight: 1294.4680

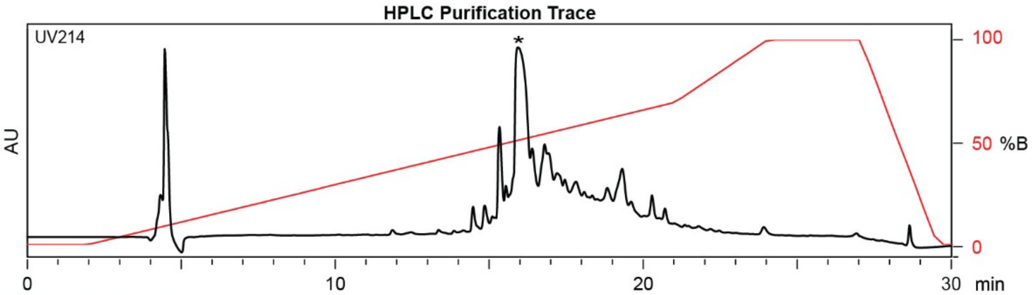

| Time (min) | Solvent B (%) |
|---|---|
| 0 | 2 |
| 2 | 2 |
| 22 | 50 |
| 24 | 100 |
| 27 | 100 |
| 28 | 2 |
| 30 | 2 |

Flow Rate: 13 mL / min
Phenomenex Kinetex EVO C18 Prep Column
(100 Å, 5 µm, 21.5 mm X 250 mm)

Solvent A: $H_2O$ + 0.1% (v/v) TFA
Solvent B: MeCN + 0.1% (v/v) TFA

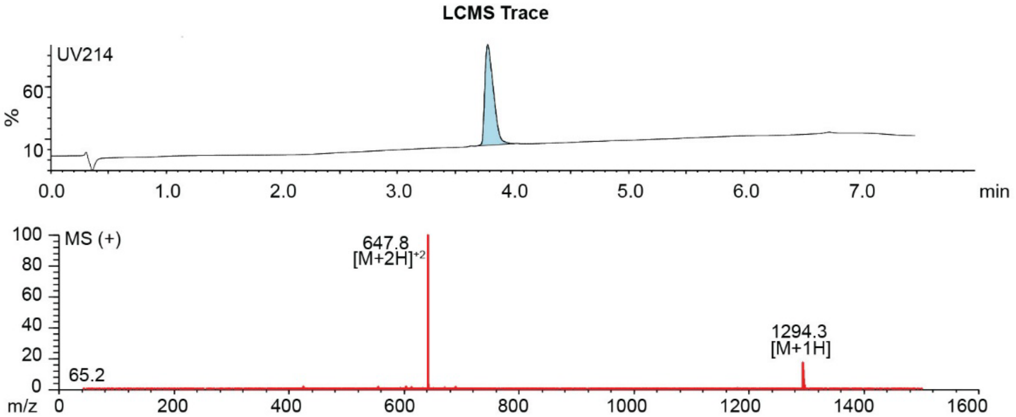

**Figure S2.68.** Synthesis summary of ACFFASKNCGGG(Z)-**DKL** (**33b**).
Expected: [M+1] = 1294.5271, $[M+2]^{+2}$ = 647.7636. Observed: [M+1] = 1294.3, $[M+2]^{+2}$ = 647.8.

#### ACFFASKNCGGG(Z)-PPA

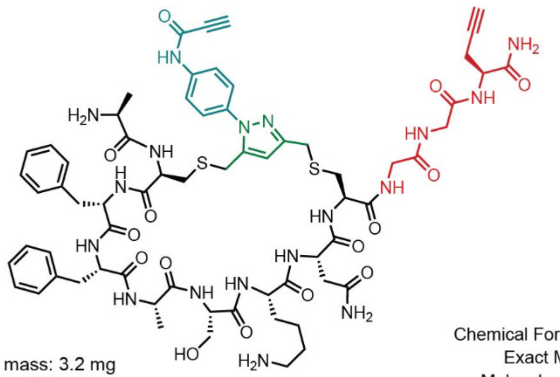

Chemical Formula: $C_{66}H_{84}N_{18}O_{15}S_2$
Exact Mass: 1432.5805
Molecular Weight: 1433.6290

Starting material mass: 3.2 mg
Final product mass: 2.2 mg

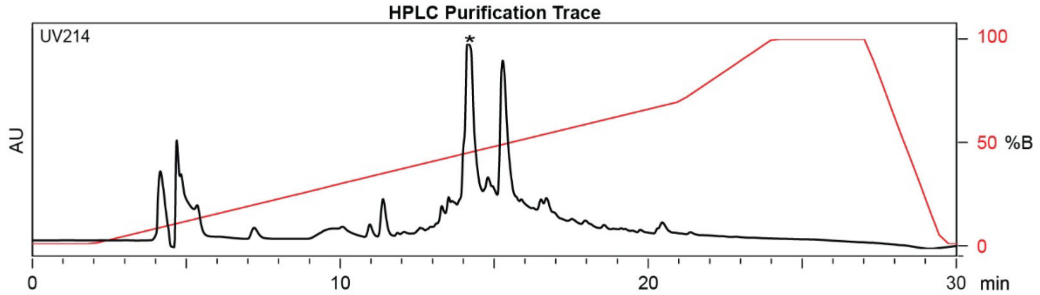

| Time (min) | Solvent B (%) |
|---|---|
| 0 | 2 |
| 2 | 2 |
| 21 | 70 |
| 24 | 100 |
| 27 | 100 |
| 29.5 | 2 |
| 30 | 2 |

Flow Rate: 13 mL / min
Phenomenex Kinetex EVO C18 Prep Column
(100 Å, 5 µm, 21.5 mm X 250 mm)

Solvent A: $H_2O$ + 0.1% (v/v) TFA
Solvent B: MeCN + 0.1% (v/v) TFA

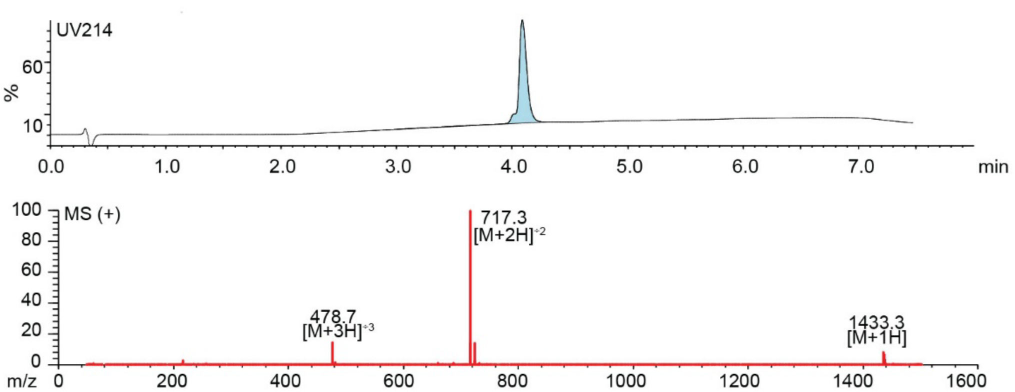

**Figure S2.69.** Synthesis summary of ACFFASKNCGGG(Z)-**PPA** (**33c**).
Expected: [M+1] = 1433.5805, $[M+2]^{+2}$ = 717.2902, $[M+3]^{+3}$ = 478.5268. Observed: [M+1] = 1433.3, $[M+2]^{+2}$ = 717.3, $[M+3]^{+3}$ = 478.7. Note: Two regioisomeric forms of the pyrazole are possible; the structure shown represents one isomer.

#### $^1$H NMR Spectra

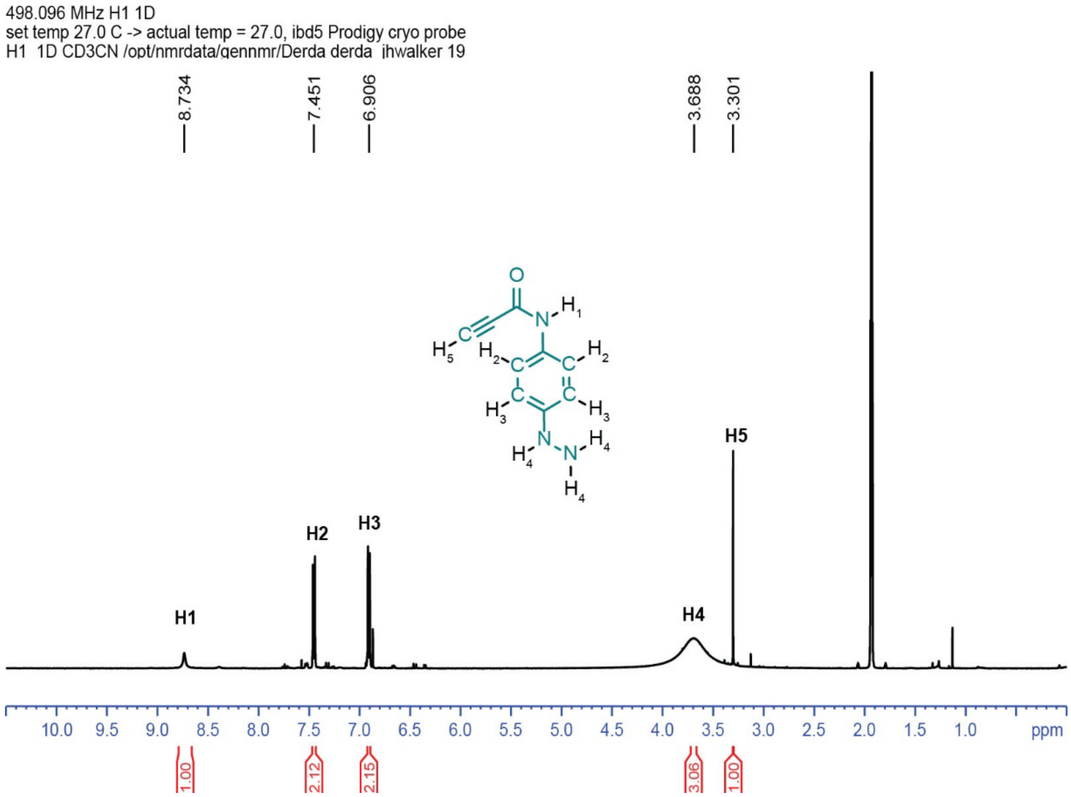

**Figure S2.70.** $^1$H NMR Spectra of *N*-(4-hydrazinylphenyl)propiolamide.
CD3CN: δ 1.93 ppm (qu).

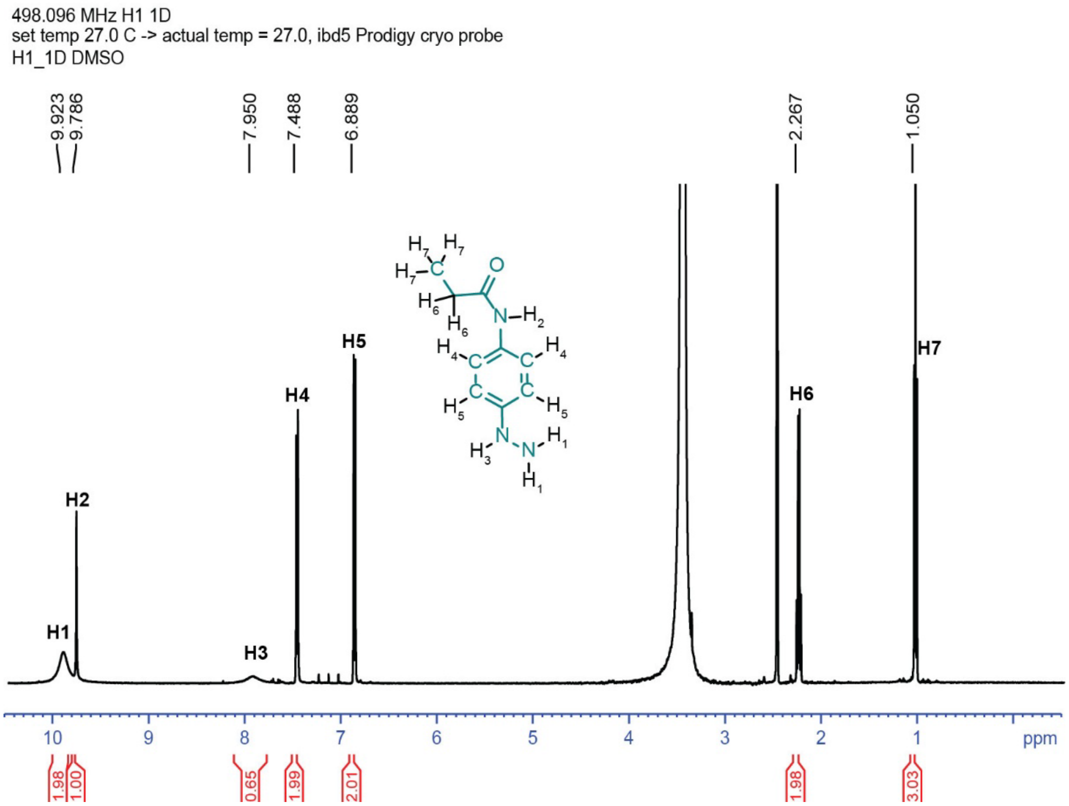

**Figure S2.71.** $^1$H NMR Spectra of *N*-(4-hydrazinylphenyl)propanamide.
